# Interactions of the Protein Tyrosine Phosphatase PTPN3 with Viral and Cellular Partners through its PDZ Domain: Insights into Structural Determinants and Phosphatase Activity

**DOI:** 10.1101/2023.03.23.533931

**Authors:** Mariano Genera, Baptiste Colcombet-Cazenave, Anastasia Croitoru, Bertrand Raynal, Ariel Mechaly, Joël Caillet, Ahmed Haouz, Nicolas Wolff, Célia Caillet-Saguy

## Abstract

The human protein tyrosine phosphatase non-receptor type 3 (PTPN3) is a phosphatase containing a PDZ (PSD-95/Dlg/ZO-1) domain that has been found to play both tumor-suppressive and tumor-promoting roles in various cancers, despite limited knowledge of its cellular partners and signaling functions. Notably, the high-risk genital human papillomavirus (HPV) types 16 and 18 and the hepatitis B virus (HBV) target the PDZ domain of PTPN3 through PDZ-binding motifs (PBMs) in their E6 and HBc proteins respectively.

This study focuses on the interactions between the PTPN3 PDZ domain (PTPN3-PDZ) and PBMs of viral and cellular protein partners. The solved X-ray structures of complexes between PTPN3-PDZ and PBMs of E6 of HPV18 and the tumor necrosis factor-alpha converting enzyme (TACE) reveal two novel interactions. We provide new insights into key structural determinants of PBM recognition by PTPN3 by screening the selectivity of PTPN3-PDZ recognition of PBMs, and by comparing the PDZome binding profiles of PTPN3-recognized PBMs and the interactome of PTPN3-PDZ.

The PDZ domain of PTPN3 was known to auto-inhibit the protein’s phosphatase activity. We discovered that the linker connecting the PDZ and phosphatase domains is involved in this inhibition, and that the binding of PBMs does not impact this catalytic regulation.

Overall, the study sheds light on the interactions and structural determinants of PTPN3 with its cellular and viral partners, as well as on the inhibitory role of its PDZ domain on its phosphatase activity.

## 1 Introduction

Kinase and phosphatase proteins play a major role in cell signaling by regulating the levels of phosphorylated species in signal transduction pathways that control cellular processes such as growth, differentiation, migration, survival, and apoptosis. Large scale genetic analyses of human tumors have highlighted the relevance of protein tyrosine phosphatases either as putative tumor suppressors or as candidate oncoproteins (Julien et al., 2011). Alterations in their expression levels and/or mutations have been suggested to play a role in many cancers (Hendriks and Böhmer, 2016). Examples notably include the promotion of cholangiocarcinoma cell proliferation and migration by gain-of-function mutations or increased expression of the protein tyrosine phosphatase non-receptor type 3 (PTPN3) than in nontumor tissues (Gao et al., 2014).

PTPN3 is a multidomain protein of 913 amino acids comprising a N-terminal FERM domain that determines subcellular localization, a central PDZ domain involved in protein-protein interactions, a short linker of 30-residues and a C-terminal protein tyrosine phosphatase (PTP) domain able to dephosphorylate protein substrates (Figure 1). The linker that connects the FERM and PDZ domains is about 200 amino acids long and predicted mostly unstructured. The FERM-PDZ linker was shown to be cleaved *in vitro* by trypsin, releasing a fragment of around 50 kDa called bidomain that corresponds to the PDZ and PTP domains connected by the 30-residue linker (Zhang et al., 1995). The proteolytic cleavage of the FERM domain of PTPN3 increases its catalytic activity (Zhang et al., 1995). More recently, it has been reported that the PDZ domains of PTPN3 (PTPN3-PDZ) and of its homolog PTPN4 (PTPN4-PDZ) exert an inhibitory effect on the catalytic activity of the adjacent PTP domain (Chen et al., 2014; Maisonneuve et al., 2014). Moreover, we showed that the binding of a PDZ-binding motif (PBM) to the PDZ domain of PTPN4 partially releases this catalytic inhibition (Maisonneuve et al., 2014) and that the linker between the PDZ and the PTP is essential for the regulation by both the PDZ domain and the PBM (Maisonneuve et al., 2016; Caillet-Saguy et al., 2017). Only a handful of PTPN3 cellular partners and substrates have been identified, and the role of PTPN3 in cell signaling remains unclear. PTPN3 was reported both as a partner and a substrate of the mitogen-activated protein kinase (MAPK) p38γ, involved in Ras oncogenesis (Hou et al., 2012). PTPN3 was also reported in breast cancer by dephosphorylating the epidermal growth factor receptor (EGFR), increasing sensitivity to tyrosine kinase inhibitors (Ma et al., 2015), and by regulation of the vitamin D receptor expression and stability, which stimulates breast cancer growth (Zhi et al., 2011). PTPN3 also regulates the activity and expression of the Tumor necrosis factor alpha-convertase (TACE) protein, which impacts the release of soluble tumor necrosis factor α (TNF-α) (Zheng et al., 2002). In that case, PTPN3 was reported to bind TACE through a PDZ-PBM interaction. Interestingly, viral proteins of oncoviruses, such as the capsid protein (HBc) of hepatitis B virus (HBV) (Hsu et al., 2007) and the E6 protein of high-risk human papillomaviruses types 16 and 18 (HPV16 and 18) (Töpffer et al., 2007), possess PBMs that are able to interact with PTPN3. Targeting host PDZ domains through PBMs is a strategy developed by many viruses to hijack cellular machinery to their advantage (James and Roberts, 2016). As previously shown for the rabies glycoprotein (Préhaud et al., 2010; Caillet-Saguy et al., 2015), viruses can compete with endogenous partners through their viral PBMs to disturb signaling pathways in infected cells. We previously studied the interaction between PTPN3-PDZ and the PBMs of HPV16 E6 (PBM-16E6) (Genera et al., 2019) and HBV HBc (PBM-HBc)(Genera et al., 2021). The PDZ-mediated interaction of E6 with PTPN3 results in the proteasomal degradation of the phosphatase (Jing et al., 2007). In the case of HBV, PTPN3 can impact multiple stages of the HBV life cycle and interacts with viral capsids (Genera et al., 2021), although the functional role of this interaction, notably in cell signaling, has not been fully established. Indeed, the biophysical and structural studies on PTPN3 have mainly focused on the PTP domain in complex with phospho-peptide substrates derived either from the MAPK p38γ (Chen et al., 2014) or the EGFR substrate 15 (Chen et al., 2015), independently from the PDZ. Therefore, the interactions mediated by PTPN3-PDZ with cellular and viral proteins should be documented to further understand its function in cellular signaling.

**Figure 1.**
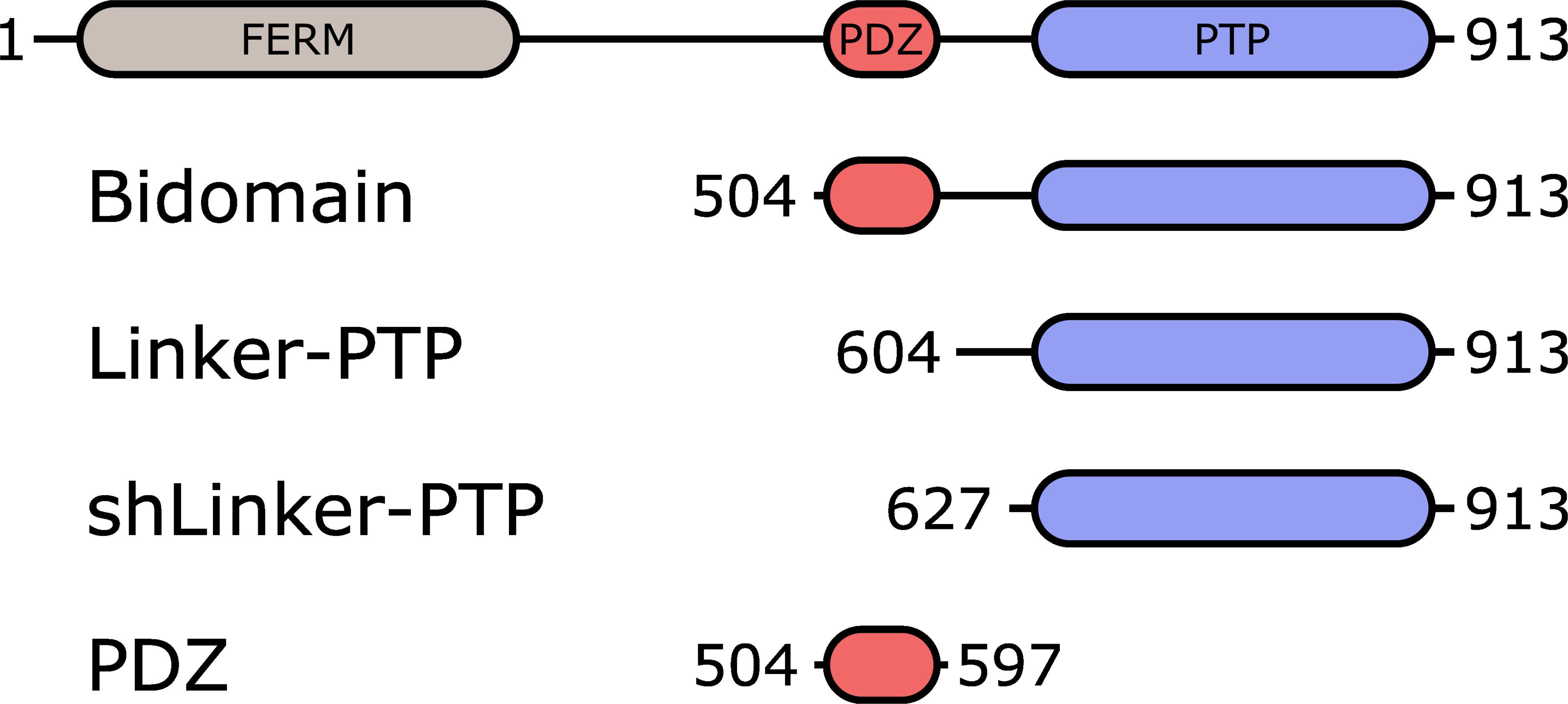
Schematic representation of PTPN3 and the constructs. The FERM, PDZ and PTP domains are represented. The boundaries of the full-length and the constructs are shown. Bidomain encompasses the PDZ and the PTP domains; Linker-PTP contains the linker and the PTP domain; shLinker-PTP contains a shorter linker and the PTP domain.

Here we performed a structural and functional study of PTPN3 and its PDZ-mediated interactions. We solved the crystal structures of the complexes formed by PTPN3-PDZ with peptides comprising the PBMs of HPV 18 E6 (PBM-18E6) and of the cellular partner TACE (PBM-TACE). We compared these structures with the ones with PBM-16E6 and PBM-HBc to highlight the atomic determinants of PTPN3-PDZ/PBM recognition. We identified the crucial positions that define the selectivity of this domain focusing on the specificity determinants shared with its close homolog PTPN4. Indeed, PTPN3 and PTPN4 compose the NT5 subfamily of non-receptor PTPs and share the same modular organization with 73% of sequence identity in their PDZ domains.

We performed bioinformatics studies to analyze the specificity of recognition of PTPN3-PDZ and PTPN4-PDZ by comparing alignments of PTPN3 and PTPN4 PDZ domains orthologous sequences versus the alignment of all human PDZ domains. We identified conserved positions that could be involved in the specificity of recognition of PBMs by these phosphatases. Additionally, sequence analysis of the PBMs captured by PTPN3-PDZ from cell lysates provided insights on the consensus sequence preferentially bound in this context. We also analyzed the PDZ domains preferentially targeted by PTPN3’s PBMs from our holdup high-throughput binding assay (Vincentelli et al., 2015).

Finally, we characterized the regulation of PTPN3 phosphatase activity with a focus on the impact of the linker between the PDZ and the PTP domains, the inhibition by the PDZ domain, and the potential effects of the binding of PBMs of its cellular and viral partners.

## 2 Materials and methods

### 2.1 Production and Purification of Recombinant Proteins

PTPN3-PDZ is encoded as an N-terminal gluthathione S-transferase (GST) tagged protein in a pDEST15 expression plasmid. PTPN3-Bidomain and PTPN3-linker-PTP are encoded as Nterminal 6xHis tagged proteins in pET15b expression plasmids. In the three cases, a TEV cleavage site is inserted between the N-terminal tags and the protein sequences. The vectors were used to transform E. coli BL21 Star (DE3) star cells (Invitrogen, Carlsbad, CA, USA).

PTPN3-PDZ, PTPN3-Bidomain and PTPN3-linker-PTP constructs were expressed and purified as previously described (Maisonneuve et al., 2014) with minor modifications.

Briefly, harvested cells were resuspended in buffer A (50 mM Tris/HCl, pH 7.5, 150 mM NaCl), 2mM β-mercaptoethanol and protease inhibitor cocktail (Roche), and then disrupted in a French press. The clarified supernatants were loaded onto a GST column (GSTrap HP, GE Healthcare) or a nickel affinity chromatography column (HiTrap HP, GE healthcare) and washed with the same buffer. The GST tag was cleaved by overnight incubation at 4°C by TEV protease (1% mol/mol) directly injected into the column. The eluted fractions containing the protein were pooled and loaded onto a size exclusion column (HiLoad Superdex 75 pg; GE) equilibrated with buffer A with 0.5 mM Tris(2-carboxyethyl)phosphine (TCEP). Purified proteins were concentrated using centrifugal filter devices (Vivaspin, Sartorius). Protein concentration was estimated from its absorbance at 280 nm. Purification of PTPN3-PDZ used for crystallogenesis was performed as previously reported (Genera et al., 2021).

The peptides, PBM-p38γ, PBM-HBc, PBM-16E6 and PBM-18E6, were synthesized in solid phase using Fmoc strategy (Proteogenix) and resuspended in H2O with pH adjusted.

### 2.2 NMR experiments

The NMR binding experiments between PTPN3-PDZ and PBM-TACE peptide to measure PTPN3-PDZ·PBM peptide affinities were performed at 15 °C on a 600-MHz Bruker Avance III HD spectrometer equipped with a cryoprobe. Briefly, the PBM-TACE peptide (stock solution at 3 mM mM pH 7.5) was added stepwise in a sample initially containing ^15^N-labeled PTPN3-PDZ at a concentration of 117 μM. A series of ^1^H, ^15^N HSQC spectra was recorded for 11 different titration points with a ratio PDZ:PBM-TACE (mol:mol) from 1:0 to 1:8.5. The NMR samples for the PTPN3-PDZ was prepared in buffer A with 0.5 mM TCEP and D2O (5% vol:vol). The chemical shift changes were followed with the CcpNmr Analysis software (27)(Vranken et al., 2005). Average (^1^H, ^15^N) chemical shift changes were calculated as Δ*δ_av_* = [(Δ*δ_H_*)^2^ + (Δ*δ_N_* × 0.15)^2^]^1/2^.

The K_D_ was obtained by fitting the titration data with a model assuming a 1:1 complex formation and with nonlinear regression using CcpNMR Analysis software. The (^1^H, ^15^N) chemical shift changes were fitted in function of the ratio ligand/protein using the equation: 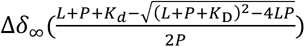, where Δδ_∞_ is the average chemical shift changes for each titration point, Δδ_∞_ is the maximal variation of the chemical shift, P is total protein concentration and L is the total ligand concentration.

A pool of 14 peaks with the best fit for each titration were kept to deduce the K_D_, and the errors are the standard deviations of all the K_D_ values fitted from the curves. Signals broaden in the moderate fast-exchange regime observed with PTPN3-PDZ and the PBM peptide, increasing the experimental errors on the chemical shift measurements used for the fitting of the K_D_.

### 2.3 Crystallization, data collection, and structure determination

The PDZ domain-peptide complexes for co-crystallization was generated by mixing PTPN3-PDZ in 20 mM HEPES pH 8, 150 mM NaCl, 0.5 mM TCEP and the peptide at a ratio of 1:2. The PDZ domain concentrations were at 5 mg/mL and 4.8 mg/mL for the complex with PBM-18E6 and PBM-TACE, respectively.

Crystallization trails were performed in 400 nanoliter sitting-drop vapor diffusion method by using Mosquito nanolitre-dispensing crystallization robot at 18°C (TTP Labtech, Melbourn, UK) and following established protocols at the Crystallography Core Facility of Institut Pasteur in Paris, France (Weber et al., 2019).

The best crystals were obtained for the complex with PBM-18E6 in crystallization condition containing 20% w/v PEG 3350, 0.2 M NaI at pH 7, and for the complex with PBM-TACE, the reservoir solution contained 20% w/v PEG 3350, 0.2 M Na-thiocyanate at pH 7. Crystals were cryo-protected in a 1:1 v/v mixture of paraffin oil and paratone oil. X-ray diffraction data were collected at a wavelength of 0.979 Å on the beamline PROXIMA-1 at Synchrotron SOLEIL (St. Aubin, France). The data were processed with XDS (Kabsch, 2010), and the programs Pointless (Evans, 2011) and Aimless (Evans and Murshudov, 2013) from the CCP4 suite (Winn et al., 2011). The structures were solved by molecular replacement with PHASER (McCoy, 2007) using as the search model PTPN3-PDZ (PDB ID 6HKS).

The locations of the bound peptides were determined from a Fo–Fc difference electron density maps. Models were rebuilt using COOT (Emsley et al., 2010), and refinement was done with phenix.refine (Adams et al., 2010). The overall assessment of model quality was performed using MolProbity. The crystal parameters, data collection statistics, and final refinement statistics are shown in Table 2. All structural figures were generated with the PyMOL Molecular Graphics System, Version 1.7 (Schrödinger)(Figure 2).

**Figure 2.**
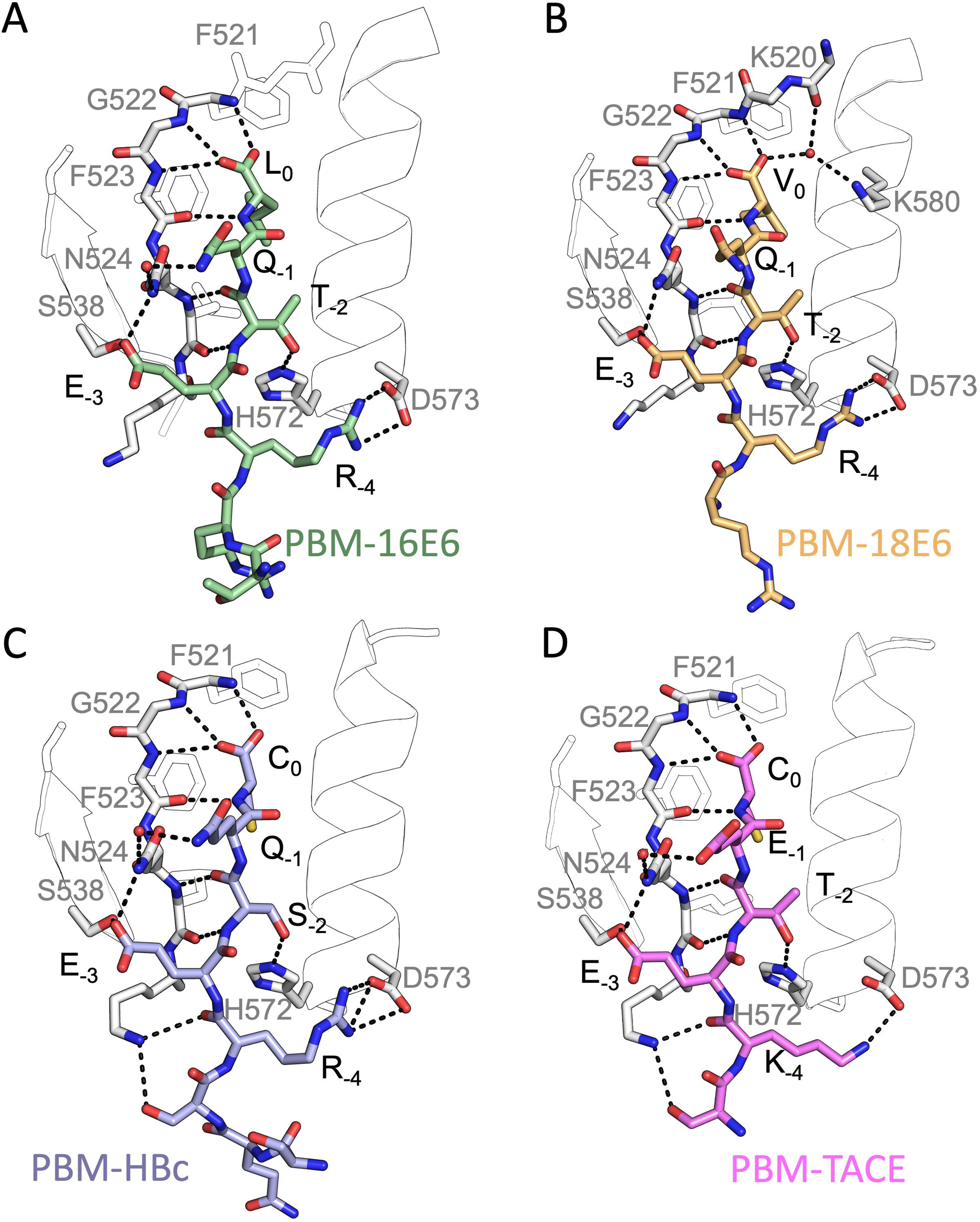
Insights of the binding network of PTPN3-PDZ with the PBM peptides of HPV16 E6, HPV18 E6, HBc and TACE. Secondary structure elements of PTPN3-PDZ are shown in white trace, and relevant residues are shown in white sticks. Peptides are shown as coloured sticks, with the corresponding name indicated below the structure. Polar interactions are shown as black dashes.

The atomic coordinates and structure factors of PTPN3-PDZ domain in complex with PBM-TACE and PBM-18E6 have been deposited in the Protein Data Bank under accession codes 8CQY and 8OEP, respectively.

### 2.4 Holdup assay

The holdup assay was conducted against the biotinylated peptide PBM-p38γ (Supplementary Material 1) following previously established protocols (Vincentelli et al., 2015; Duhoo et al., 2019) with some slight modifications. In summary, we utilized a high-throughput technique to measure the affinities and specificities of motifs in a library of human PDZ domains, which involved the use of both robotic and microfluidic methodologies. To achieve this, we expressed 266 PDZ domains fused with the Maltose Binding Protein (MBP) tag, which constituted 97% of the human PDZome. Bacterial extracts containing overexpressed PDZ domains were incubated with PBM peptide-coated resins in 96-well plates, then filtered and evaluated using microfluidic capillary electrophoresis to determine binding intensities (BIs) in the flowthroughs. The minimal BI threshold value is 0.2 to define a significant interaction as previously reported (Vincentelli et al., 2015). For PBM-HBc, we used our previously reported data (Genera et al., 2021). Our holdup data were recently assembled into an open-access database (https://profaff.igbmc.science)(Gogl et al., 2022).

### 2.5 Sequence analysis

For all protein alignments, logo representations were created using the online WebLogo service357 at https://weblogo.berkeley.edu/. The sequence conservation is shown as a frequency plot. The amino acids are colored according to their chemical properties: polar amino acids (G,S,T,Y,C,Q,N) are green, basic (K,R,H) blue, acidic (D,E) red and hydrophobic (A,V,L,I,P,W,F,M) amino acids are black. Letter width is scaled depending on position occupancy, with reduced width for increasing gap percentages.

Protein sequences for PTPN3 (349) and PTPN4 (354) orthologs have been retrieved from the NCBI gene databank, using one sequence per orthologous gene. Sequences are accessible at : https://www.ncbi.nlm.nih.gov/gene/5774/ortholog/?scope=7776&term=PTPN3 and https://www.ncbi.nlm.nih.gov/gene/5775/ortholog/?scope=89593&term=PTPN4. Sequences for the two proteins orthologs were pooled (703 in total) and aligned using the E-INS-i algorithm from the MAFFT package (Katoh and Standley, 2013), suitable for multidomain proteins. Positions ungapped in the human PTPN3 protein sequence (NCBI entry NP_002820.3; UniprotKB entry P26045) were manually selected for display (Figure 3A).

**Figure 3.**
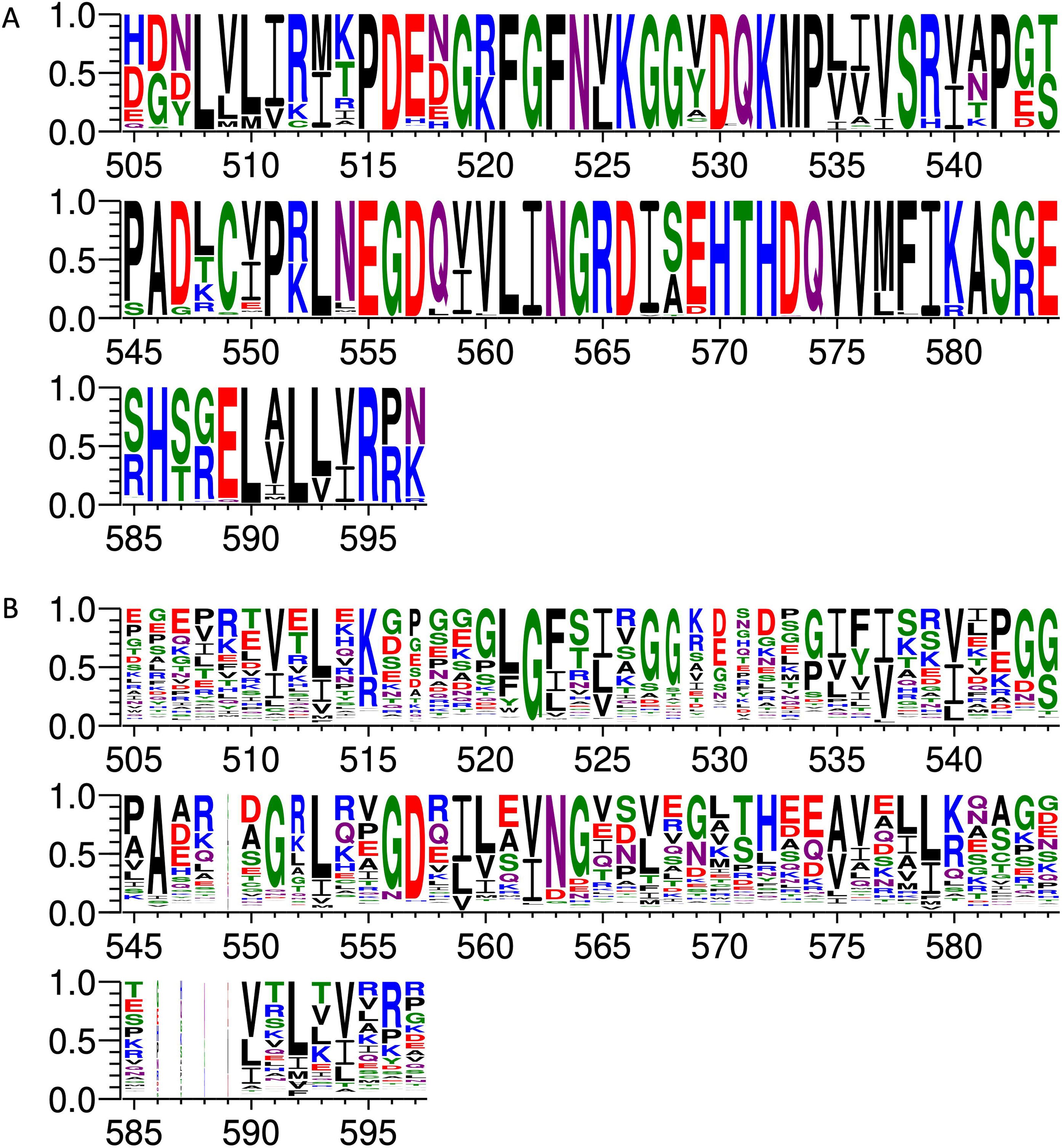
Comparison of PDZ domain sequence conservations from PTPN3 and PTPN4 orthologs and of the human PDZome. The result is displayed using a logo representation (Crooks et al., 2004), where the height of each residue one-letter code translates to its conservation at the corresponding position of the PTPN3-PDZ numbering in the sequence alignment. Sequence logo showing the sequence conservation of A) PTPN3 and PTPN4 orthologs and of B) the human PDZome.

Protein sequences for the human PDZome were retrieved from the database used for the holdup library (Vincentelli et al., 2015; Duhoo et al., 2019) and aligned using the G–INS-i algorithm from the MAFFT package, suitable for compact single domain alignment. Positions ungapped in the human PTPN3 protein sequence were manually selected for display (Figure 3B).

Sequences for HBV core (251), HPV16 E6 (1415) and HPV18 E6 (93) proteins have been retrieved from UniprotKB, respectively accessible at : https://www.uniprot.org/uniprotkb?query=HBVgp4, https://www.uniprot.org/uniprotkb?query=(protein_name:E6)%20AND%20(organism_id:333760), https://www.uniprot.org/uniprotkb?query=(protein_name:E6)%20AND%20(organism_id:333761). Protein sequences for TACE (283) and p38gamma (259) have been retrieved from the NCBI gene databank, using one sequence per orthologous gene. Sequences are accessible at: https://www.ncbi.nlm.nih.gov/gene/6868/ortholog/?scope=89593&term=ADAM17 and https://www.ncbi.nlm.nih.gov/gene/6300/ortholog/?scope=7776&term=MAPK12. For each of the 5 forementioned proteins, the last 5 residues were retrieved for display (Figure 4).

**Figure 4.**
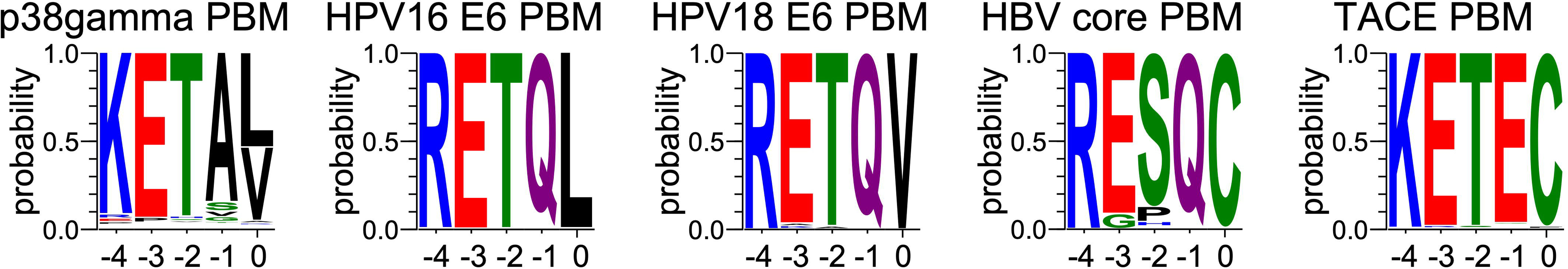
Comparison of sequence conservations of PBMs of p38γ, HPV16 E6, HPV18 E6, HBc and TACE. For each extension, all UniprotKB sequences of the corresponding protein were aligned. The result is displayed using a logo representation (Crooks et al., 2004), where the height of each residue one-letter code translates to its conservation at the corresponding position in the sequence alignment. Amino acids are coloured according to their chemical properties: polar amino acids (G,S,T,Y,C,Q,N) are green, basic (K,R,H) blue, acidic (D,E) red and hydrophobic (A,V,L,I,P,W,F,M) amino acids are black. The positions of the PBM are indicated above the sequences.

Sequences from proteins identified by pull-down and mass-spectrometry (Figure 5) were retrieved from the UniprotKB database. C-terminal PBMs were sorted according to class consensus (class1: [ST][X][ACVILF]; class2: [VLIFY][X][ACVILF]; class3: [ED][X][ACVILF], where X corresponds to any residue).

**Figure 5.**
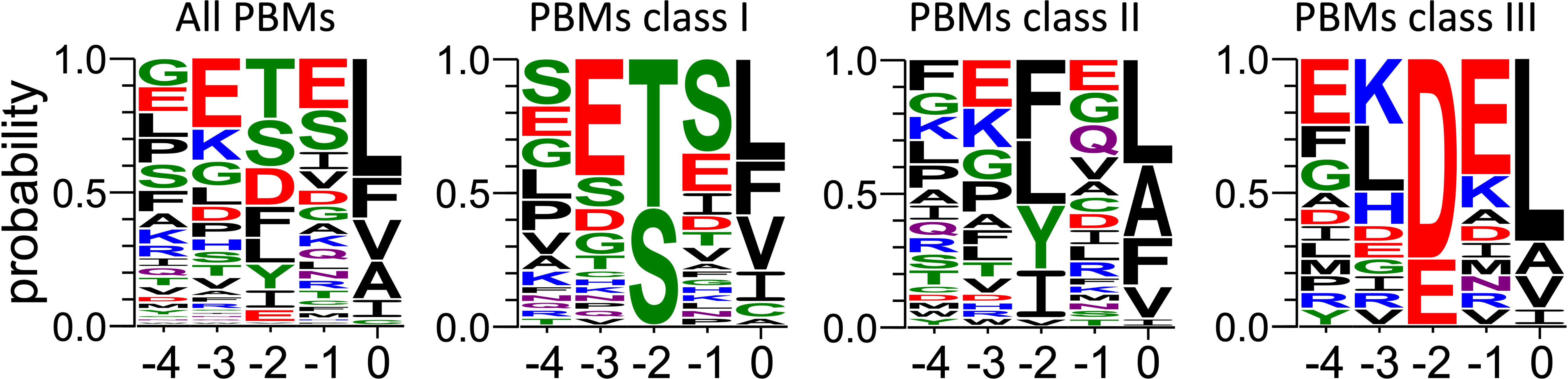
Frequency plot of the C-terminal residues of the PBM-containing partners of PTPN3-PDZ identified from cell lysate by pull-down. For each extension, all UniprotKB sequences were aligned (Supplementary Material 3). The result is displayed using a logo representation (Crooks et al., 2004), where the height of each residue one-letter code translates to its conservation at the corresponding position in the sequence alignment. Amino acids are coloured according to their chemical properties: polar amino acids (G,S,T,Y,C,Q,N) are green, basic (K,R,H) blue, acidic (D,E) red and hydrophobic (A,V,L,I,P,W,F,M) amino acids are black. The positions of the PBM are indicated above the sequences.

Protein sequences for PDZ domains recruited by HBV core (Genera et al., 2021) and p38gamma PBMs (Supplementary Material 1) in the Holdup experiments were aligned using the G–INS-i algorithm from the MAFFT package. Positions ungapped in the human PTPN3 protein sequence were manually selected for display (Figure 6).

**Figure 6.**
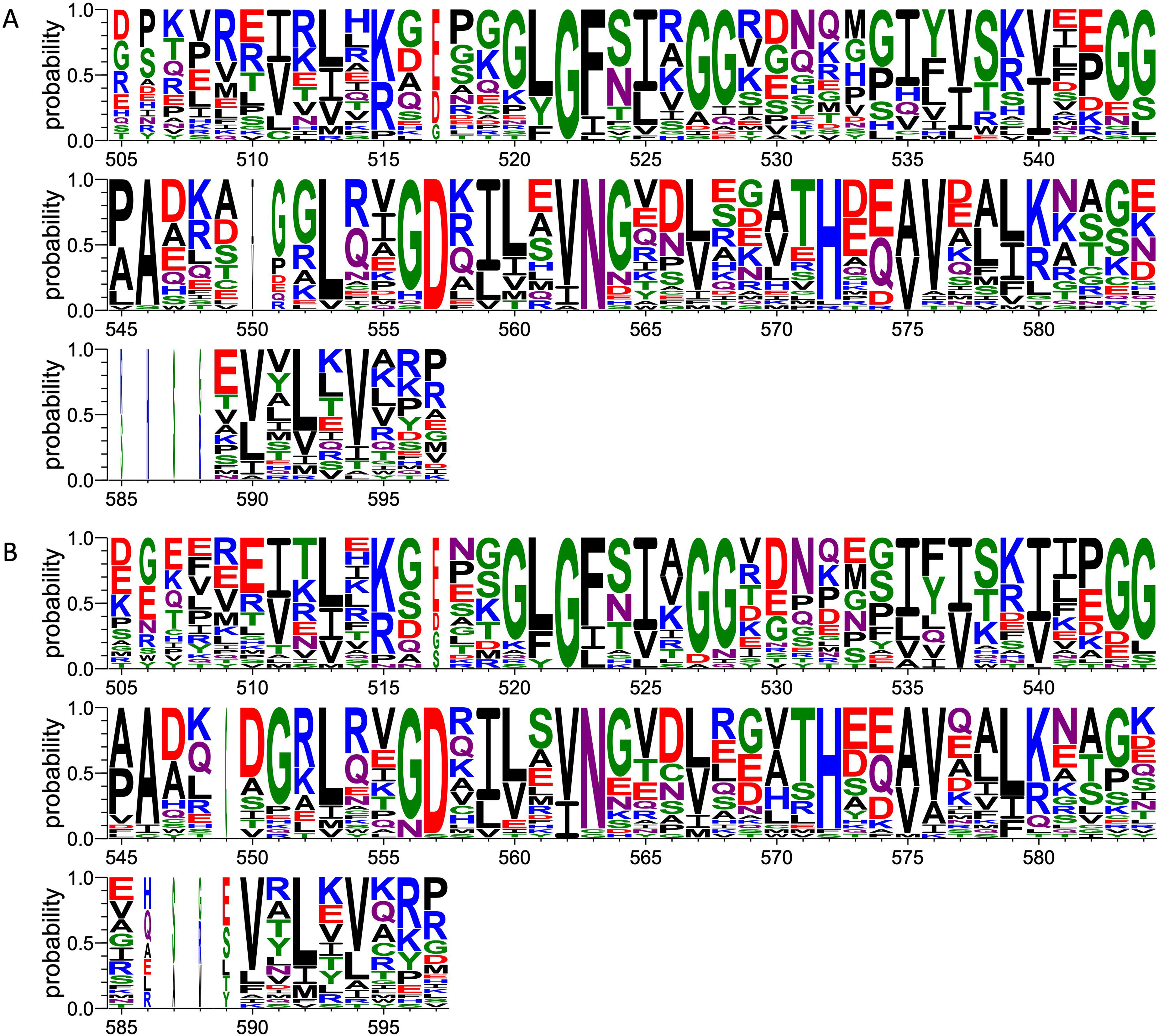
Comparison of PDZ domain sequence conservations recruited by PBM-p38γ and PBM-HBc. The result is displayed using a logo representation (Crooks et al., 2004), where the height of each residue one-letter code translates to its conservation at the corresponding position of the PTPN3-PDZ numbering in the sequence alignment. Sequence logo showing the sequence conservation of PDZ domains recruited during holdup experiment by A) PBM-p38γ and B) by PBM-HBc.

The analysis of human PBMs composition was performed based on the Swiss-prot databank.

All protein sequences for *homo sapiens* were downloaded, resulting in 20402 sequences, reduced to 20374 by removing entries of less than 20 residues.

For each sequence, the last 3 residues are considered for sorting according to PBM class consensus (class1: [ST][X][ACVILF]; class2: [VLIFY][X][ACVILF]; class3: [ED][X][ACVILF], where X corresponds to any residue) or as non-PBM. For the list of all PBM-containing proteins, the last 5 residues are extracted for the logo representation (Figure S1). The occupancy of residues at each PBM position were calculated as raw percentages, *i.e.* the number of sequences with a given residue a at position divided by the total number of considered sequences.

### 2.6 Enzymatic assays

PTPN3-Bidomain, PTPN3-linker-PTP, PTPN3-shLinker-PTP (short linker, missing 23 residues) and PTPN3-PDZ (Figure 1). In all experiments, the phosphatase activity was assessed using the synthetic nonspecific phosphatase substrate p-nitrophenyl phosphate (pNPP), whose hydrolysis into p-nitrophenol (pNP) can be followed spectrophotometrically at 410 nm. Reactions were performed in 50mM Tris-HCl, pH 7.5, 1 mM MgCl2, 150 mM NaCl, 0.5 mM

TCEP. The initial reaction rates were measured independently at pH 7.5 and at 25°C. pNPP was assayed for concentrations ranging from 19μm to 10 mM at an enzyme concentration of 75 nM. The dephosphorylation reaction followed Michaelis-Menten kinetics and exhibited a substrate inhibition effect at high concentrations of pNPP. The experimental data was therefore fitted to a corrected Michaelis-Menten equation to take into consideration this inhibition. Phosphatase activity was measured by following the hydrolysis of pNPP as previously described (Maisonneuve et al., 2014). Absorbances were measured continuously at 410 nm for pNP, using a Thermo Scientific UV spectrometer equilibrated at 25 °C. Initial linear reaction rates were calculated during a 60 second reaction. The k_cat_ and K_M_ constants were deduced from fitting the Michaelis-Menten equation with the Prism software. K_M_, k_cat_ and k_cat_/K_M_ are listed in table 3. The data are representative of three independent experiments.

A large excess (molar ratio 600:1) of each peptide was incubated with PTPN3-Bidomain for 30’ at 25°C, and the initial rates of the dephosphorylation reaction were measured in the same range of pNPP concentrations.

Then, to assess whether the linker that connects the PDZ and PTP domains is required for the catalytic regulation, we measured at 25°C the catalytic activity at an enzyme concentration of 75 nM and at a fixed concentration of 2.5 mM pNPP, where the Bidomain and linker-PTP constructs exhibited the highest initial rate of reaction (Figure 7A). We compared the initial rate for PTPN3-Bidomain, PTPN3-linker-PTP alone, and for PTPN3-linker-PTP with a large excess PTPN3-PDZ added in *trans* (molar ratio 80:1) and incubated for 1h at 4°C.

**Figure 7.**
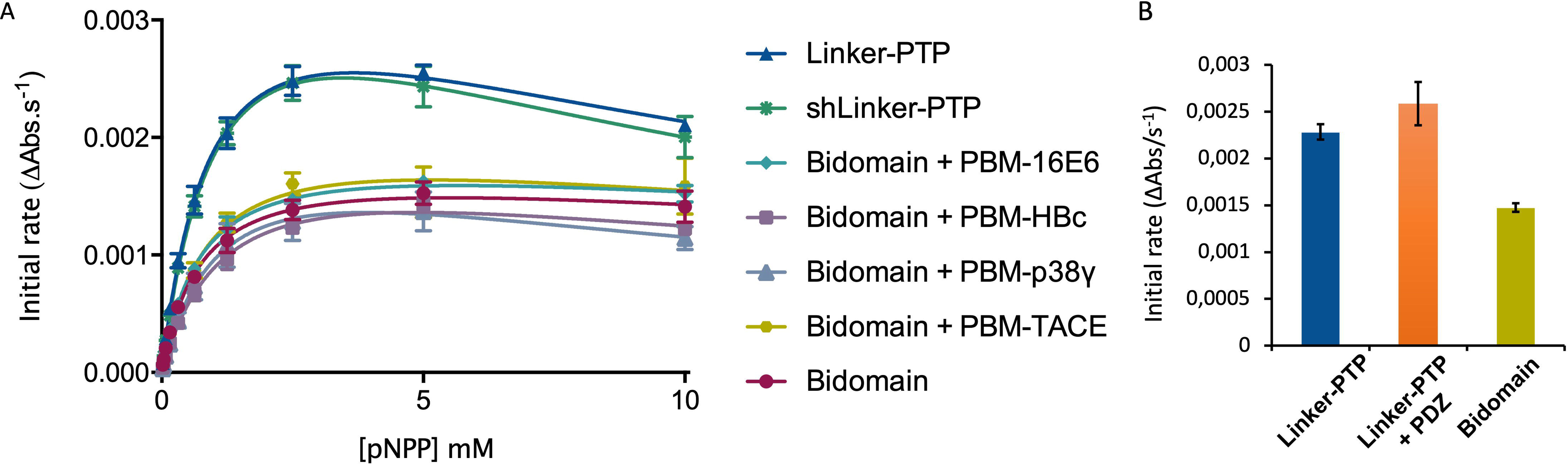
Regulation of the phosphatase activity of PTPN3. (**A**) Michaelis-Menten plots of the initial rates of pNPP hydrolysis by PTPN3 constructs. The K_M_ and *k*_cat_ constants were deduced by fitting the data to a modified Michaelis-Menten equation, considering the substrate inhibition observed at high concentrations of pNPP. The data and error bars are representative of three independent experiments. The curves are nonlinear fits to a substrate-inhibition equation. (**B)** Initial rates of pNPP dephosphorylation at 2.5 mM pNPP by PTPN3-linker-PTP, PTPN3-linker-PTP pre-incubated with PTPN3-PDZ, and PTPN3-Bidomain. PTPN3-linker-PTP and PTPN3-Bidomain were at a concentration of 75 nM, while PTPN3-PDZ was added at a concentration of 6 µM. The data and error bars are representative of three independent experiments.

### 2.7 Analytic ultracentrifugation (AUC) experiments

Sedimentation velocity experiments of PTPN3-Bidomain were carried out at 20°C using an analytical ultracentrifuge (Beckman Coulter Optima AUC) equipped with a AN60-Ti rotor. The protein sample at 14 μM was centrifuged for 17h at 42000 rpm. Data were analyzed with SEDFIT 15.1 (Schuck, 2000) using a continuous size distribution c(S) model. The partial specific volume, the viscosity and the density of the buffer were calculated with SEDNTERP.

### 2.8 Small Angle X-Ray Scattering (SAXS) experiments

To minimize the contribution of small aggregates to the scattering, synchrotron radiation X-ray scattering data were collected on the SWING beamline at Synchrotron Soleil (France) using the online HPLC system. SAXS samples were injected into a size exclusion column (superdex 75 increase 5 x 150 Cytiva) using an Agilent High Performance Liquid Chromatography system cooled at 25 °C and eluted directly into the SAXS flow-through capillary cell at a flow rate of 200 μL·min^−1^. For the experiments corresponding to PTPN3 bidomain complexed with PBM-p38γ, the column was equilibrated with buffer containing 40 μM PBM-p38γ. SAXS data were collected online throughout the whole elution time, with a frame duration of 1s. The first 100 frames collected during the first minutes of the elution flow were averaged to account for buffer scattering. The 10 frames corresponding to the top of the elution peak were averaged and were used for data processing after baseline subtraction (see Supplementary Material 2).

The data were analyzed using foxtrot and primus from atsas (Konarev et al., 2003) suite, from which Guinier was generated. From the corrected scattering curves, the pair distribution functions were computed using gnom.

Models for PTPN3 Bidomain were generated using CORAL from residue 488 to 913 based on the X-ray structure of the PDZ (PDB id 6T36) and the catalytic domain (PDB id 2B49). The two domains were rigid. The linker, the N-terminal, and the C-terminal part of the PTPN3 were set in random conformations. 50 models were generated, and the best model is presented. The model of PTPN3 complexed with PBM-p38γ was generated similarly with the structure of the PDZ complexed with the PBM.

## 3 Results

### 3.1 Similar affinities of cellular and viral PDZ-binding motifs for PTPN3-PDZ suggest viral mimicking of cellular PBM sequences

We determined the dissociation constant (K_D_) of PTPN3-PDZ for the PBM-TACE peptide comprising the last C-terminal 12-residues of TACE and encompassing the PBM (sequence RQNRVDSKETEC)(Table 1) following the ^1^H, ^15^N chemical shift perturbations of PTPN3-PDZ NMR signals in the ^1^H-^15^N HSQC spectra as a function of peptide concentration (Figure S2). The PBM-TACE peptide binds to PTPN3-PDZ with a K_D_ value of 30 μM. We previously obtained K_D_ values of 26 μM, 29 μM, 53 μM and 37 μM for PBM-p38γ, PBM-HBc, PBM-16E6 and PBM-18E6, respectively, using the same methodology (Table 1). Thus, the PBM-TACE K_D_ value is similar to the K_D_s previously measured for the PBM peptides of other cellular or viral partners (Genera et al., 2019) falling in the few tenth-of-micromolar range, common for PDZ-PBM interactions.

**Table 1.**
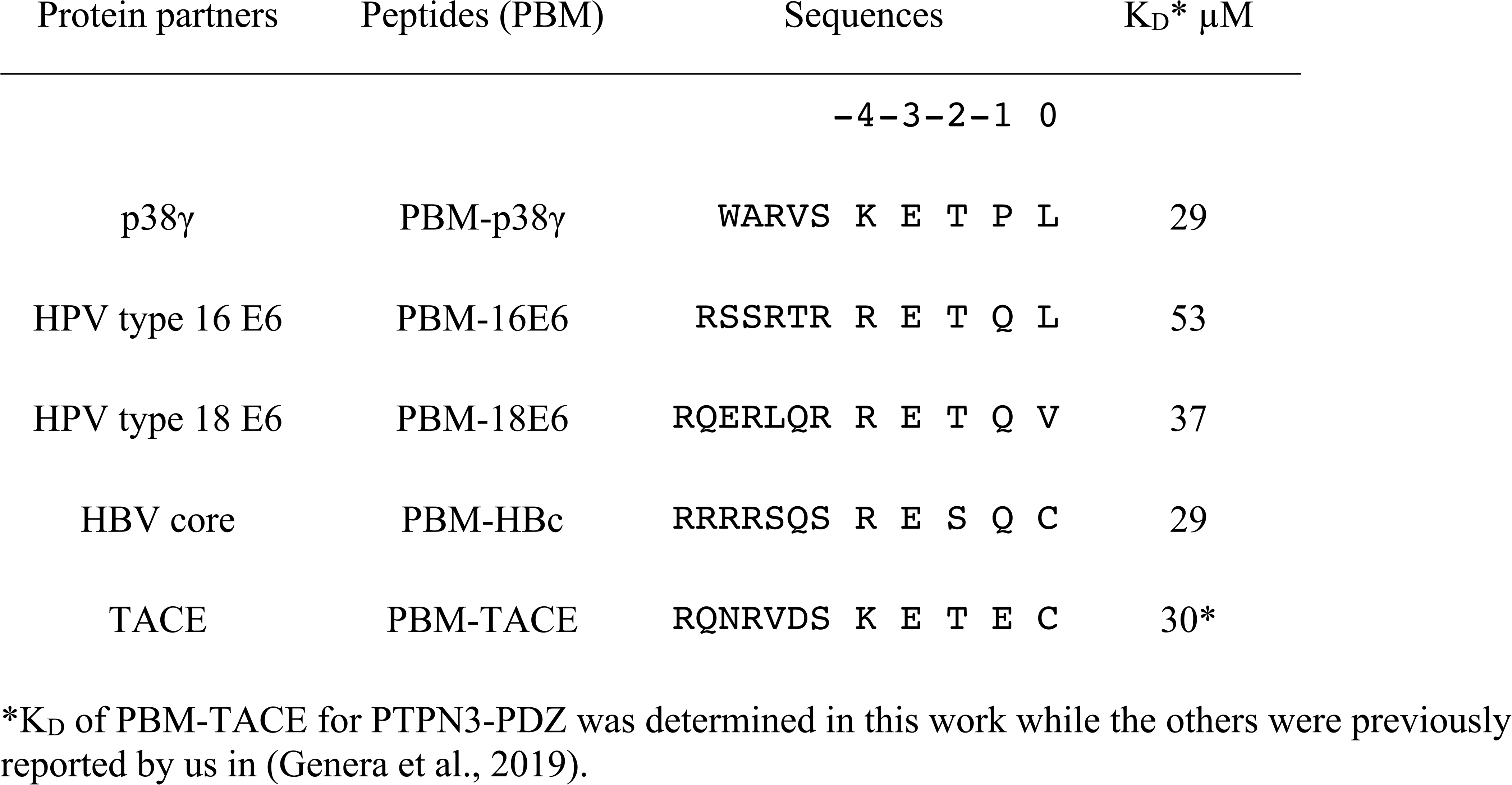
Affinities and peptide sequences of PBMs of cellular and viral partners of PTPN3-PDZ.

All these PBMs are type 1 PBMs with the canonical consensus sequence S/T-X-Φ_COOH_ (where X is any residue and Φ is a hydrophobic residue). Interestingly, the cellular and viral sequences have strong similarities in their sequences at the C-terminal positions (Table 1). Indeed, an atypical cysteine is found at the last position (position 0 or P0) for both the viral PBM-HBc and the cellular PBM-TACE, while common leucine and valine are found for viral HPV PBMs and the cellular p38γ. In addition, we observed conserved positions in all PBMs with a negative aspartic acid at position -3 (P-3) and a positive arginine or lysine at P-4. This conservation among the viral and cellular sequences is in agreement with a viral mimicking of the cellular PBM sequences, allowing the viral proteins to interact with host PDZ domains with similar affinities to cellular partners, as observed here for PTPN3-PDZ. This result is consistent with the hypothesis that viral PBM sequences can compete with endogenous ligands of PTPN3-PDZ, disrupting cellular PDZ/PBM complexes.

### 3.2 Crystallographic studies of PTPN3-PDZ and its binding specificities to PBMs

To explore the binding specificities of PTPN3-PDZ towards PBMs, we conducted crystallographic studies of PTPN3-PDZ in complex with PBM-18E6 and PBM-TACE peptides (Table 1). The statistics for data collection and refinement are provided in Table 2. These structures were compared to our previously published crystal structures of PTPN3-PDZ in complex with PBM-16E6 (Genera et al., 2019) and PBM-HBc (Genera et al., 2021).

**Table 2.** Data collection and refinement statistics.

| PTPN3-PDZ in complex with PBM peptides |  |  |
| --- | --- | --- |
|  | PBM-18E6 | PBM-TACE |
|  | RQERLQRRETQV | RQNRVDSKETEC |
| **Wavelength** | 0.9786 | 0.98 |
| **Resolution range** | 31.7 - 1.873 (1.94 - 1.873) | 66.58 - 1.7 (1.761 - 1.7) |
| **Space group** | P 65 2 2 | P 32 2 1 |
| **Unit cell** | 82.23 82.23 139.2 90 90 120 | 76.878 76.878 46.253 90 90 120 |
| **Total reflections** | 291759 (27178) | 244763 (23662) |
| **Unique reflections** | 23304 (2137) | 17566 (1726) |
| **Multiplicity** | 12.5 (12.7) | 13.9 (13.7) |
| **Completeness (%)** | 98.70 (93.07) | 99.50 (98.40) |
| **Mean I/sigma(I)** | 18.12 (1.91) | 12.51 (1.57) |
| **Wilson B-factor** | 33.51 | 42.30 |
| **R-merge** | 0.08981 (1.385) | 0.1035 (1.433) |
| **R-meas** | 0.09363 (1.443) | 0.1079 (1.489) |
| **R-pim** | 0.02571 (0.3905) | 0.02982 (0.3998) |
| **CC1/2** | 0.999 (0.846) | 0.997 (0.887) |
| **CC\*** | 1 (0.957) | 0.999 (0.97) |
| **Reflections used in refinement** | 23304 (2134) | 17573 (1726) |
| **Reflections used for R-free** | 1164 (107) | 903 (80) |
| **R-work** | 0.1983 (0.3067) | 0.1576 (0.2825) |
| **R-free** | 0.2488 (0.3477) | 0.1929 (0.3172) |
| **CC(work)** | 0.954 (0.845) | 0.816 (0.522) |
| **CC(free)** | 0.926 (0.857) | 0.791 (0.461) |
| **Number of non-hydrogen atoms** | 1763 | 808 |
| **macromolecules** | 1630 | 784 |
| **ligands** | 1 | 2 |
| **solvent** | 132 | 22 |
| **Protein residues** | 202 | 99 |
| **RMS(bonds)** | 0.012 | 0.032 |
| **RMS(angles)** | 1.93 | 2.81 |
| **Ramachandran**<br>**favored (%)** | 100.00 | 95.79 |
| **Ramachandran**<br>**allowed (%)** | 0 | 4.21 |
| **Ramachandran**<br>**outliers (%)** | 0.00 | 0.00 |
| **Rotamer outliers (%)** | 0.54 | 8.89 |
| **Clashscore** | 6.99 | 10.79 |
| **Average B-factor** | 45.16 | 43.93 |
| **macromolecules** | 45.18 | 44.09 |
| **ligands** | 84.60 | 42.06 |
| **solvent** | 44.62 | 38.20 |
| **Number of TLS**<br>**groups** | 4 | 2 |
| **PDB code** | 8OEP | 8CQY |

Statistics for the highest-resolution shell are shown in parentheses.

**Table 3.**
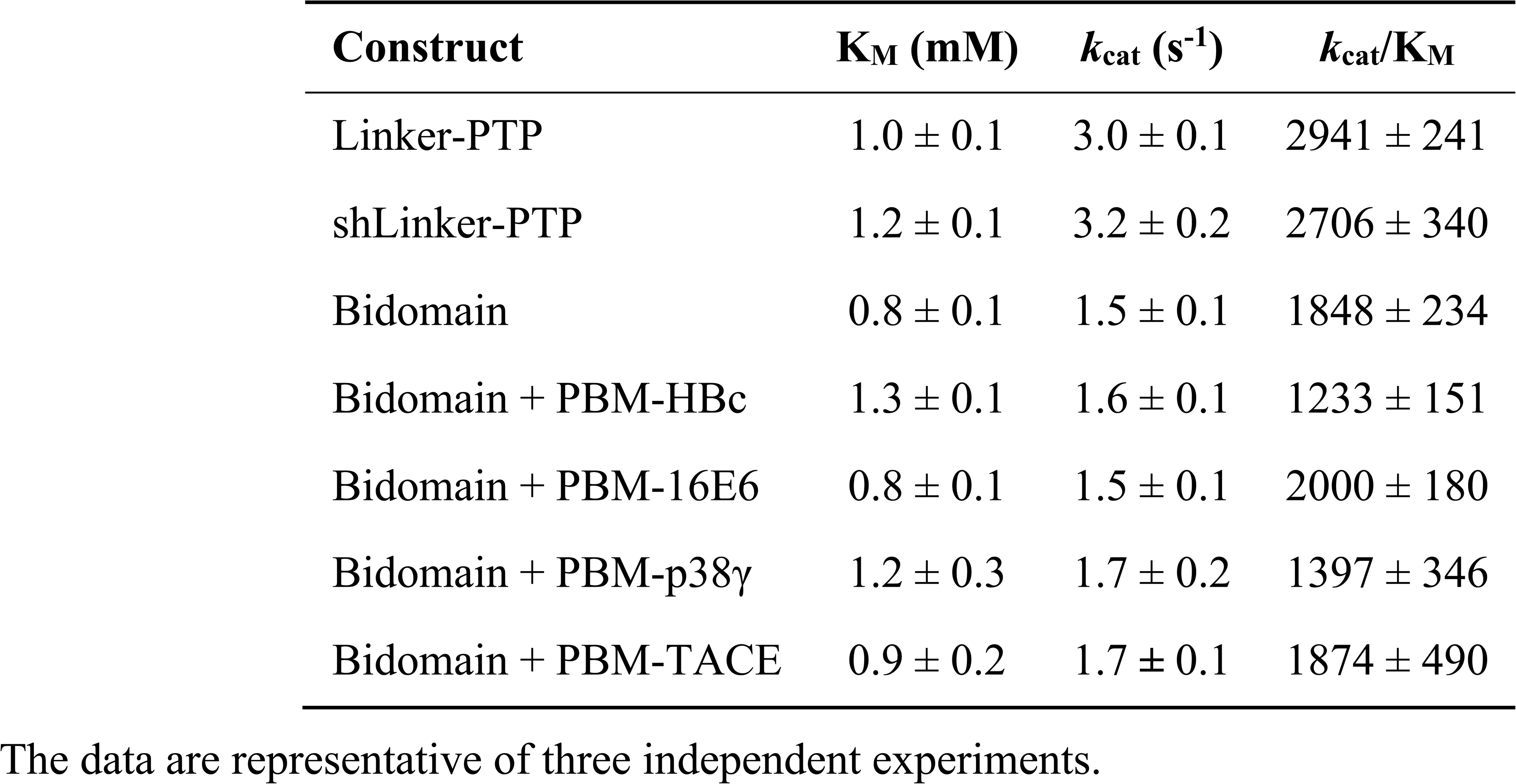
Kinetic parameters of hydrolysis of pNPP by PTPN3 constructs.

| **Construct** | **K<sub>M</sub> (mM)** | **k<sub>cat</sub> (s<sup>-1</sup>)** | **k<sub>cat</sub>/K<sub>M</sub>** |
| --- | --- | --- | --- |
| Linker-PTP | 1.0 ± 0.1 | 3.0 ± 0.1 | 2941 ± 241 |
| shLinker-PTP | 1.2 ± 0.1 | 3.2 ± 0.2 | 2706 ± 340 |
| Bidomain | 0.8 ± 0.1 | 1.5 ± 0.1 | 1848 ± 234 |
| Bidomain + PBM-HBc | 1.3 ± 0.1 | 1.6 ± 0.1 | 1233 ± 151 |
| Bidomain + PBM-16E6 | 0.8 ± 0.1 | 1.5 ± 0.1 | 2000 ± 180 |
| Bidomain + PBM-p38γ | 1.2 ± 0.3 | 1.7 ± 0.2 | 1397 ± 346 |
| Bidomain + PBM-TACE | 0.9 ± 0.2 | 1.7 ± 0.1 | 1874 ± 490 |

The data are representative of three independent experiments.

In all cases, the overall PDZ fold of PTPN3-PDZ is highly conserved, and the peptides bind in the conventional mode as an anti-parallel extension to the β2-strand (see Figure 2). Compared to our previous PTPN3-PDZ structure in complex with PBM-16E6 (PDB ID 6HKS)(Figure 2A), all structures present a very low root mean square deviation (rmsd) ranging from 0.19 Å to 0.30 Å for the backbone atoms of PTPN3-PDZ, indicating that none of these peptides induce significant conformational changes in the backbone when binding to PTPN3-PDZ. All the complexes possess the classical bonding network of class I PDZ/PBM interactions (S/T-X-Φ_COOH_). The C-terminal residues that exhibit well-defined electron density maps start from P-5 for PBM-TACE, P-6 for PBMs of HPVs or P-7 for PBM-HBc until P0 (see peptide sequences in Table 1).

The interactions at P0 and P-2 are essential in PDZ/PBM recognition. Position -2 in particular can be considered as the class determinant for PBMs (Songyang et al., 1997). The binding modes of each PBM peptide to PTPN3 are shown in Figure 2. As expected, the C-terminal carboxylate in each peptide forms three H-bonds with the amide nitrogens of F521, G522 and F523 of the “GLGF motif” on PTPN3-PDZ. The PBM-18E6 valine at P0 (V0) is additionally bonded to the carbonyl of G519 on the α1-β1 loop and to the Nζ of K580 of the α2-helix through a molecule of water (Figure 2B). At P-2, the S or T side chains form H-bonds with the Nε2 of H572 at the N-terminus of the α2-helix of PTPN3-PDZ, which is conserved in class I PDZ domains. These interactions found in these two key positions, P0 and P-2, correspond to the expected bonding pattern of a class I PBM for all the PBMs tested.

An interesting feature of PBM-TACE and PBM-HBc (Genera et al., 2021) is the presence of the C-terminal cysteine at P0 (Figure 2C,D). The carboxylate-binding pocket at the top of the peptide-binding groove of PDZ domains is lined with hydrophobic side chains (F521, F523, L525, I579), which determines the preference for peptides with C-terminal hydrophobic residue (position P0)(Songyang et al., 1997) (Figure S3A). We observed that in the complexes of PTPN3-PDZ with PBM-TACE and PBM-HBc, the cysteine side chain is oriented towards the interior of the peptide-binding groove, without making any contacts with the hydrophobic side chains that line this pocket (Figure S3B). The cysteine side chain is short enough to fit within the binding pocket and occupies the same position as the conventional leucine side chain at P0 of HPV16 E6 (Figure 2A).

At P-1, the Q side chain of PBM-16E6 and PBM-HBc and the E side chain of PBM-TACE form a H-bond with a water molecule that is in turn bonded to the Nδ2 of N524 in the β2-strand of PTPN3-PDZ. In PBM-18E6, on the contrary, the side chain does not contact the PDZ domain (Figure 2). This position is not considered as a significant determinant for PDZ/PBM interaction and is not specified in any of the three main classes of PDZ domains. In the case of PTPN3, sequence analysis of its known partners suggests a bias towards Q or E residues with rather long and polar side chains at P-1 (Table 1). In line with this, N524 is strictly conserved in PTPN3 and PTPN4 orthologs (Figure 3A), while short polar residues S or T are more often found in this position in the full human library of all the 273 known PDZ domains (PDZome)(Figure 3B). The short side chains of S and T could probably not establish bonds with Q or E at P-1. Moreover, N524 is also H-bonded to P-3 of the PBM which also requires a long side chain. Indeed, in all solved structures of complexes between PTPN3-PDZ and viral and cellular PBMs (PBM-HBc, PBM-16E6 and PBM-18E6 and PBM-TACE), N524 interacts with the E side chain at P-3 (Figure 2). This interaction is also observed between PTPN4-PDZ and the PBMs that contain E at P-3, such as the ones of p38γ (Maisonneuve et al., 2016), the attenuated rabies virus glycoprotein, and the ionotropic glutamate receptor GluN2A (Babault et al., 2011). Interestingly, the Q and E at P-1 and the E at P-3 are strongly conserved in PBMs (Figure 4). Thus, the conservation of N524 could originate from these two interactions with the -3 and the -1 residues of the PBM.

### 3.3 Exploring the impact of P-3 and P-4 on PDZ ligand selection within the NT5 subfamily

We propose that P-3 and P-4 have a significant influence on the PDZ ligand selection by the NT5 phosphatase subfamily which encompasses PTPN3 and PTPN4. The E at P-3 and R or K at P-4 are strictly or strongly conserved in ligands of PTPN3-PDZ (PBM-HBc, PBM-16E6 and PBM-18E6 and PBM-TACE, PBM-p38γ)(Figure 4). Similarly, the most affine ligands of PTPN4-PDZ also feature an E at P-3 (Babault et al., 2011). In all our PTPN3-PDZ/peptide complexes and in the previously reported PTPN4-PDZ/peptide complexes (Babault et al., 2011; Maisonneuve et al., 2016), the E side chain at P-3 forms a bifurcated H-bond with the amide nitrogen of N524 and with the side chain hydroxyl of S538 in the β3 strand (Figure 2). Like N524, S538 is strictly conserved in PTPN3 and PTPN4 orthologs (Figure 3A), while this position is conserved only in about 20% of cases in the human PDZome with also K, T and A, commonly found (Figure 3B). In addition, the aliphatic carbon chain of E at P-3 is well positioned to establish hydrophobic contacts with the carbon side chain of K526, which contributes to its stabilization. In PTPN3 and PTPN4 orthologs, a K residue is conserved at position 526, while in the PDZome, R and V are more frequently found, followed by S, A, and K (Figure 3B). In agreement with this, similar hydrophobic contacts have been observed in the complexes of PTPN4-PDZ with the PBMs of p38γ, GluN2A, and the attenuated rabies virus glycoprotein (Cyto13-att)(Babault et al., 2011; Maisonneuve et al., 2016)(Figure S4), suggesting that these contacts can also influence the PDZ ligand selection of the NT5 subfamily.

In all four PBMs (PBM-HBc, PBM-16E6 and PBM-18E6 and PBM-TACE), the long and positively charged side chains of R or K at P-4 form ionic bonds with the carboxylate oxygens of D573 at the N-terminus of the α2-helix in PTPN3-PDZ (Figure 2). This bond is also found in our previous structure of PTPN4-PDZ complexed to PBM-p38γ where a K is conserved at P-4 (Maisonneuve et al., 2016). PTPN3 and PTPN4 orthologs present a conserved D at position 573 (Figure 3A). In the PDZome, an E is most frequently found in this position (about 25%), followed by D, A, S, Q, and K (Figure 3B). Both E and D should be able to establish ionic bonds with a positively charged residue at P-4 of the PBM. We previously identified the main structural elements of PBM binding to PTPN4-PDZ and optimized the sequence of a synthetic peptide of higher affinity. P-3 and P-4 of the optimized PBM were shown to be critical with an E in P-3 forming H-bonds with the conserved S538 (S545 in PTPN4) and the R in P-4 forming H-bond with D573 (D580 in PTPN4)(Maisonneuve et al., 2016). However, G and I at P-4 in the two PTPN4-PDZ ligands, the attenuated rabies virus G protein and the glutamate receptor GluN2A, respectively, bind to PTPN4-PDZ without providing adequate side-chains to interact with D573, and the affinity of the interaction decreases accordingly (Babault et al., 2011)(Figure S4). Thus, D573 contributes to the selectivity of the NT5 family by improving affinity for PBMs with a R or K at P-4 but is not a determinant of specificity since other residues are allowed at this position.

In all the PTPN3-PDZ/peptide complexes, the amide nitrogen of R or K at P-4 forms a H-bond with the Nε2 of the Q531 side chain (loop β2-β3). Q531 is strongly conserved in PTPN3 and PTPN4 orthologs (Figure 3A), but the conservation of residue at this position is low in the PDZome; S, N, G, and H, are all found with higher frequency than Q (Figure 3B). However, although these interactions with Q531 likely contribute to the affinity of the complexes, they are not involved in NT5 subfamily ligand selectivity since they involve the backbone of the PBM, and thus any residue (except proline) could fill the position. This interaction is also observed in the complexes of PTPN4-PDZ with the attenuated rabies virus G protein and GluN2A, which have G at P-4 and I at P-4, respectively (Babault et al., 2011)(Figure S4). These residues do not establish ionic contacts with the conserved D (D573 in PTPN3) as do R and K at P-4 in PTPN3, but both still form H-bonds with Q531.

Lastly, the side chain of K526 from the β2-strand is pointed towards PBM-HBc and PBM-TACE, allowing its Nζ to form H-bonds with the carbonyl oxygen of the K or R at P-4 and with the side chain hydroxyl of S at P-5, contributing to the stability of the complex. In the two other PTPN3 partners, there is R instead of S at P-5. Forming a H-bond with the short, polar side chain of S could favor the K526 side chain to orient towards the peptide, while in the other cases it adopts an extended conformation to maximize the hydrophobic contacts with the Cβ-Cγ carbon chain of E at P-3.

In conclusion, PBM-containing partners with E at P-3 are favored because of their capacity to form H-bonds with the conserved N524 and S538 of β2 and β3 strands, respectively, as well as hydrophobic contacts with the aliphatic carbon side chain of K526 of β2 strand. Additionally, partners with R or K at P-4 can form ionic bonds with the conserved D573 from α2-helix. These interactions likely contribute to the affinity of the complex, which will favor their binding over other potential partners, but they are not mandatory for the binding to occur.

### 3.4 Sequence insights of PTPN3-PDZ captured PBMs from cell Lysates

We previously performed pull-down experiments using PTPN3-PDZ as bait to fish new cellular PBM-containing partners in HeLa S3 cell lysates (Genera et al., 2021). From 326 proteins bound exclusively to GST-PTPN3-PDZ and absent from GST controls and identified by LC-MS/MS, 83 encode for a C-terminal PBM and are potential PTPN3 interactants through PDZ/PBM interactions (Supplementary Material 3). We used these data to gain insights on the consensus sequence of PBMs preferentially bound in this cell lysate context.

Among the 83 PBM-containing partners, we identified 34 of class I (S/T-X-Φ_COOH_), 33 of class II (Φ-X-Φ_COOH_), and 16 of class III (D/E-X-Φ_COOH_) (see Supplementary Material 4, 5 and 6 respectively). We performed a sequence conservation analysis on them from P0 to P-4 (Figure 5). The preferred residues of PTPN3-PDZ at position 0 are L, F and V with 37, 13 and 13 occurrences, respectively. Together, these account for 63 over the 83 binders with L being predominant. We found 2 proteins with a C at P0, both encompassing a class I PBM: NADH-ubiquinone oxidoreductase 75 kDa subunit mitochondrial (Uniprot P28331) and PCI domain-containing protein 2 (Uniprot Q5JVF3). For the class II PBMs, we found more often L, A, and F, at P0 with 13, 9, and 7 occurrences over 33 respectively partners. P0 of class III displays mainly a L (11 occurrences overs 16).

PTPN3-PDZ is classified as a class I PDZ domain. Accordingly, S and T residues, representative of this class at P-2 of the partner PBMs, are the most abundant in all PBMs bound in the cell lysate (Figure 5). Interestingly, partners with PBMs of classes II and III are also abundantly fished, with D being the next most abundant residue at P-2 after T and S (Figure 5). Several PDZ domains were previously reported to bind both class I and class II PBMs (Kalyoncu et al., 2010). The preferred residues at P-2 for the class II partners are F, L, and Y.

At P-1, PTPN3-PDZ shows a slight tendency to bind PBMs with E or S, with S prevalent for class I PBMs and E for class II and III PBMs (Figure 5). In our crystal structures, the Q or E found in this position is bonded to N524 through a molecule of water (Figure 2). A serine would be able to bond directly to N524 thanks to its shorter side chain, which could explain why this residue would be favored in this position.

At P-3, a preference for E is observed in all PBMs (Figure 5). In fact, E is selected preferentially in classes I and II and not in class III. In class I PBMs, E is most abundant (more than 40%), while S, D and G are also found at P-3 at a 10-20% frequency. D should be able to form a H-bond with S538 in a similar way to E in our structures as E and D have side chains with similar chemical properties. S and G cannot form this bond and thus do not contribute to the interaction. Finally, the conservation at P-4 is very low with no preferential residue observed at this position.

Altogether, these data are informative about the PBMs sequences preferentially fished from a cell lysate by PTPN3-PDZ and define a preferred target motif of PTPN3-PDZ in this context.

### 3.5 Investigation of specificity profiles of PBMs of p38γ and HBc recognized by PTPN3-PDZ against the human PDZome

Then, we investigated the specificity profiles of the PBMs of the PTPN3-PDZ ligands p38γ and of HBc against the full human PDZome (library that contains all the known human PDZ domains)(Duhoo et al., 2019). PBM-HBc and PBM-p38γ harbor from P-5 to P0 the sequences -SRESQC_COOH_ and -SKETPL_COOH_, respectively. P-5 is conserved with a serine in both cases, P-4 with a lysine or arginine, and P-3 with a glutamate. We used the holdup assay, an *in vitro* automated high-throughput chromatography assay that exhibits high sensitivity for low-to-medium affinity PDZ/PBM pairs and provides affinity-based ranking of identified PDZ domains matching a profile of specificity. 12-mer peptides encompassing the protein C-terminal PBM sequences linked to a biotinyl group are used as baits to quantify the interaction between PBMs and the library of human PDZ domains expressed in *Escherichia coli* (Duhoo et al., 2019).

We generated a PDZome-binding profile of PBM-p38γ. Mean values of binding intensities (BI) ranked based on affinity are reported in Supplementary Material 1. The highest BI values indicate the PDZ domains recognized with the best affinities by the peptides used as bait. 28 PDZ domains exhibited significant binding with BI values greater than 0.2, a previously defined strict threshold (Vincentelli et al., 2015). In recent work we established the PDZome-binding profile of PBM-HBc with 28 PDZ domains also identified as significant binders (Genera et al., 2021). Thus, this similar number of PDZ domains recognized by the two PBMS of class I represents about 10% of the human PDZome. PTPN3-PDZ has BI values of 0.52 and 0.45 for PBM-p38γ and PBM-HBc respectively, while PTPN4-PDZ has BI values of 0.60 and 0.58 for the two peptides. This is in agreement with the similar affinities reported previously (Maisonneuve et al., 2016; Genera et al., 2019).

We compared the sequence alignments of PDZ domains recruited by p38γ and HBc PBMs (Figure 6) to the full human PDZome (Figure 3B). We focused on key positions for PDZ/PBM interactions. As expected, the “GLGF motif” at position 520-523 (numbering of PTPN3) in interaction with the hydrophobic residue at P0 of all PBMs is conserved as a signature of PDZ domains in the PDZome and in the pool of PDZ domains recruited by PBM-p38γ (Figure 6A) and by PBM-HBc (Figure 6B). H572 is almost the only residue found at this position with the class I PTPN3’s PBMs PDZ-binding profiles, canonically allowing the interaction with S or T at P-2. N524 is strictly conserved in PTPN3 and PTPN4 orthologs and is also enriched at this position following S in the pools of PDZ domains recruited by PBM-p38γ and PBM-HBc (Figure 6) compared to the whole PDZome where S or T are the most abundant residues (Figure 3B). This is likely related to the H-bond of N524 to E at P-3 found in all reported ligands of PTPN3-PDZ (Figure 4). Similarly, S538 H-bonded to E side chain at P-3 is also favored in the subsets of PDZ domains recruited by PBM-p38γ and PBM-HBc compared to the entire PDZome, despite the good conservation of a serine at this position (Figure 3B). The R or K at P-4 form ionic bonds with D573. D and E are preferred at this position in agreement with their equal ability to form the ionic bond (Figure 6). In the PDZome, E and D are also abundant at this position (Figure 3B). These results agree with a tendency that position -3 and possibly -4 of PTPN3’s PBMs favor specific residues at certain positions in PDZ domains in agreement with the interactions observed in the X-ray structures. These likely contribute to the affinity of the complex and favors the binding to the PDZ domains, as observed with the screening of the human PDZome library.

### 3.6 Investigation of the regulatory elements controlling the catalytic activity of PTPN3

While it is demonstrated that the PDZ domains of PTPN3 and PTPN4 regulates their catalytic activities (Chen et al., 2014; Maisonneuve et al., 2014), the regulatory mechanisms remain poorly understood in PTPN3. Here, we investigated the effects of the PDZ domain, the PBM binding, and the linker connecting the PDZ to PTP (residues 598-628), on the phosphatase activity. We used four constructs of PTPN3: PTPN3-Bidomain, PTPN3-Linker-PTP, PTPN3-shLinker-PTP (short linker, missing 23 residues) and PTPN3-PDZ (Figure 1). In all experiments, the phosphatase activity was assessed using p-nitrophenyl phosphate (pNPP).

We measured and compared the kinetic parameters, the Michaelis constant (K_M_), the turnover number (k_cat_) and the catalytic efficiency (k_cat_/K_M_), of the dephosphorylation reaction catalyzed by PTPN3-Bidomain and PTPN3-linker-PTP (Figure 7A)(Table 3). The k_cat_ of PTPN3-Bidomain is twice lower than the one of PTPN3-linker-PTP (1.5± 0.1 s^-1^ vs 3.0 ± 0.1 s^-1^), whereas the K_M_ values are similar. The PDZ domain inhibits the catalytic activity of PTPN3 as previously reported (Chen et al., 2014).

To assess whether the PBM binding to the PDZ domain releases the catalytic inhibition as observed for PTPN4 (Maisonneuve et al., 2014), we added the PBM peptides of the PTPN3 cellular partners, PBM-TACE and PBM-p38γ, and the viral peptides PBM-16E6, PBM-HBc, in large excess (molar ratio 500:1) to PTPN3-Bidomain (Figure 7A)(Table 3). For all peptides, a 2-fold decrease in k_cat_ compared to PTPN3-linker-PTP is measured, as in the unbound Bidomain, while the K_M_ remained unaffected (Table 3). Thus, we concluded that the PBM binding has no effect on PTPN3 catalytic activity in our conditions. Accordingly, the specificity constants (kcat/K_M_) of the complexed Bidomain are in the same range in comparison with the values of PTPN3-Bidomain (Table 3). Altogether, these data confirm the existence of a PDZ-mediated inhibited state of PTPN3 in the Bidomain construct as reported for PTPN4 (Maisonneuve et al., 2014). However, the binding of PBM ligands of either cellular or viral origin does not affect the PTPN3 regulation of the PTP activity by the PDZ domain in the conditions assayed while we previously reported a partial release of the PTPN4 catalytic inhibition upon PBM binding in similar conditions (Maisonneuve et al., 2014).

Then, we assessed whether the linker that connects the PDZ and PTP domains is required for the catalytic regulation as observed for PTPN4 (Caillet-Saguy et al., 2017). We measured the catalytic activity at a fixed concentration of 2.5 mM pNPP, where the Bidomain and linker-PTP constructs exhibited the highest initial rates of reaction (Figure 7A). We compared the initial rates for PTPN3-Bidomain, for PTPN3-linker-PTP alone, and for PTPN3-linker-PTP with a large excess of PTPN3-PDZ added in *trans* (molar ratio 80:1). We observed similar initial reaction rates for PTPN3-Linker-PTP with and without PTPN3-PDZ added in trans (Figure 7B) whereas, as expected, PTPN3-Bidomain presents a significant lower initial rate. These results indicate that the PDZ-mediated inhibition on PTPN3 catalytic activity requires that the two domains are covalently linked. Additionally, we observed that the PTPN3-shLinker-PTP construct, in which the linker lacks the 23 N-terminal residues (Figure 1), has the same catalytic activity as PTPN3-Linker-PTP, with the full-length linker (Figure 7A)(Table 3). This indicates that the linker alone, freely exposed at the N-terminal has no effect on the catalytic activity. Thus, the PTPN3 regulatory mechanism is *cis*-acting and requires the PDZ domain covalently linked to the PTP domain.

### 3.7 Insights of PTPN3 Bidomain in solution from AUC and SAXS experiments

We conducted AUC and SAXS experiments on the PTPN3 Bidomain to investigate its integrity, oligomeric state, and shape both free and complexed with a PBM in solution. AUC data of PTPN3-Bidomain highlighted a main species at 2.8S with a frictional ratio of 1.6, corresponding to an elongated monomer (Table 4, Figure 8A). In agreement with the AUC experiments, the SAXS experiments on the Bidomain alone or complexed to PBM-p38γ showed that the free Bidomain behaves as a monodisperse distribution of monomers in solution (Table 4)(Figure 8B,C, D). The estimated molecular mass of free Bidomain derived from the extrapolated intensity I(0) at the origin is consistent with the theoretical value of 52.5 kDa. The maximum distance (Dmax) of the protein and the radius of gyration (Rg) derived from the electron pair distance distribution function P(r) (Figure 8C) are similar for the Bidomain free and complexed with PBM-p38γ, indicating that the PBM binding does not alter the overall shape of the Bidomain. Indeed, the Dmax and Rg values are respectively 133 Å and 33 Å for Bidomain alone and 132 Å and 34 Å for Bidomain complexed to PBM-p38γ (Table 4). These results are consistent with the values measured for PTPN4 (Maisonneuve et al., 2014).

**Figure 8.**
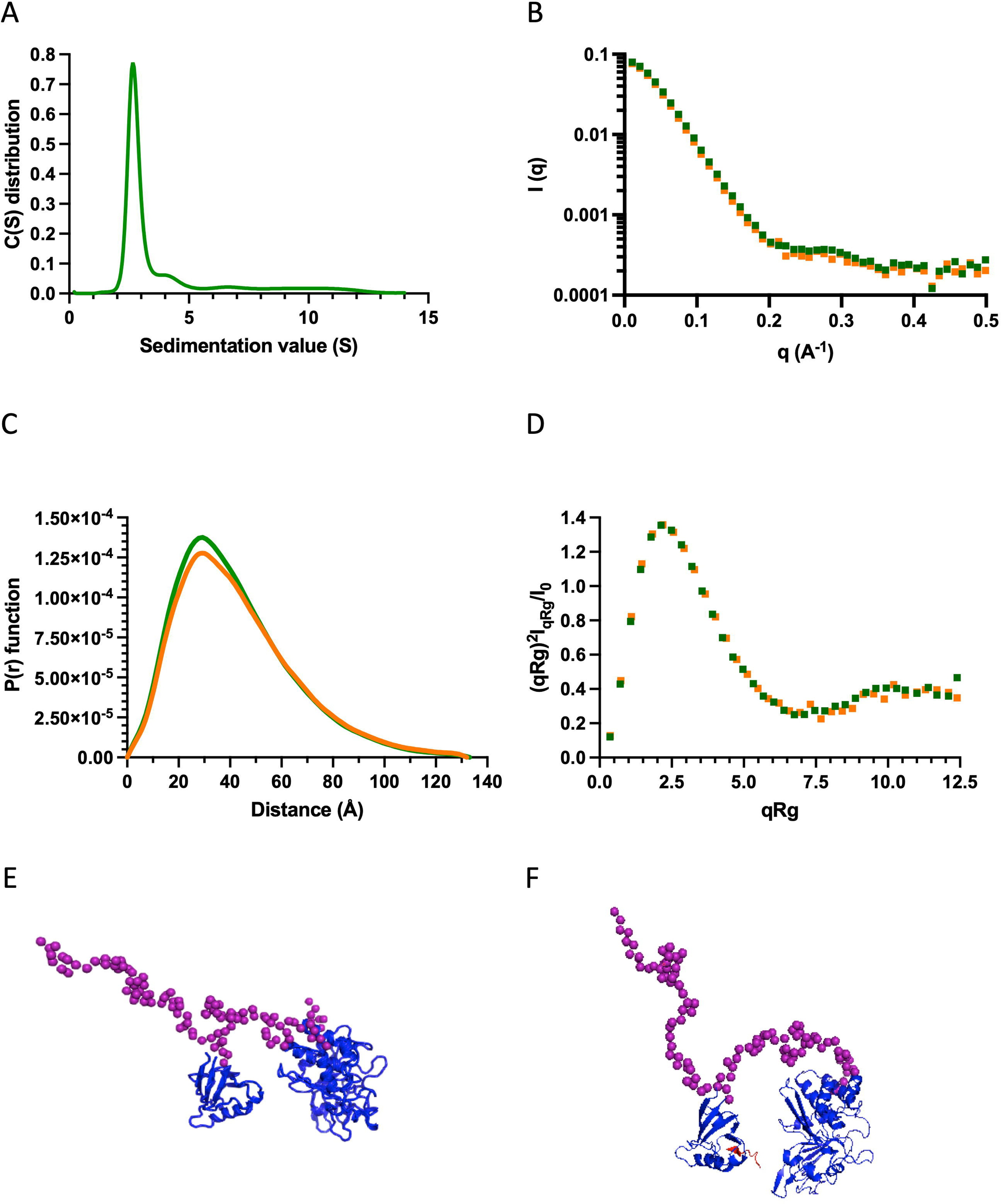
AUC and SAXS analysis of PTPN3 Bidomain. Data of PTPN3 Bidomain WT free and complexed to PBM-p38γ peptide are represented in green and orange, respectively. (A) Sedimentation coefficient distributions of free PTPN3 Bidomain. (B) Experimental SAXS data (I(q) versus q). (C) The P(r) function of the SAXS data, where P is the pair distance distribution function and r is the distance vector. (D) Dimensionless Kratky plots. (E)(F) Models of Bidomain obtained from SAXS data. Models were generated using CORAL based on the X-ray structures of the PDZ (PDB code 6T36) and the catalytic domain (PDB code 2B49) of PTPN3. The two domains were rigid and the linker, the N-terminal and the C-terminal part of the PTPN3 were set in random conformations. 50 models were generated, the best model is presented. The model of PTPN3 free (E) and in complex with PBM-p38γ (F) were generated comparably including the PBM (in red) within the X-ray structure of the PDZ. Known structures are in blue. The interdomain linker, the N- and C-terminal parts of the bidomain are shown as purple spheres.

**Table 4.** Hydrodynamic parameters of PTPN3-Bidomain derived from the analysis of AUC and SAXS data.

| | Hydrodynamic parameters | PTPN3-Bidomain/ PTPN3-Bidomain+ PBM-p38γ |
| --- | --- | --- |
| AUC | Mass theoretical (kDa) | 52.5 |
|  | S<sub>0,w,20</sub> (S) | 2.8 |
|  | f/f<sub>0</sub> | 1.6 |
| SAXS | R<sub>g</sub> (Å) | 33/34 |
|  | D<sub>max</sub> (Å) | 133/132 |

The normalized sedimentation coefficients (S<sub>0</sub>) were processed to get the standard sedimentation coefficients in water at 20 °C (S<sub>0,w,20</sub>).

We used the SAXS intensity profile to model the 3D arrangement of both domains using the known structures of the PDZ domain and the catalytic domain (Figure 8E). The models were comparable with and without the ligand, illustrating that the PBM binding does not affect the overall arrangement of both domains of the Bidomain in solution that remain extended in solution in agreement with the frictional ratio measured.

## 4 Discussion

Several viral and cellular proteins target the PDZ domain of PTPN3 through PBMs, potentially affecting the function of PTPN3.

We focused on studying the molecular basis of the selectivity of PTPN3 recognition for such short and unstructured PBMs. We characterized the determinants of PDZ ligand recognition of PTPN3 by solving the crystal structures of PTPN3-PDZ complexed to PBM peptides derived from viral and cellular PTPN3 partners. Indeed, we provided two structures of PTPN3-PDZ in complex with the PBMs of HPV18 and TACE. We compared them with our previous reported strutures of PTPN3-PDZ in complex with viral PBMs from HPV16 and HBV (Genera et al., 2019, 2021). We found that all these PBMs establish a similar binding pattern involving conserved residues in the PDZ domains of the two homologs PTPN3 and PTPN4, whereas these residues are less often conserved in the PDZome. We propose that they could therefore represent determinants of the sequence preference of PBM ligand for the NT5 phosphatase subfamily.

Interestingly, the binding of PBMs exposing a C-terminal cysteine has only been reported in a handful of cases, many of which bind to the PDZ domain of GIPC1, interacting with the PBMs of HBc (Razanskas and Sasnauskas, 2010), the lutropinchoriogonadotropic hormone receptor (Hirakawa et al., 2003), the complement component C1q receptor (Bohlson et al., 2005), the insulin-like growth factor 1 receptor (Ligensa et al., 2001), and the dopamine 2 and 3 receptors. These latter share the atypical C-terminal sequence –KILHC_COOH_ (Jeanneteau et al., 2004). Other PDZ domains, MAGI-1 PDZ5, the Scrib and PDLIM-4 PDZ domains, bind also to PBM with a C-terminal cysteine (Cuppen et al., 2000; Petit et al., 2005; Chastre et al., 2009). There is a lack of structural data on complexes involving this type of PBM. To our knowledge, only two structures are available: the one of PDZK1 PDZ1 domain complexed with the C-terminal PBM of the prostacyclin receptor (-ACSLC_COOH_)(Birrane et al., 2013) and the one of GRIP1 PDZ6 domain complexed to the liprin-α PBM (-RTYSC_COOH_)(Im et al., 2003). We observed that the cysteine side chain has the same orientation for GRIP1 PDZ6, where the cysteine can be accommodated towards the interior of the hydrophobic carboxylate-binding pocket despite the polar character of the thiol group. It is widely reported that the hydrophobic character of the side chains that compose the carboxylate binding-pocket imposes a specificity requirement for PBMs containing hydrophobic residues in the C-terminal position (Sheng and Sala, 2001). The preference of some hydrophobic residues over others at P0 has been attributed to variations in the size and the geometry of the hydrophobic carboxylate-binding pocket (Songyang et al., 1997). However, the selectivity at P0 is not highly stringent as observed for PTPN3, whose PDZ domain can accommodate different hydrophobic residues at P0. Nonetheless, it is remarkable that both GIPC1 and PTPN3 have multiple C-terminal cysteine PBM partners, while this residue is not frequently found at P0 (less than 8.5% in all C-terminal human PBMs; Figure S1; Supplementary Materials 7 and 8). Interestingly, the PBM of RGS-GAIP containing an alanine at P0 (13.8% in all C-terminal human PBMs) was reported to interact with the PDZ of GIPC1 (De Vries et al., 1998). It is not surprising that the PDZ of GIPC can accommodate an alanine since both alanine and cysteine are of similar size.

We used proteomics data to explore the sequence consensus of the cellular partners bound to PTPN3-PDZ, highlighting conserved motifs consistent with the observations derived from the crystal structures. Some of the conserved positions in PTPN3 and PTPN4 PDZ domains mediate contacts beyond the canonical 3-residue PBM that are not required for the interactions to occur, but possibly favor the selection of some PBM partners over others by increasing their interaction affinities. PDZ domains are promiscuous protein-protein interaction modules that bind to their partners with low-to-medium affinities (1-100 μM), which is related to the transient nature of signaling interactions. The specific polar bonds and hydrophobic contacts that the preferred PTPN3 ligands establish via their positions -3 and -4 are likely to enhance their binding over other potential PBM-containing partners. To go further on the selectivity of PTPN3-PDZ, we reported the PBM sequence analysis on the interactome of PTPN3-PDZ. We documented the PBMs sequences that were selectively captured from a cell lysate by PTPN3-PDZ and identified a motif that is favored by PTPN3-PDZ in this context with E or S at P-1, and a preference for E at P-3 (Figure 5). No preferential residue at P-4 was observed. However, knowing that using the pull-down methodology we fished full-length proteins, so we cannot exclude that the interactions occur through a different interaction motif, for example via internal PBMs, that we cannot identify. Additionally, in a growing number of complexed PDZ domain structures (Kang et al., 2003; Sugi et al., 2007; Elkins et al., 2010), class II PBMs peptides are inserted perpendicular to the PDZ domain, with only position 0 and sometimes position -1 interacting with the PDZ domain. Thus, it is possible that class II PBMs interact with PTPN3-PDZ through a non-canonical binding mode. Although it is possible that this perpendicular binding is solely an artifact of the crystal packing, the observation of this binding mode by NMR for the autoinhibited X11α PDZ1 domain (Long et al., 2005) suggests that this type of non-canonical binding could be relevant in solution.

PDZ domains are a common structural domain found in many proteins and play a crucial role in mediating protein-protein interactions. They are often located in conjunction with other catalytic or non-catalytic domains and contribute to the overall function and regulation of the protein. It is likely that a further degree of selectivity is achieved thanks to the spatial segregation of the protein by the PTPN3 FERM domain, which targets the phosphatase to the interface of the membrane and the cytoskeleton, promoting interaction with certain ligands or substrates over others. This is supported by the observation that both the FERM and PTP domains of PTPN3 are required for attenuation of HBV genome expression (Hsu et al., 2007). Interestingly, two of the three isoforms of PTPN3 that have been described are likely to lack this spatial segregation due to their truncated FERM domains. These isoforms are likely to be more active than full-length PTPN3, as suggested by *in vitro* limited proteolysis studies (Zhang et al., 1995). Unfortunately, to the best of our knowledge, there is currently no data on the subcellular location or the physiological relevance of these isoforms. One can only hypothesize about their potential role, and the relevance of their PDZ domains for selecting substrates or anchoring these enzymes to signaling complexes. PTPN3, for example is able to specifically dephosphorylate the MAPK p38γ thanks to the recognition by its PDZ domain of the C-terminal PBM of p38γ (Hou et al., 2010). There is an increasing awareness that non-catalytic scaffold domains can perform direct regulatory functions on the catalytic domain to which they are linked, exceeding their established roles as inert binding domains.

In this work, we performed and analyzed the kinetics of the phosphatase activity of PTPN3 in the context of the isolated PTP domain and the PDZ-PTP bidomain construct. The PDZ domain inhibits the activity of the adjacent PTP domain by decreasing the turnover number, without affecting the affinity for the substrate. This indicates that the PDZ domain is not blocking the accessibility of the phosphatase substrate to the PTP active site. Therefore, the inhibition is non-competitive, as found for PTPN4 (Maisonneuve et al., 2014). We were interested in exploring whether PTPN3 features a similar allosteric regulatory mechanism as PTPN4. We showed that the linker of PTPN3 is necessary for the inhibition as the one of PTPN4 (Maisonneuve et al., 2014; Caillet-Saguy et al., 2017). The binding of a PBM to PTPN4 releases the inhibition whereas the PBM binding to PTPN3-PDZ does not affect the catalytic regulation.

PTPN3 and PTPN4 share 51% of global sequence identity, but this rises to 71% for their PDZ domains and 61% for their catalytic domains. We have previously shown that a conserved hydrophobic FQYI sequence (residues 620-623 in PTPN4) in the PDZ-PTP linker in PTPN4 is implicated in the regulation (Caillet-Saguy et al., 2017; Spill et al., 2021). However, these residues at these positions are not strongly conserved between PTPN3 and PTPN4 (Figure S5). We can hypothesize that this patch is strictly conserved in orthologous PTPN4 (Figure S5C) to allow the regulation upon PBM binding. The linker (34-residue long) in PTPN3 is predicted mostly unstructured by Alphafold (https://alphafold.ebi.ac.uk/entry/P26045) (Jumper et al., 2021; Varadi et al., 2022) as the one of PTPN4 (https://alphafold.ebi.ac.uk/entry/P29074) which was experimentally validated by NMR in solution (Maisonneuve et al., 2014; Caillet-Saguy et al., 2017). However, the linker of PTPN4 is yet resistant to proteolysis, which could support an interaction, most likely transient, with the PTP domain, as previously proposed (Caillet-Saguy et al., 2017; Spill et al., 2021). On the contrary, the linker of PTPN3-Bidomain is sensitive to *in vitro* proteolysis even in the presence of protease inhibitors (Figure S6), indicating that it is most likely predominantly exposed and possibly having little or no interaction with the PTP domain. Multidomain proteins frequently employ intrinsically disordered regions for the purpose of allosteric regulation (Huang et al., 2020). Linkers between domains have been shown to enhance the local concentration of domains and enable allosteric regulation of weakly interacting partners, resulting in a rather complex allosteric mechanism and novel protein behavior (Huang et al., 2020). NMR mapping of the chemical shift changes that occur in PTPN3-PDZ and PTPN4-PDZ upon ligand binding showed long-range structural and dynamics perturbations (Babault et al., 2011; Genera et al., 2019). Studying the dynamics of free and PBM-bound PTPN3-Bidomain by NMR would provide information about any structural rearrangements that might occur upon PDZ ligand binding. Unfortunately, the proteolysis of the linker prevented us to record usable HSQC spectra. A well-documented case is the regulation of the catalytic activity of the tyrosine phosphatase SHP-2 by its two SH2 adjacent domains (Hof et al., 1998). The WPD loop, which is located near the active site of SHP-2, contains a conserved aspartic acid residue that plays a critical role in the dephosphorylation catalytic activity. The conformational changes of the WPD loop are also important for substrate recognition and catalysis. The unbound N-terminal SH2 domain of SHP-2 interacts with the phosphatase domain, sterically blocking the active site in an open but inactive conformation, preventing the closure of the WPD loop resulting in competitive inhibition. The binding of a phosphoprotein ligand to the SH2 domain triggers allosteric conformational rearrangements that prevent binding of the complexed SH2 to the PTP domain, releasing the inhibition. The SH2 domain thus works as an allosteric molecular switch. Similarly, the PDZ domains of PTPN4 and PTPN3 lock the phosphatase domain in an auto-inhibited conformation, and the catalytically active state is restored upon binding of a PBM for PTPN4 (Maisonneuve et al., 2014) but not for PTPN3. Both the non-competitive inhibition and release of inhibition processes occur through long-range intramolecular allosteric mechanisms that require the covalent binding of the two domains. A modulation of the WPD loop through the linker was recently proposed for the PTPN4 inhibition (Spill et al., 2021). We also hypothesize such molecular mechanism for PTPN3 and suggest that the variability of the hydrophobic patch observed in the linker for PTPN3 (Figure S5B) could explain the absence of release of inhibition by the PBM.

## 6 Author Contributions

MG and CCS contributed to the conception and design of the study. MG and BR performed and analyzed the AUC and SAXS experiments. MG, AC and CCS performed and analyzed the NMR experiments. MG, BCC and CCS performed sequence analysis. MG, AM, AH, CCS performed and analyzed the X-ray crystallography experiments. MG and AC performed and analyzed the kinetics experiments. MG and CCS performed and analyzed the holdup experiment. JC contributed to the cloning of constructs. MG and AC contributed to protein production. MG, BCC, NW, and CCS wrote the first draft of the manuscript. All authors contributed to manuscript revision, read, and approved the submitted version.

## 8 Acknowledgments

We thank the staff of the crystallography platform at Institut Pasteur for carrying out robot-driven crystallization screening. We thank the Institut Pasteur molecular biophysics platform for the technical help and access to instruments. We thank the Institut Pasteur biological NMR technological platform for access to instruments and help with NMR experiments. We acknowledge synchrotron SOLEIL (Saint-Aubin, France) for granting access to their facility and the staff of Proxima1 for helpful assistance during the data collection and Aurelien thureau and Javier Perez for the SWING beamline.

## 9 Data Availability Statement

The original contributions presented in the study are included in the article/Supplementary Material, further inquiries can be directed to the corresponding author.

