## Supplementary_Material for "Interactions of the Protein Tyrosine Phosphatase PTPN3 with Viral and Cellular Partners through its PDZ Domain: Insights into Structural Determinants and Phosphatase Activity"

### 1 Supplementary Data

Supplementary Material 1. List of all raw BI values obtained by holdup assay using PBM peptide from p38γ protein. Based on BI values higher than 0.2 for significant PDZ–peptide interactions (Vincentelli et al., 2015), 28 PDZ domains were identified.

| Protein name_PDZ number | Binding Intensity (BI) | PDZ name_PDZ number | BI |
|---|---|---|---|
| NHERF3_1 | 0.88 | SYNPO2L_1 | 0 |
| SCRIB_3 | 0.75 | PPP1R9B_1 | 0 |
| MAST2_1 | 0.72 | PDZD7_2 | 0 |
| SNX27_1 | 0.60 | TJP1_3 | 0 |
| PTPN4_1 | 0.60 | PTPN13_3 | 0 |
| MAST1_1 | 0.55 | CASK_1 | 0 |
| PTPN3_1 | 0.52 | MAGI1_5 | 0 |
| PDZRN3_1 | 0.51 | RHPN2_1 | 0 |
| SYNJ2BP_1 | 0.49 | PARD3_1 | 0 |
| SCRIB_1 | 0.49 | MAGI2_4 | 0 |
| SNTB1_1 | 0.47 | PDZD2_3 | 0 |
| SNTB2_1 | 0.41 | MPDZ_9 | 0 |
| PDZRN4_1 | 0.37 | MPDZ_3 | 0 |
| DLG1_2 | 0.36 | APBA3_2 | 0 |
| NHERF2_2 | 0.35 | GRIP1_5 | 0 |
| DLG4_2 | 0.33 | DLG5_2 | 0 |
| FRMPD4_1 | 0.31 | TJP3_2 | 0 |
| LAP2_1 | 0.30 | RIMS2_1 | 0 |
| SNTA1_1 | 0.29 | APBA3_1 | 0 |
| TX1B3_1 | 0.27 | PDZD2_1 | 0 |
| DLG2_2 | 0.27 | PTPN13_1 | 0 |
| RGS12_1 | 0.25 | MPDZ_8 | 0 |
| DFNB31_1 | 0.25 | PREX2_1 | 0 |
| DLG3_2 | 0.24 | PDZRN3_2 | 0 |
| SHANK1_1 | 0.23 | MPDZ_12 | 0 |

| | | | |
|---|---|---|---|
| PARD6A_1 | 0.23 | PDZD4_1 | 0 |
| DLG4_3 | 0.21 | HTRA1_1 | 0 |
| MAGI1_2 | 0.20 | MPP7_1 | 0 |
| MAGI3_2 | 0.20 | CARD14_1 | 0 |
| PARD3B_3 | 0.20 | PRX_1 | 0 |
| ARHGEF12_1 | 0.19 | GRIP1_2 | 0 |
| MPDZ_7 | 0.19 | ARHGEF11_1 | 0 |
| NHERF4_3 | 0.19 | PDLIM2_1 | 0 |
| DLG2_1 | 0.18 | LNX2_3 | 0 |
| DLG1_1 | 0.17 | PARD3_2 | 0 |
| PDZD2_6 | 0.16 | TJP3_3 | 0 |
| IL16_1 | 0.16 | PARD6G_1 | 0 |
| NHERF2_1 | 0.15 | MAGI3_1 | 0 |
| SHANK3_1 | 0.15 | LNX2_2 | 0 |
| MAGI3_5 | 0.15 | DFNB31_3 | 0 |
| NHERF3_2 | 0.14 | MPDZ_6 | 0 |
| InaDl_3 | 0.13 | SDCBP_2 | 0 |
| SNTG2_1 | 0.12 | PSCDBP_1 | 0 |
| MAGI2_6 | 0.12 | PARD6B_1 | 0 |
| HTRA3_1 | 0.12 | SDCBP2_2 | 0 |
| LNX2_4 | 0.12 | IL16_3 | 0 |
| SHROOM3_1 | 0.11 | MPDZ_2 | 0 |
| InaDl_4 | 0.11 | APBA1_1 | 0 |
| MLLT4_1 | 0.10 | PREX2_2 | 0 |
| APBA1_2 | 0.10 | PDLIM7_1 | 0 |
| SHANK2_1 | 0.09 | GORASP1_1 | 0 |
| SHROOM2_1 | 0.09 | CARD11_1 | 0 |
| HTRA4_1 | 0.09 | SIPA1L1_1 | 0 |
| PTPN13_2 | 0.09 | FRMPD3_1 | 0 |
| ARHGAP21_1 | 0.09 | LIMK2_1 | 0 |
| GOPC_1 | 0.09 | TJP3_1 | 0 |
| SIPA1L3_1 | 0.09 | LNX1_3 | 0 |
| RHPN1_1 | 0.09 | PPP1R9A_1 | 0 |
| PDZD7_3 | 0.09 | MAGI2_1 | 0 |
| SIPA1_1 | 0.08 | FRMPD2_3 | 0 |
| NHERF4_4 | 0.08 | TIAM1_1 | 0 |
| PDZD9_1 | 0.07 | MAGI1_1 | 0 |
| DLG1_3 | 0.06 | PCLO_1 | 0 |
| SHROOM4_1 | 0.06 | PDLIM4_1 | 0 |
| DFNB31_2 | 0.06 | ARHGAP23_1 | 0 |
| DVL3_1 | 0.06 | GRIP1_4 | 0 |
| DLG4_1 | 0.06 | GRIP2_3 | 0 |
| MPP5_1 | 0.06 | GORASP2_1 | 0 |
| FRMPD2_1 | 0.06 | MPDZ_4 | 0 |
| InaDl_2 | 0.05 | SDCBP_1 | 0 |
| FRMPD1_1 | 0.05 | LIN7A_1 | 0 |

| | | | |
|---|---|---|---|
| MAGI1_6 | 0.05 | PTPN13_5 | 0 |
| MPDZ_11 | 0.05 | GRIP1_7 | 0 |
| PREX1_1 | 0.05 | LDB3_1 | 0 |
| TIAM2_1 | 0.05 | FRMPD2_2 | 0 |
| DLG3_3 | 0.05 | LNX1_1 | 0 |
| MPP4_1 | 0.05 | MPP1_1 | 0 |
| GRIP1_1 | 0.05 | DLG5_4 | 0 |
| MAGI2_2 | 0.05 | InaDl_8 | 0 |
| MPDZ_13 | 0.05 | SCRIB_2 | 0 |
| TJP1_1 | 0.04 | GRIP1_6 | 0 |
| NHERF3_4 | 0.04 | SYNP2_1 | 0 |
| InaDl_6 | 0.04 | NHERF4_2 | 0 |
| AHNAK1_1 | 0.04 | PICK1_1 | 0 |
| SNTG1_1 | 0.03 | GRIP2_7 | 0 |
| MAGI3_4 | 0.03 | LRRC7_1 | 0 |
| MAGI2_5 | 0.03 | RGS3_1 | 0 |
| PARD3_3 | 0.03 | InaDl_1 | 0 |
| MAGI3_6 | 0.03 | GRID2IP_1 | 0 |
| MPP6_1 | 0.03 | TJP2_3 | 0 |
| PSMD9_1 | 0.03 | InaDl_5 | 0 |
| NHERF_2 | 0.03 | MAST4_1 | 0 |
| MYO18A_1 | 0.03 | USH1C_1 | 0 |
| LMO7_1 | 0.03 | GRIP2_5 | 0 |
| LNX1_4 | 0.02 | IL16_2 | 0 |
| NOS1_1 | 0.02 | GRIP2_4 | 0 |
| DLG5_1 | 0.02 | CNKSR2_1 | 0 |
| RIMS1_1 | 0.02 | DVL1_1 | 0 |
| MAGI1_3 | 0.02 | IL16_4 | 0 |
| DLG3_1 | 0.02 | GRIP2_6 | 0 |
| GRASP_1 | 0.02 | MPP2_1 | 0 |
| NHERF3_3 | 0.02 | AHNAK2_1 | 0 |
| PREX1_2 | 0.02 | InaDl_7 | 0 |
| MAGI2_3 | 0.02 | RAPGEF6_1 | 0 |
| SDCBP2_1 | 0.01 | RAPGEF2_1 | 0 |
| SIPA1L2_1 | 0.01 | GRIP1_3 | 0 |
| MAGIX_1 | 0.01 | DEPTOR_1 | 0 |
| PARD3B_2 | 0.01 | TJP2_2 | 0 |
| MAGI1_4 | 0.01 | USH1C_3 | 0 |
| GIPC2_1 | 0.01 | PDZD7_1 | 0 |
| CNKSR1_1 | 0.01 | TJP2_1 | 0 |
| NHERF_1 | 0 | LIN7C_1 | 0 |
| PDZD2_2 | 0 | MAGI3_3 | 0 |
| PDLIM1_1 | 0 | MPDZ_5 | 0 |
| PDZD8_1 | 0 | DVL2_1 | 0 |
| LNX2_1 | 0 | LNX1_2 | 0 |
| PDLIM3_1 | 0 | LIMK1_1 | 0 |
| SCRIB_4 | 0 | PDLIM5_1 | 0 |

| | | | |
|---|---|---|---|
| STXB4_1 | 0 | InaDl_9 | 0 |
| PARD3B_1 | 0 | GRIP2_1 | 0 |
| MPP3_1 | 0 | InaDl_10 | 0 |
| PDZRN4_2 | 0 | HTRA2_1 | 0 |
| MPDZ_10 | 0 | GIPC1_1 | 0 |
| DLG5_3 | 0 | MPDZ_1 | 0 |
| TJP1_2 | 0 | APBA2_1 | ND |
| NHERF4_1 | 0 | APBA2_2 | ND |
| MAST3_1 | 0 | CNKSR3_1 | ND |
| RADIL_1 | 0 | DLG2_3 | ND |
| PDZD11_1 | 0 | GIPC3_1 | ND |
| LIN7B_1 | 0 | GRID2IP_2 | ND |
| PTPN13_4 | 0 | GRIP2_2 | ND |
| | | INTU_1 | ND |

Supplementary Material 2. Table of SAXS collection parameters.

| | |
|---|---|
| Data collection parameters | |
| Instrument | SWING beamline, Synchrotron Soleil |
| Beam geometry | KB mirrors + guard slits |
| Wavelength (Å) | 1.033 |
| *q* range ($Å^{-1}$) | 0.0006–0.5000 |
| Exposure time (s) | 1s per frame |
| Temperature (K) | 290 |
| Structural parameters | |
| *I*(0) ($cm^{-1}$) [from *P*(*r*)] | 0.8317 E-01[a]; 0.7992 E-01[b] |
| $R_g$ (Å) [from *P*(*r*)] | 33[a]; 34[b] |

[a]Bidomain alone ; [b]Bidomain complexed to PBM-p38γ

Supplementary Material 3. List of 83 proteins presenting a C-terminal PBM previously identified by LC-MS/MS after pull-down experiments using PTPN3-PDZ as bait with HeLa S3 cell lysates (Genera et al., 2021). Uniprot code and C-terminal sequences from P-4 to P0 are provided in a fasta format.

Supplementary Material 4. List of 34 proteins presenting a class I C-terminal PBM previously identified by LC-MS/MS after pull-down experiments using PTPN3-PDZ as bait with HeLa S3 cell lysates (Genera et al., 2021). Uniprot code and C-terminal sequences from P-4 to P0 are provided in a fasta format.

Supplementary Material 5. List of 33 proteins presenting a class II C-terminal PBM previously identified by LC-MS/MS after pull-down experiments using PTPN3-PDZ as bait with HeLa S3 cell lysates (Genera et al., 2021). Uniprot code and C-terminal sequences from P-4 to P0 are provided in a fasta format.

Supplementary Material 6. List of 16 proteins presenting a class III C-terminal PBM previously identified by LC-MS/MS after pull-down experiments using PTPN3-PDZ as bait with HeLa S3 cell lysates (Genera et al., 2021). Uniprot code and C-terminal sequences from P-4 to P0 are provided in a fasta format.

Supplementary Material 7. List of human proteins presenting a C-terminal PBM. Uniprot code and C-terminal sequences from P-4 to P0 are provided in a fasta format.

Supplementary Material 8. Frequency of residues at the P0 of the C-terminal PBMs of all the human PBM-containing proteins.

```
% residue per position
position 0 :
A(13.7622%)
C(8.50162%)
F(13.089%)
V(19.3219%)
I(12.8646%)
L(32.4607%)
```

### 2 Supplementary Figures

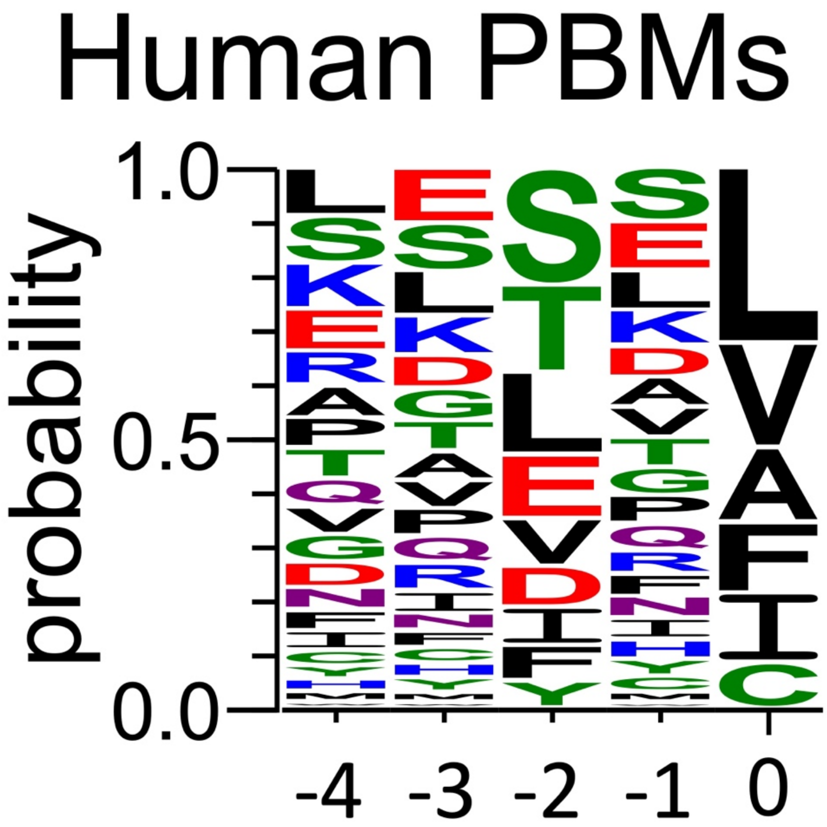


**Figure S1. Frequency plot of the C-terminal residues of all the human PBM-containing proteins.**

For each extension, all UniprotKB sequences were aligned (Supplementary Material 7). The result is displayed using a logo representation (Crooks et al., 2004), where the height of each residue one-letter code translates to its conservation at the corresponding position in the sequence alignment. Amino acids are coloured according to their chemical properties: polar amino acids (G,S,T,Y,C,Q,N) are green, basic (K,R,H) blue, acidic (D,E) red and hydrophobic (A,V,L,I,P,W,F,M) amino acids are black. The positions of the PBM are indicated above the sequences.

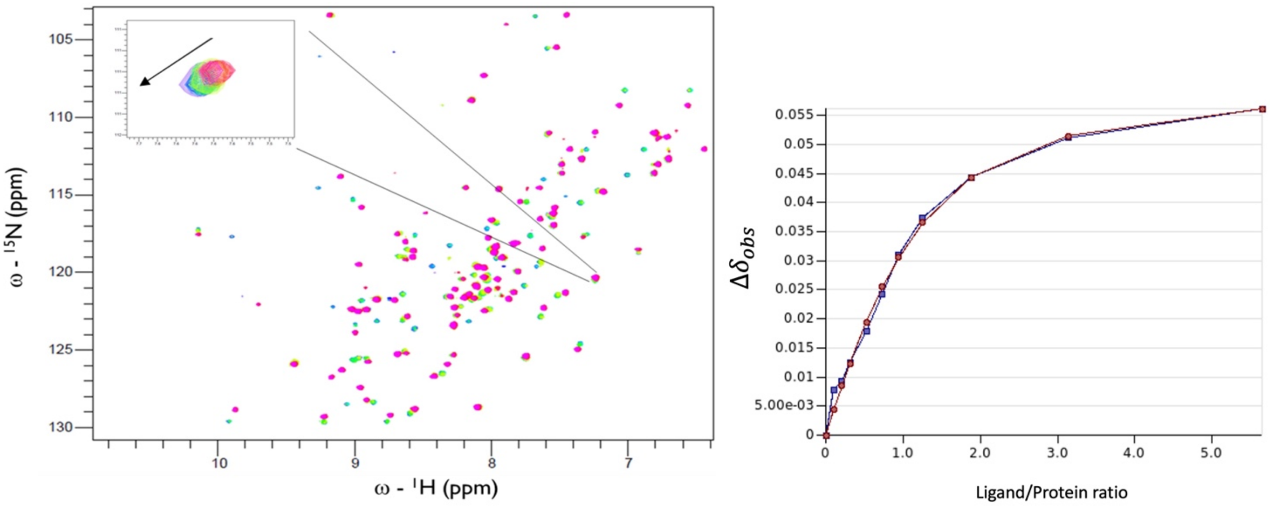


**Figure S2. NMR titration of PDZ-PTPN3 with TACE peptide.**

A) 1H-15N HSQC Spectra of PDZ-PTPN3 titration with TACE peptide. Experiments were performed at 20°C with protein-ligand ratio (mol:mol) starting from 1:0 to 1:5.6 (11 experiments). B) Exemple of titration following one peak in blue and the related fit in red used in CCPNmr Analysis (Vranken et al., 2005).

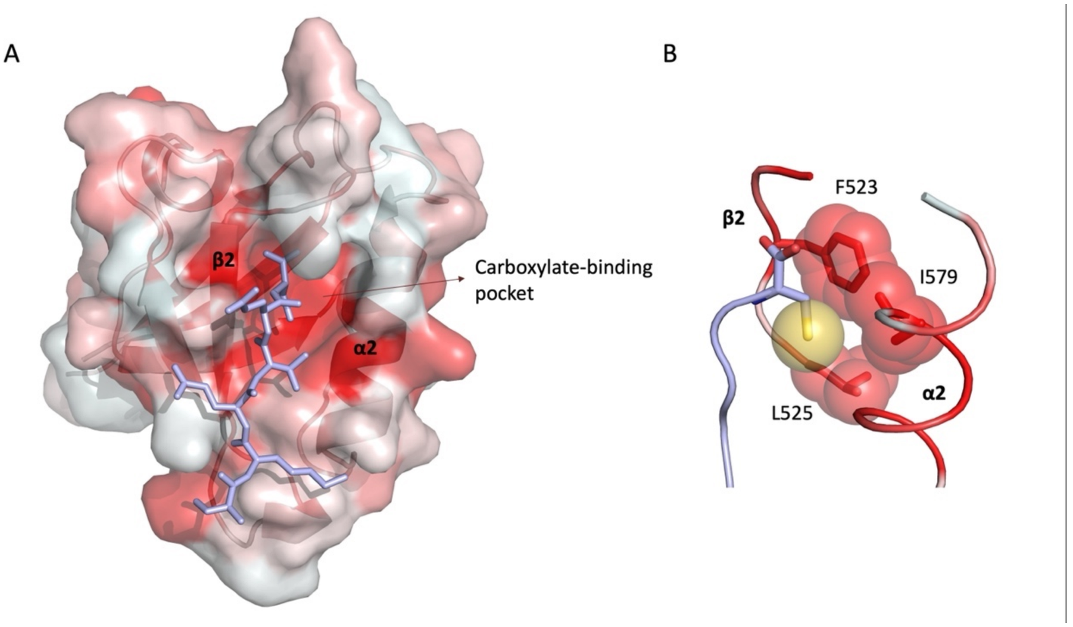


**Figure S3. Surface hydrophobicity of PTPN3-PDZ.**

A) The surface of PTPN3-PDZ is color-coded according to the normalized consensus hydrophobicity scale346. Darker red indicates a more hydrophobic environment, while white indicates polar regions. The peptide shown is PBM-TACE. B) Detail of the position of the terminal cysteine of PBM-TACE in the carboxylate-binding pocket of PTPN3-PDZ. The spheres represent the Van der Waals radii of the sulphur (yellow) and carbon (red) atoms.

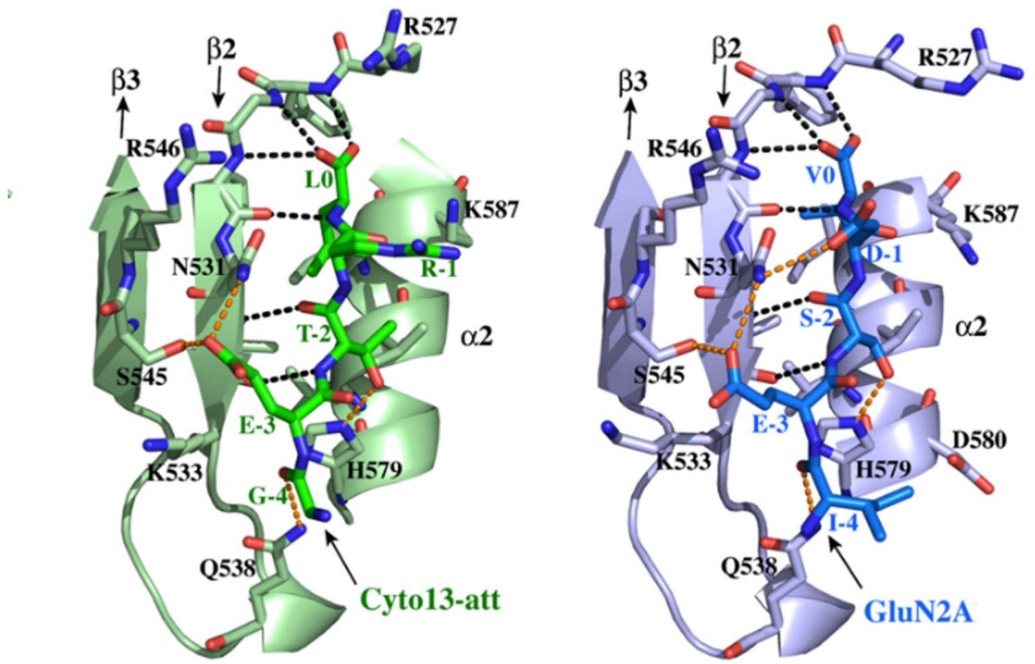


**Figure S4. Insights of the binding network of PTPN4-PDZ with the PBM peptides of Cyto13-att and GluN2A-16.**

Important residues are shown as sticks in CPK colors. Peptides form an antiparallel β sheet with the β2 strand via a complete set of intermolecular canonical backbone H-bonds (black dashed lines). Other intermolecular H-bonds are shown in orange dashed lines.

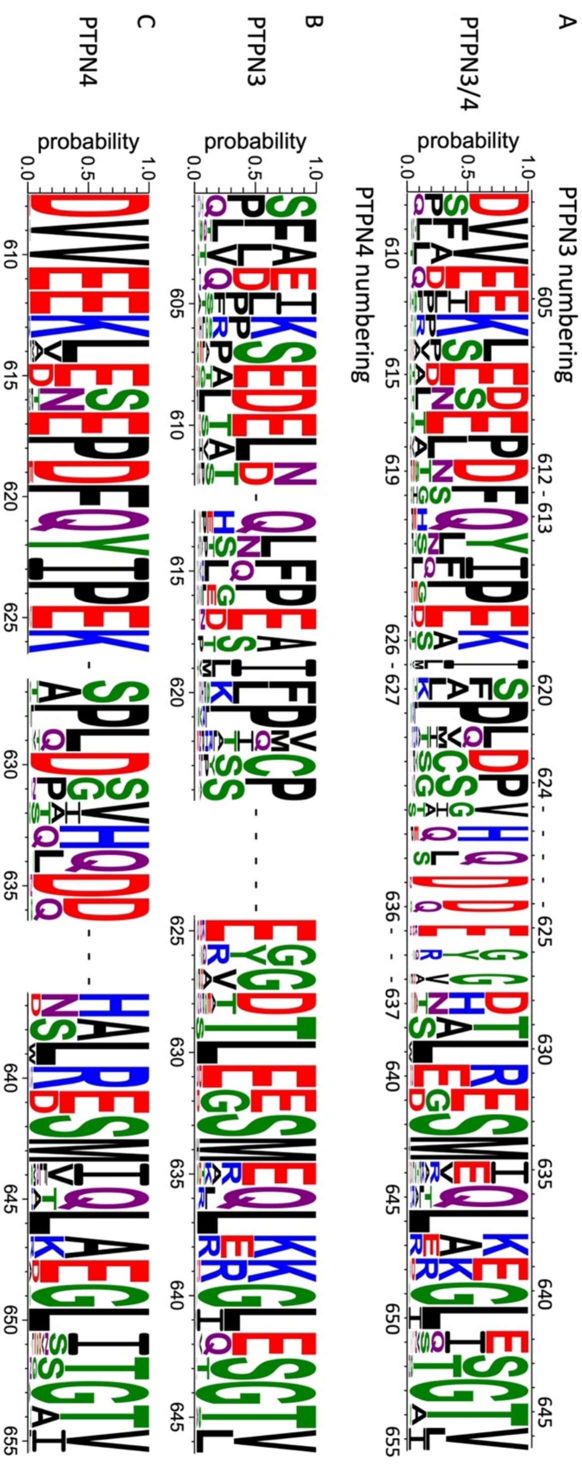


**Figure S5. Comparison of the sequence conservations of the linker between PDZ and PTP domains in PTPN3 and PTPN4 orthologs.**

The result is displayed using a logo representation (Crooks et al., 2004), where the height of each residue one-letter code translates to its conservation at the corresponding position of the PTPN3 or PTPN4 numbering in the sequence alignment. Sequence logo showing the sequence conservation of interdomain linker of A) PTPN3 and PTPN4 orthologs, B) only PTPN3 and C) only PTPN4 orthologs.

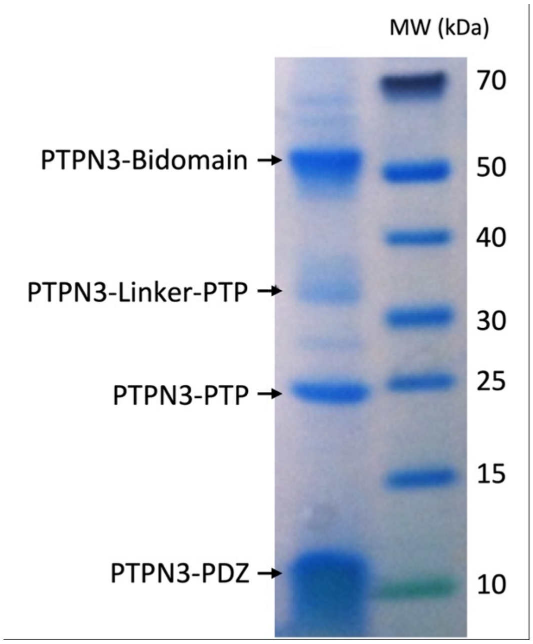


**Figure S6. Sensitivity of PTPN3-Bidomain to proteolysis.**

PTPN3-Bidomain was purified in 50 mM Tris-HCl pH 7.5, 150 mM NaCl, 0.5 mM TCEP and cOmplete protease inhibitor cocktail (Roche) and kept at 25°C for 15h. The state of the sample was then checked by SDS-PAGE, showing that a large proportion of the protein had been cleaved at the interdomain linker.
