## Supplementary_Material 3 for "Interactions of the Protein Tyrosine Phosphatase PTPN3 with Viral and Cellular Partners through its PDZ Domain: Insights into Structural Determinants and Phosphatase Activity"

>Q9Y2T3
FSSSV
>P49189
VESAF
>Q9HB71
GDTEF
>Q15046
VGTSV
>O43707
GESDL
>O95336
KHSTL
>P20020
LETSL
>P12814
GESDL
>P23634
LETSV
>P55060
SVTLL
>P27816
QETSI
>P53985
EESPV
>Q96BN8
EETSL
>P35354
RSTEL
>P48960
SESGI
>P52569
KTSEF
>P62829
AGSIA
>Q96TA1
VQTEF
>Q9Y6M5
PESSL
>O15427
PETSV
>P23219
PSTEL
>P28331
EPSIC
>P81605
LDSVL
>O14734
SESKL
>P09543
SCTII
>P11169
TTTNV
>P26599
SKSTI

>Q13151
GGSSF
>Q5JVF3
LSTVC
>Q8TAA9
SETSV
>Q8TBC3
NETSF
>Q9Y6M7
AETSL
>O00592
EDTHL
>Q02543
PNTFF
>P25705
AGFEA
>P08195
FPYAA
>P21796
LEFQA
>P45880
LELEA
>Q9P0L0
GKFIL
>Q9Y277
FELEA
>P35610
CRYVF
>P51571
SHIQA
>P62081
PEFQL
>O00303
KLVNL
>P46778
YEFMA
>O14975
KTLKL
>O95292
GKIAL
>P16070
MKIGV
>P61619
GALLF
>Q92973
AFYGV
>O75533
LDYIL
>O75964
IGYDV
>O95399
WKYCV
>P00403
PVFTL

>P31930
FWLRF
>P62277
SALVA
>Q59GN2
TKLGL
>Q969Q0
QVIQF
>Q05193
PPFDL
>Q13347
FEFEA
>Q5JTV8
RGICL
>Q7Z2K6
DLFVF
>Q96P70
QTIGI
>Q9NZ01
IPFLL
>Q9P2X0
RGLRF
>Q9UMS4
KFYSL
>O00743
TPYFL
>P04843
ILDAL
>P11021
EKDEL
>Q3ZCM7
EEEVA
>Q15084
GKDEL
>P13010
LLDMI
>P22695
FVDEL
>P14625
EKDEL
>P20674
GLDKV
>P62826
EDDDL
>P48047
MREIV
>Q71DI3
RGERA
>O75947
PIENL
>P13639
FLDKL
>P27797
AKDEL

>Q14257
YHDEL
>P14314
DHDEL
