## Supplementary_Material 7 for "Interactions of the Protein Tyrosine Phosphatase PTPN3 with Viral and Cellular Partners through its PDZ Domain: Insights into Structural Determinants and Phosphatase Activity"

>O00192
VDSWV
>Q86Y97
GGEEL
>Q6P1M9
LILKL
>Q96L34
NDLEL
>Q6ZQN5
EGTEV
>P04921
KEYFI
>Q9BTT6
VTTSV
>Q9Y4F3
PITKL
>P35462
KILSC
>Q13588
QPVHL
>Q9Y4F5
ERFLI
>Q9ULJ7
KETPL
>Q9Y4F9
VATAF
>Q9NS75
KETRV
>A8MPS7
EPSLL
>Q9H251
EITEL
>Q9BYP9
QPSCC
>Q6PML9
DLEIL
>Q9P021
KQTSV
>Q5HYA8
QRFLI
>Q8TDG4
STDKA
>A0A1B0GUQ0
RALGI
>Q8NBX0
SSSEV
>P47211
NCTHV
>Q92633
DHSVV
>Q3KP66
IISQV
>Q5TGJ6
DRDSL

>Q4V321
KQSQC
>Q2M3T9
RSIQL
>Q9H5J0
PKTNI
>P13667
TKEEL
>Q96L42
KAINV
>Q8IYA2
NLDPI
>Q5T5C0
KWYQF
>Q9ULK0
HGTSI
>Q8IYA8
DDDGF
>Q9Y4G2
QNIFA
>Q9ULK5
SETSV
>Q00978
ILSLV
>Q9Y4G8
QVSAV
>Q8NGT1
THEHL
>Q8IWZ6
FFDAA
>Q04864
EFFQV
>Q8WWF3
KFSKF
>Q9UBT3
KIEKL
>P30260
ESDEF
>Q8N743
TDTSV
>Q9BYQ0
QPSCC
>O94979
NKLGV
>Q9BYQ2
QPFCC
>Q9H267
SEVKA
>Q9BYQ3
QPSCC
>Q9BYQ4
QPSCC
>Q9BYQ5
ASSCC

>P48546
LESYC
>Q9BYQ6
ASSCC
>Q9BYQ7
CGSSC
>Q9UI72
ACLLF
>Q9BYQ8
ASSCC
>Q9BYQ9
ASSCC
>Q9P031
IQEKC
>Q9C056
AGDAL
>P57735
CCISL
>A8MUP6
KVSIC
>Q9H720
PKYFL
>Q9NX47
EQEEA
>Q8TDH9
KFSTF
>P0DL12
MMFHI
>Q92643
MKFIF
>Q4VCS5
VEYLI
>P61266
GTLGL
>Q504Y0
QNIKI
>Q16613
RNSGC
>P0DJJ0
ECYGF
>Q9H5K3
AREML
>Q504Y3
LMSEF
>Q9UN36
MEVSC
>Q16617
GYETL
>Q9ULL1
SSSFA
>Q9ULL4
KVTDL
>Q9ULL8
GPSNF

>P49888
FRTEI
>Q30KQ4
SYSHI
>Q30KQ5
SELYI
>Q9HBW0
MDSTL
>Q9HBW1
QETQI
>Q30KQ6
EDDMF
>Q9NS93
TPLLL
>Q8NGU9
CESAF
>Q9UBU7
TFTGF
>P16233
TLTPC
>P16234
EDSFL
>Q9BYR0
ASSCC
>Q9BYR2
ASSCC
>Q9BYR3
QSLCC
>P30279
RDIDL
>Q6ZVL6
FEFQV
>Q9BYR4
GSSCC
>Q9BYR5
SGSCC
>Q6RFH8
LYELL
>Q9BYR9
RTSSC
>P11021
EKDEL
>Q9H0P7
STYYL
>P57740
YEIQL
>Q8TDI0
YVTDI
>O75747
GNSII
>A8K554
SVEVL
>P29317
VGIPI

>Q96ES6
PSTFL
>Q58FF3
EKDEL
>P61278
TFTSC
>Q16623
GGIFA
>Q9ULM0
GPTLL
>P17568
PKVAL
>Q8N0S2
IKELF
>Q9Y4I1
FISRV
>Q6AI39
SILEC
>P35499
KESLV
>A3KMH1
MLSSV
>Q8NGV6
FKSNV
>O94991
TFSQF
>Q15303
RNTVV
>P30281
TAIHL
>Q9HBX8
FASHV
>Q6ZVM7
ALFAL
>Q9BYS8
FSLQL
>O43424
RGTSI
>P21554
SAEAL
>Q8TF17
GGLAL
>Q86TA4
SSLVL
>P60409
QKSSC
>Q9C073
SSLMV
>P12814
GESDL
>O75751
SRSHL
>P43353
SCTLL

>P43354
DTLPF
>P43355
EEEGV
>Q8TDJ6
ILDIL
>Q9NX65
GRDAL
>Q5VY09
AIVAF
>P43358
EEEGV
>P29320
GPVPV
>Q9H741
ELFPA
>Q2M3W8
RRETV
>Q92667
YYTSL
>P13693
EMEKC
>Q16633
SVEGF
>Q9Y613
PGLEV
>Q9Y614
IYSKC
>Q8IYD8
LKSDI
>P22888
RYTEC
>Q6QHC5
AKDGL
>Q9HBY0
NKESF
>Q15311
KETSI
>Q8WWI5
GASSA
>Q5TD97
MDTDI
>Q9HBY8
DILDC
>Q9BYT1
THEDL
>Q8TF20
KTIQV
>P60410
SRLAC
>Q9BYT5
RTSSC
>P60411
QKSSC

>P60412
KKSSC
>Q8IV04
LDTRF
>P60413
SRLAC
>Q5PRF9
KTSTI
>Q9H0R1
GSLLL
>Q8TF27
PDECV
>Q9H0R5
HKLKI
>Q674X7
EVTNV
>P25445
IQSLV
>Q3LIE5
RAFHC
>P43362
EQEGV
>P43364
EGEGV
>Q8NDC4
RNLTA
>Q9NX76
EPLNA
>Q92673
PMVIA
>Q76G19
SVTTV
>Q9NVP2
SMDCI
>Q9NVP4
SLDDC
>Q12767
MNSPF
>A6QL63
KGSVV
>Q12768
FRTVL
>Q75V66
AKSTL
>Q02246
GSLEL
>Q5SVS4
KKLDL
>Q9Y624
SSFLV
>Q8IYE0
KPVEI
>Q6H8Q1
KALLF

>P52569
KTSEF
>Q96JQ0
TELHI
>Q96JQ2
DVSRL
>P22897
EHSVI
>A6NIX2
HVTEL
>Q9UBX2
LLEEL
>Q9UBX5
SQYPF
>Q9BYU5
RTSSC
>Q8TF32
MWESL
>Q7KZI7
NELKL
>Q9NQU5
QTSTC
>O15013
PLLNI
>Q86TC9
ESDEL
>O15015
LSFSL
>O15016
SISQV
>P57773
TDLQI
>Q5T0L3
AYLCL
>A0A1B0GUV1
NDSTI
>P0DTE7
AESKL
>Q7Z7B8
TIFTV
>P0DTE8
AESKL
>O43900
NCIVA
>Q9BVA1
GEDEA
>P20248
ETLNL
>Q6IB77
NCVPL
>Q3KNT9
AKSGL
>Q9BTZ2
TPSRL

>Q9ULP0
MEVSC
>Q8WY21
AQYAI
>Q9Y4L1
KNDEL
>Q8N0V4
VDLSL
>P0CI25
CCIHF
>P0CI26
CCIHF
>Q9HDB5
REYYV
>Q9HDB8
LEIKL
>P15407
TLLAL
>P15408
TLLAL
>Q8NGY5
IGILA
>A8MPY1
WGVYV
>Q9UBY9
TEIKI
>Q8TF40
AQILL
>P16278
WLDHV
>O43451
WISTL
>Q9BYV7
TFIPI
>Q8TF46
NNYGI
>P59025
FRSSV
>Q9NVR0
SQIEC
>Q7Z7C7
DDEDL
>A0A1B0GUW7
TNENL
>O43913
LYDFL
>Q16663
KPYSI
>Q12789
KWIHL
>Q16666
PDFFF
>Q2TBC4
HCTMC

>P63000
KCLLL
>Q6PHW0
IMVTV
>P25929
DNEKI
>Q8TA94
WRETI
>Q8N0W4
STTRV
>Q8NGZ0
CLFLC
>P01019
PLSTA
>P47736
PQLGC
>Q8WWL7
QGLVL
>O60513
FWFGA
>Q15349
TSTRL
>Q9BYW1
QSTEL
>Q9NQW1
HKLLV
>Q8IV32
PYSAV
>Q8N5S1
KTLGI
>P55145
ARTDL
>P07550
NDSLL
>O43463
RKYLF
>O15031
KVTDL
>O75792
SATSL
>O15037
LSLNF
>Q9UGV2
MEVSC
>A0A1B0GUX0
RDLAL
>Q96EX3
AEVAA
>Q12791
QEERL
>Q5VWM6
LHLCC
>Q16671
TLSPV

>O43924
RLFYV
>Q9Y653
SSSRI
>Q16676
RISNC
>O43929
SLSWL
>P56470
SYVQI
>Q3KNV8
KMDLL
>Q6ZQV5
SMEKF
>Q5T5J6
LNYRI
>Q96JT2
AKYSA
>Q8NA29
LASIL
>Q96BD0
LQSSV
>P02808
QQYTF
>Q7Z407
VCTMV
>Q7Z408
VCTAV
>P37235
SASQF
>Q495Z4
PFSLL
>Q96Q77
FHIRI
>Q8TF62
KTVKL
>Q6ZNB5
AMIIF
>P56937
SGSCL
>Q8IV45
LDEKL
>O15040
EWEVI
>P59044
LISTF
>P11086
QKVGL
>P59046
LDIGC
>Q14032
VTSQL
>O15049
ESSKI

>A0AVF1
NRVSI
>Q01415
VLLEA
>P29372
QDTQA
>P41212
QEDEC
>Q9H799
WALDL
>Q96EY8
ESEGL
>O43933
KVTLA
>P06241
PGENL
>O95415
GATFA
>P42081
SDTCF
>O15504
ELLNV
>Q49A92
VYSGL
>Q8IYI6
TTSVV
>P19320
QKSKV
>Q02747
ACTGC
>Q15369
NFLDC
>Q8TF71
SDSII
>Q8TF72
LTSPL
>Q0VFX4
SNSSC
>Q5HYJ1
IPFIL
>Q9H0W5
QEDQL
>P81408
RETGL
>Q9H0W7
KSTFI
>P59052
VPLAL
>Q86TG7
SYSTL
>Q9H0W8
SRLLA
>Q96GA3
EGLKL

>Q8NDH6
ELLNA
>Q5XPI4
TSSAA
>Q96EZ4
IILTC
>P41225
PLTHI
>Q16690
TATSC
>P41226
LHYEL
>Q16695
RGERA
>Q6P1W5
NGINF
>Q02297
DPIAV
>Q8IYJ0
PLVNL
>Q9Y676
PQSAL
>Q07960
DPSGL
>Q14500
RESEI
>Q9Y4P1
EILSL
>Q8WY64
NLTVI
>P01040
ELTGF
>Q5K4L6
GNLRI
>Q9NSA0
ESTSL
>Q9NSA2
KISSL
>Q15375
TGIQV
>Q7Z2K6
DLFVF
>Q9BYZ2
NKLKL
>Q6ZVT6
YYELV
>Q9NQZ5
RIEYA
>Q9H0X4
YQSEA
>Q5HYK3
SGFKL
>O15062
PSTLL

>P15907
RTIHC
>Q5HYK9
SEEKA
>Q7Z7G1
HLLPL
>P41231
KDIRL
>P41235
KQEVI
>Q5T742
PCSVI
>Q9Y680
QHDEL
>Q9NVV9
VEVPA
>Q86YD3
DEIWL
>Q86YD5
GTEEV
>Q658P3
KTSHV
>P25963
QRLTL
>O95436
ECTAL
>O15524
FPFQI
>Q6XR72
NRTHF
>Q8IYK4
SRDEL
>A6NC51
LPVQL
>Q4G0M1
VLLGV
>Q14512
QDTSC
>Q8IYK8
DLSVL
>Q2M218
TLYIC
>Q14515
ENLLF
>A6NC57
STVTL
>Q14517
QHTEV
>A0A075B734
ALEHF
>P47775
SPSDV
>P54315
TLTPC

>P54317
SLYPC
>Q02763
AEEAA
>Q9BS26
DRDEL
>P10243
RALIL
>Q15389
RPLDF
>Q9UQQ2
QYTPL
>Q5T292
QVIEL
>P42566
EISEA
>P60484
QITKV
>Q86TI2
LQEYL
>Q8N5W8
INEGL
>Q8N5W9
LSDEC
>A6NNZ2
EEEVA
>O15075
PNSPF
>Q14061
LGFKI
>O15079
GGSQL
>Q5VWQ0
STTSV
>Q32P28
PKDEL
>P41240
HELHL
>A2RUQ5
MELTF
>Q9NVW2
RESVV
>Q99645
VGSLV
>Q9BVG3
NTVRI
>Q5VWQ8
NSSNC
>Q6ZSA7
EEEEI
>Q5K651
DIEIV
>Q9BVG8
RPLPV

>P06276
SCVGL
>Q8N2C7
DESHV
>Q86YE8
RSIKV
>P59533
SRTLC
>Q9Y698
RTTPV
>O95447
RKIII
>A6NKF1
PGSPA
>O15535
VAELV
>Q9ULV1
SETVV
>Q14520
SESGF
>Q8NA61
KPSRV
>Q9ULV4
AKIAA
>Q14524
RESIV
>Q8NA66
DLFVA
>P22001
IFTDV
>P36402
SSSPA
>Q14527
TLIDL
>P49023
LKLFC
>Q7Z444
GCSVA
>P36406
VLDVA
>Q96BH3
TWVYC
>Q5PT55
RNFLI
>Q15391
STDTL
>P06730
NRFVV
>Q15393
TRYAF
>Q9H2A3
ARDCI
>P10253
LVSWC

>O95907
AAESV
>Q13200
PNYDL
>P0C5J1
LNLTL
>Q9UIB8
YEIVI
>O15083
EGIWA
>Q96GD4
LQSVA
>A2RUR9
NLDPI
>P60953
RCVLL
>P06280
LKDLL
>Q9NVX7
QFVLA
>O15540
HYEKA
>Q8IYM0
LTLDV
>O95455
EPFPV
>Q8IYM1
SDDEF
>P59544
CQIRC
>Q86YF9
DTSDV
>Q9ULW0
TRFHC
>Q9ULW2
SPTCV
>Q7L4I2
GMDAV
>P20774
IGSYF
>Q96JY6
LSSRA
>E7EW31
SLFPL
>Q9NSD5
LESHC
>Q86V24
EEDAL
>Q86V25
YQIRI
>P10265
IGYPF
>Q6ZNG0
TPVQA

>P10266
SAVIL
>Q4J6C6
KYLKF
>Q9C0A0
KEYFF
>Q9C0A1
TLLAL
>P59091
PALGA
>Q5T0T0
EIIHV
>Q32P44
SSLDV
>P10721
VHDDV
>P19827
VPDIF
>Q99665
DSLML
>Q2NL98
FGTTV
>Q86YG4
LSSAI
>O15551
RKDYV
>P46937
FLTWL
>Q6IQ20
DDENF
>P50454
MRDEL
>Q96D15
HHDEL
>Q6IQ23
LGSVC
>Q4G0P3
KGITL
>Q6IQ26
KGIDI
>Q9ULX7
ATTEA
>P18507
SYLYL
>Q9NSE2
YPFQL
>Q02790
VETEA
>P06753
DMTSI
>P31213
IPFIF
>O95926
RGTAV

>A5PLL1
QLVNF
>Q8N5Z0
IKESL
>Q86TL0
DFVFL
>Q8N5Z5
LGVPI
>Q13224
IESDV
>A1L4L8
KDTLV
>Q86TL2
FGLPV
>Q13228
SDIWI
>A8MWD9
ALERV
>Q96GF1
WLLIA
>Q9C0B5
YEISV
>A6PVI3
RQTQI
>Q7Z7K2
PPLFF
>P18054
NSVTI
>P19835
AVIRF
>Q15878
EDDKC
>Q9NVZ3
GWVQF
>Q99674
ENDEI
>A6NM03
AHSTL
>Q99677
LESTF
>Q70EL3
SVTQA
>O95471
SKEYV
>Q9UF02
SSSPC
>Q70EL4
PESSF
>P14625
EKDEL
>Q96LB1
RSSLV
>O95477
KESYV

>O95478
AVLLV
>Q8TAA1
CGSKL
>Q14554
KKEEL
>P04114
LTIIL
>Q8NA97
MDTAL
>Q8TAA9
SETSV
>P41732
QYEMV
>P15498
YSEYC
>Q8N7B6
CSICC
>Q86V42
EEFYI
>P27707
FLSTL
>Q5JZY3
QGVQV
>Q6IEU7
CKIAV
>P14174
NSTFA
>A8MWE9
TVVPL
>Q9C0C4
EESSV
>Q96GG9
KSTTV
>Q15884
RETVL
>O00300
KISCL
>O00303
KLVNL
>Q5TGY3
TVTSL
>P10745
LQDHL
>Q7Z7L8
SDSAL
>Q9UNA1
YVEFL
>O95484
KRDYV
>Q6PJG9
EESVV
>P63092
QYELL

>Q8TAB3
KDIVL
>Q8TAB5
PQDQA
>P63098
MVVDV
>P50479
KVELV
>Q9ULZ9
QALTL
>P41743
AEECV
>Q8WWU5
KVESV
>Q8N7C4
TIDVC
>Q71RG6
LVLQV
>P19397
QTIGL
>Q6PG37
FISLL
>Q7Z2Q7
EHSAL
>O95948
TCTKA
>Q9UIF3
QLELA
>O71037
VTVSV
>Q9C0D5
IESNV
>P0DV77
SLLDL
>Q6X4W1
FDDVL
>Q9NXC2
MSLYC
>Q6RUI8
ELLLV
>Q9C0D9
KNIGL
>Q8TDW7
HQTQV
>P40429
HGLLV
>P18077
YPSRI
>Q5JWF2
QYELL
>Q7Z7M8
PRLQC
>Q7Z7M9
KYYEA

>O95490
LVTSL
>Q13705
KESSI
>Q8N2H4
PKSNV
>Q86YJ5
RVTTV
>B2RTY4
NEFMV
>Q9H5Z6
EEFFI
>H0Y7S4
DQYCC
>O95498
NIVML
>P14649
HILSV
>Q01959
HWLKV
>P22059
CPDIF
>Q5JT25
NRSYC
>Q5VTB9
RRIYL
>Q9BS86
AKTCL
>P00709
LCEKL
>P31249
KLTHL
>Q9HA90
LLVSC
>Q13255
SSSTL
>Q9BQQ7
VKTAI
>Q9UIG5
LSSLI
>Q13258
MESSL
>P36915
GEDEC
>Q9UIG8
MESVL
>A6PVL3
GAEEC
>Q96GI7
ILESI
>Q8N402
EDFAA
>O00322
FYTML

>P10768
KYLNA
>P62244
LGFFF
>P14651
KLTHL
>A8MT33
APDAL
>Q9UF33
KGFHV
>Q6PJI9
LESTF
>Q1HG44
ITTNL
>A6NKL6
ESTGI
>Q14584
ERLSA
>Q14586
TREKL
>A0PJK1
YAYFA
>Q96BN2
GLLLC
>Q3Y452
SPYEL
>Q9BS91
DEDTF
>Q96BN8
EETSL
>Q9NSI5
NTTVV
>Q8WWW8
RQSKV
>O95967
GAYTF
>Q9H2G9
ELIAL
>Q96I15
LEDQA
>Q6ZNL6
DASVL
>Q8NF86
ARVSF
>Q13268
YSTRL
>P39023
KEEGA
>P54845
SHLFL
>Q8TDY8
VSSSA
>Q9NP55
FVIKV

>Q9NP59
NTSVV
>O14734
SESKL
>Q9H7C9
FHSTC
>Q9BVN2
TSLVL
>Q8IYS4
PLEQL
>P35609
GESDL
>Q01970
ENTQL
>Q14592
ETLPL
>Q01974
VQLEA
>P43005
QTSQF
>P43007
KESVL
>P08034
RCSAC
>P62714
PDYFL
>Q9UK00
NAIHA
>O95970
VDLSA
>Q8WWX8
WGYFA
>Q86V86
SSESL
>Q9UK08
FCVLL
>Q86V88
SPFEA
>Q96I25
LAEQV
>P52209
SSYNA
>Q9BQS8
GSDFL
>Q9NXF8
PEFSV
>Q9BX66
KPLYL
>Q9BX67
SSFVI
>O00341
LETNV
>Q08426
PSSKL

>O14744
YTIGL
>O14745
LFSNL
>C9JQI7
RKEVI
>P35610
CRYVF
>Q9Y6A5
KMEKI
>P14679
YQSHL
>Q96D71
PFSHL
>Q8IYT4
EFESV
>P84074
SASQF
>Q9Y4Z2
FSDFL
>P53539
SLLAL
>Q8TAF5
SCEEL
>P18564
LSTDC
>O75417
KDFDV
>Q8TAF8
GTEEV
>Q5GLZ8
GFSLI
>P30408
QQYDC
>Q8N7G0
GLLRF
>Q5VTE6
FRLEL
>Q8WWY8
PELQL
>O95983
EMEHV
>O95985
AAYFV
>Q9UK17
KVSAL
>P58294
KNINF
>P31275
ALSFF
>Q9H2I8
RDDQL
>Q6IEZ7
VRFVL

>P04626
LDVPV
>P04628
LHECL
>Q3KR16
SASEV
>Q9C0H9
SSISF
>Q08431
ELLGC
>Q9NP72
YCSVL
>Q96GL9
ISTDV
>Q9NXG6
ARVEL
>Q13740
HKTEA
>Q9BVP2
STDYV
>A5LHX3
GTETV
>P62277
SALVA
>Q6ZSJ9
TEVTV
>A8MT65
ERETL
>A8MT66
PDVTI
>P25100
RETDI
>Q8IYU2
GYTMA
>A8MT69
LLLDF
>P40925
FLSSA
>Q8WYA0
DRLIL
>Q8IYU4
DLLSA
>Q8WYA6
LLENF
>P18577
LAVGF
>A0A0J9YWL9
SVDPA
>B2RBV5
GCECC
>Q8WWZ7
DRVVF
>Q9H2J1
GNLGA

>P09848
PVSSF
>O95994
LKTEL
>P00746
DSVLA
>O95997
CDIDI
>Q9UK28
DLTKL
>Q9H2J7
PESDL
>P00749
NGLAL
>Q13296
LCDLF
>P22557
VTTYA
>P36957
LLLDL
>Q8NDT2
VRDTA
>Q9NP80
FFSKL
>Q6TDP4
SSTSL
>Q8N441
IHYQC
>Q68CJ6
PGTSL
>Q8N448
PGSLV
>Q68CJ9
AGDEL
>Q9Y6C2
ELEHA
>Q9BVQ7
NVEGI
>A6NKP2
AVERL
>Q5TF21
LFFVL
>P04181
TILSF
>Q96LI9
LHSIF
>P84095
SCILL
>O75437
EILQV
>Q6DHV7
RVLHI
>Q96BR1
EDLFL

>P08069
QSSTC
>Q9UK32
TSTGL
>P17252
LQSAV
>Q96QE4
SISGA
>Q96I51
AKSFI
>Q5T2D2
EVYLI
>Q9NZ01
IPFLL
>Q8IVB5
HSSNC
>Q5T2D3
AALNI
>Q5HYW3
GRIRV
>Q92800
ETDVL
>A0A0U1RQE8
NLVPL
>Q92806
SESKV
>Q9NP90
GSSCC
>Q9BX93
EKEEL
>Q9NXI6
ISSIA
>Q9NP94
IALWA
>Q5TA79
SGDCC
>Q9NP98
ETEEL
>O14775
LRVWA
>P07203
GPSCA
>Q8TC07
RLTPA
>Q8IYW2
SLFLI
>Q9Y6D9
RQTVA
>Q8IYW4
SSDQI
>Q96LJ7
YTSKF
>Q9NU53
DKTCI

>O75445
TDTHL
>Q8N912
GLLRL
>Q5TF39
KGTNV
>P29017
YQDIL
>Q14D04
VTTYL
>Q9UK41
RFLHA
>B7U540
RGSEI
>Q96QF0
FKEEL
>Q5SSQ6
GPTRV
>Q9P202
FNVML
>Q9P203
KKSAL
>Q8IVC4
KVVSC
>Q9P206
QKELA
>Q6IV72
KPSSL
>P49591
EVTDA
>A7MCY6
ENSKI
>Q9P0I2
QTSIF
>P0DTW1
GQSQC
>Q92819
MVLDV
>Q6T423
RSSVL
>Q8N465
LPSQA
>Q96N20
QKLHL
>Q15007
QGSVL
>Q96N21
AFLNA
>O14786
TYSEA
>O14787
AFYGV
>Q96LK0
SADEF

>Q86YQ8
LQTQI
>Q9Y6E7
LIDPC
>Q8IYX1
KDFFL
>Q49AH0
PKTEL
>Q8IYX3
DEDGV
>Q5SNV9
AQLQL
>Q8NAB2
SNLCA
>P05981
MVTQL
>Q8IYX7
LEVLA
>Q7Z2Y5
SNYDV
>Q9UK55
NPTLL
>P27797
AKDEL
>Q9UK59
DDDAA
>O75912
LETAV
>Q75QN2
AKLYF
>Q6JEL2
STLPV
>O75916
PWESL
>Q6EMB2
AHTKI
>Q8IVD9
GAVQF
>Q8WV07
SGLSF
>Q92823
MNSFV
>Q6ZU52
EALMV
>Q12913
NGYIA
>Q92828
GSEQL
>Q15012
PYLPA
>Q15018
QTSQI
>O14798
LIVFV

>Q9Y6F1
LEVHL
>Q86YR6
TKTNI
>Q9Y6F7
KIDEF
>P53582
FMSQF
>Q9Y6F8
KIDEF
>H3BS89
NGTVC
>Q96LL4
VNDNV
>Q9Y6F9
LSLCL
>O75460
TPDAL
>Q96LL9
IGFYI
>Q75LS8
KQDEL
>Q07157
LIDHF
>P16415
WKDTL
>Q5VTJ3
PQTRV
>Q9NSP4
SLEDL
>Q7Z2Z1
WLEDL
>Q16348
KKTKL
>Q8IVE3
GPTLL
>Q11130
GWFQA
>Q9P0K1
WETSI
>Q07617
RQYEL
>A6NFX1
SYSLA
>Q92834
SCTIL
>Q9NXL2
KMTYA
>Q8NDX6
GQFSL
>Q92839
YRVQV
>P13866
HAYFA

>Q8N488
NDESF
>Q13790
GSLEI
>Q9Y6G1
VLLLL
>Q6PJQ5
SLFDL
>Q8N2Q7
STTRV
>O43149
WNVEC
>P07237
VKDEL
>Q08AD1
LPTKA
>P61916
IVSHL
>P78411
GMSDI
>O75474
SGSLL
>P78413
KPFCA
>Q8TAL6
DYLKV
>P78417
CDYGL
>Q9H461
PLSQV
>P16422
RELNA
>Q9H469
VNLQV
>Q8N7M2
TDETL
>O43603
TVDVA
>Q9BSA4
NQFPA
>A0MZ66
DSSNC
>P11217
PDEAI
>Q8IVF6
EVLLC
>A8MWP4
PLSGV
>Q8IVF7
DESNC
>Q86TX2
IPSKV
>Q9P0L0
GKFIL

>Q9P0L2
NELKL
>H0YKK7
NITII
>A6NFY4
NPSTA
>Q12933
DLTGL
>Q02410
QPVYI
>Q08495
KASLF
>Q15034
GFSLA
>Q15036
GDEDL
>P52732
AQINL
>Q9UP38
GETTV
>P52736
SLYSL
>P52737
MWESL
>Q9BVV6
GADTF
>Q9Y6H3
YYSNI
>Q9Y6H6
RVSMI
>O43157
KVTDL
>Q9Y6H8
EDLAI
>Q1MX18
EESFV
>Q8TAM1
SEDEL
>Q52M93
QSSHA
>A0A0A6YYD4
CASSL
>P78423
VLVPV
>O75486
AFLAC
>Q08AE8
TISEI
>A6NCE7
MKLSV
>Q9H477
PLTLF
>Q8N7N1
LNLTL

>Q5TDP6
LEYFI
>A2RRL7
KDLQA
>Q5VTL8
KDETV
>O43612
GQSGI
>Q9BSB4
DTLAL
>Q6UX27
AALKV
>Q9Y345
LGTQC
>Q6UVK1
GQYWV
>Q9P244
LESTV
>Q6PEW0
FCSSI
>P11226
CEFPI
>Q6PEW1
FSEPL
>Q8IVG5
DIEVI
>Q9NZ53
EDTHL
>Q86TY3
SEDEF
>O75947
PIENL
>Q8WV35
LTLTL
>A6NFZ4
INEGL
>Q17R89
ESTAL
>Q8NDZ6
YLTVA
>Q15040
WRTDV
>Q92859
AITTA
>Q15042
DTSFF
>Q12948
DCSKF
>Q15046
VGTSV
>P52743
QKIHF
>Q15048
DYFSL

>Q9H7L9
RRSAA
>Q9Y6I3
NPFLL
>Q9Y6I4
GSDKL
>Q9Y6I7
LSYRI
>Q8TC59
NLFFL
>Q5VV41
VETDV
>Q8N961
ITSRV
>Q8N967
ISSVA
>P16444
CLSLL
>Q9H488
LRDEF
>P60602
MGIRC
>P60604
KSLGL
>Q96QK1
EGLIL
>Q6NVV0
YDLDL
>P52292
GTFNF
>Q9UK96
FCTIL
>P52294
EGFQL
>Q9P253
QLSWL
>O75954
KKYDA
>Q8WV41
MYDNL
>Q86TZ1
YASVI
>Q6PEX3
SCSGL
>B4DXR9
TGEKL
>Q6PEX7
FPSEL
>Q9HAD4
LYLAV
>P15121
FHEEF
>Q9P0N5
AFSRI

>P08588
SESKV
>Q12951
EGTEV
>Q9P0N8
EETPV
>Q96GT9
GKSQV
>Q16831
KLSKA
>Q15053
PSLPV
>Q12959
AKEKL
>Q6ZSR3
CADSF
>Q9H7M9
NFEVI
>Q9BVX2
NSTAI
>P52757
EDVLF
>Q96N76
QALQL
>Q96N77
GTSVF
>O43175
FQFHF
>P47900
GDTSL
>P47901
ETIIF
>Q9Y6J9
LYLPL
>Q5XUX1
WRLQA
>P16452
PELSA
>Q9H492
ETFGF
>Q01118
IQSQI
>Q8IX03
SADDV
>Q96S37
KSTQF
>Q8IX04
QQLKL
>Q8N7P7
ADTAL
>A6NHC0
CCVLV
>P11245
GSLTI

>Q6PEY0
QVLSV
>Q6UVM3
EETQL
>Q9P266
RVERV
>Q6PEY1
GKVWV
>O75964
IGYDV
>Q8IVI9
TATKA
>O75969
LMVNL
>P30953
TFFSL
>Q12965
YVTKI
>Q12968
RSDGL
>Q02446
NMEEF
>O60235
QQTGI
>Q15067
LQSKL
>Q8N2U0
LALAL
>P56645
VEDSC
>O43182
PETLV
>Q8TC76
KVSHV
>Q86YW7
ECETI
>Q86YW9
HPSHF
>Q9Y6K8
IDSIF
>A6NKX1
DRVSF
>Q5VV63
QGTCV
>Q8WYJ6
QSDAL
>Q8TAP6
YRSVL
>Q9UDX4
ELTPV
>Q8TAP9
GRYFC
>Q15528
PTEHA

>Q8N7Q2
IIFLL
>P55327
TQESL
>Q9BSE5
KVTTV
>Q16394
DIERL
>Q9P278
AQILL
>P68402
QTTIA
>P30968
GYFSL
>Q12972
PSLLI
>Q92887
NSTKF
>O60241
FQTEV
>Q12979
FSTDV
>O60242
FQTEV
>P11712
CFIPV
>O60248
PLTHL
>P11717
DLLHI
>O43196
ATSIL
>Q9Y6L6
SETHC
>P28223
KVSCV
>Q96LR2
DVTFL
>Q9UNP9
CGEYV
>P61962
EILRV
>P61964
WKSDC
>Q8WYK0
FVSTF
>Q8WYK1
REYFI
>Q2M2E3
KDTHV
>A0A1B0GTB2
IYTTL
>Q96LR9
AGLFF

>Q9UDY2
RDTEL
>Q7Z4G1
VIETV
>A6NCI8
YGYTI
>P16473
MQTVL
>Q8N999
IIFDV
>Q5NDL2
KHDEL
>Q8N7R0
QPEDV
>Q5T442
TTVWI
>Q6UX68
PSFFI
>O43657
QYEIV
>Q9P281
VPILC
>Q9Y385
FDFEL
>Q6NVY1
SDLKF
>O95136
GNTVV
>Q9P287
EYLSV
>Q9NZ94
STTRV
>P15153
ACSLL
>A0A1W2PP81
KRIMF
>Q9H977
RFSSV
>Q9NXR1
SSSSC
>Q12983
STSTF
>Q15084
GKDEL
>Q9NPB6
SGFSL
>Q9NPB8
NVENA
>Q9NPB9
STFSI
>P42261
GATGL
>P42262
ESVKI

>Q8TC90
QDFNC
>P42263
ESVKI
>P24347
ANTFL
>Q9Y6M5
PESSL
>P01213
ELFDA
>Q8TC99
YLFPC
>Q9Y6M7
AETSL
>A8MTA8
KQLLA
>P26992
SSLLI
>Q96DC8
SHEPV
>O60711
KLFPL
>Q15544
KIIFF
>A2A2Z9
IYFLL
>Q02928
DKDQL
>Q7Z4H8
SREEL
>Q96S65
EPVPV
>Q6UX71
VSEQC
>Q8N7S2
SEDDF
>Q6UX73
QSVGL
>P21796
LEFQA
>Q9P291
VLTKL
>Q8IVL0
LESTL
>Q8IVL1
LESTL
>P11274
FSTEV
>Q8IVL5
PKDEL
>Q8IVL6
VREEL
>P50135
IVIEA

>Q8IVL8
CMSLL
>Q07687
AGTIF
>Q5TAA0
RAVSF
>P30988
QESSA
>Q96GX5
SGFSL
>Q9NXS2
EYLGL
>Q9NXS3
GLTAL
>Q9BXC1
TPELC
>P0DN77
SVSPA
>P0DN78
SVSPA
>Q99807
LSERL
>P55808
EPENV
>O60260
HWFDV
>Q8WTQ7
VCLLL
>P52790
QLTRV
>Q9NPC6
ESEDL
>O60262
PCIIL
>O60264
KKLKL
>P52799
IYYKV
>P63218
VCSFL
>P15621
WKDIL
>P01222
VGFSV
>Q9Y6N9
ELTFF
>Q96LT6
PAVAL
>Q8TAS1
YQTLL
>P01229
GLLFL
>Q01151
KTELV

>P20929
YVEAI
>O60721
CPVSV
>A0PJZ0
ALDKV
>A0A3B3IT33
CCSHF
>O60725
VKVDL
>Q5JTB6
LGDGF
>Q6ZPB5
HRTDI
>O95150
GAFLL
>Q06828
SLIEI
>Q86VF5
CLTFI
>O95153
RRVQC
>Q86VF7
KALLL
>Q9H2V7
ASVLI
>Q9HC07
PDSGF
>P25685
QVLPI
>P25686
RCLIL
>A6NHG4
FIYFI
>Q96A05
RKFFI
>Q14232
IKLYL
>Q8IVM8
EVTQF
>Q9H992
TFDIA
>P0DN82
NSSEA
>Q8WV99
NCSLC
>P68431
RGERA
>Q9H7R0
WRDTL
>A3QJZ7
DQYCC
>Q6P3V2
QSSHA

>Q8WTR7
QQYCL
>Q9NPD7
TWLSF
>O60279
EMEKV
>P42285
ASLYL
>Q8WYN0
EILSV
>P20933
KVDCI
>A6NCL2
QCLPL
>Q8WYN3
ETVPV
>P09001
SITFA
>Q9NSY0
RGTQA
>Q9HC10
KILGA
>P81605
LDSVL
>Q8N7U7
LLLDL
>P14314
DHDEL
>Q9H2W6
FVSDL
>O95164
CCVIL
>O95166
SVYGL
>O15254
LKSKL
>P50150
FCTIL
>O15255
SCVLA
>O95169
VHYEI
>Q14240
VADLI
>Q8NFH4
WVTEV
>Q8IVN8
SFIFI
>Q14244
TAEVI
>Q8NFH8
PVTVL
>Q96GZ6
AKTSI

>P23508
NETSL
>Q9NPE2
FLYRI
>Q16890
EELQC
>Q05513
TEESV
>Q99828
FKIVL
>O95620
DHYGI
>O60284
KGIHV
>O60287
AASDA
>O95628
HTTVA
>P50613
KKLIF
>P82930
KGTPV
>P0C2L3
ISTDV
>A0A0A6YYL3
TKTSI
>Q7Z628
KETLV
>Q08AM6
RRVVL
>Q2M2I8
QLIDL
>Q96DF8
ASDFF
>O60741
FASNL
>Q96S90
EIFKL
>Q5VTT2
PIVPI
>Q96S95
PPSGV
>P09016
DLTTL
>Q9BSJ1
CCSHF
>P09017
DITRL
>Q8NH01
ATSDA
>Q96S97
VFVKV
>Q8NH02
VRFVL

>Q8NH04
KVTTF
>Q8NH05
IPLPC
>O95170
LFSGV
>Q05066
HWTKL
>P01701
SSLSA
>P0C7H8
RTSSC
>P01705
CCSYA
>P32241
EVSLV
>Q9UC06
SGEKL
>Q9HC29
TRLLL
>Q96A22
NGTLV
>O15266
EALGL
>P32247
AEDRF
>Q5T2Q4
KAILF
>O15269
QAVLL
>P51946
LVESL
>Q14254
TGVQV
>Q96A28
GSSPA
>Q5VYP0
LWEGI
>Q14257
YHDEL
>P23515
VMLAV
>P29597
VFSVC
>Q8WTT0
KKIYI
>Q99835
ADSDF
>Q9NXV6
PQELL
>Q8N4B5
HLSTL
>Q6NSI1
NNENI

>P24385
RDVDI
>O60299
ESTEI
>P11766
TVVKI
>P24387
CLSGL
>Q9Y6Q1
DLTEL
>Q8NC54
NDYIF
>Q765I0
WKYCV
>Q76N32
PCEGV
>Q96LW1
ADLHV
>Q96LW2
YPIPA
>P47972
RLLDL
>Q5MNZ6
TDDKL
>A6NCN2
RALVL
>Q15583
AKLTA
>Q29983
STEGA
>Q6V0L0
NGLCL
>P19544
LQLAL
>P58400
KEYYV
>P58401
KEYYV
>O60759
EESRF
>Q9BSK0
KQEVA
>P42765
IQSTA
>Q8N7W2
SCLSL
>P42766
YAVKA
>O95180
ADDPV
>Q9UKA4
LLENA
>Q86VI4
PYVSA

>Q8NH18
KHFLC
>Q96QS3
GKEVC
>Q9UKA9
SKSTI
>Q9P0V3
DDFVI
>Q9NPG1
DGTSA
>Q5T953
AIVAF
>P13010
LLDMI
>P24390
LSLPA
>Q8N4C9
HQTYF
>P63252
RESEI
>Q6ZSZ5
DVIFF
>Q8NC60
GKINV
>Q5T7N3
RSLGL
>Q9Y6R0
FEIEL
>Q9Y6R1
RHTSC
>Q8NC67
ISIDF
>P40123
AEIMA
>Q9NUC0
KGSKI
>P28289
CRSGV
>Q96LX7
PPVSF
>Q8TAW3
AGEKL
>P40126
YTEEA
>Q8WYQ5
CTVDV
>P47989
WSVRV
>O60760
PQTKL
>O60763
DLDHI
>Q32M78
TRENI

>P05156
SQYNV
>Q15599
IFSNF
>A4D256
PILLF
>Q86X19
CIEEI
>Q9H4A9
LILWL
>Q8NH21
SSVKF
>Q53SZ7
QKSSV
>Q8N7X4
LRLRA
>Q05084
ELLNA
>Q9UKB3
RNYEI
>Q9UKB5
FEISC
>Q13409
TRIPA
>Q96QT4
VRLML
>Q17RB0
EDEDF
>Q9P0W2
ASEHL
>Q01658
DDDDI
>Q6ZUB0
LNLSI
>Q9P0W8
CPLDV
>Q3KRB8
REDNV
>Q68CZ1
DDLEA
>P00403
PVFTL
>Q5W064
IQDNL
>Q9UNW1
TSDEL
>A0A1B0GV03
KIISI
>Q96F07
LATTC
>Q9UNW8
IEESC
>Q96LY2
HRSVL

>Q14738
SQEAL
>Q8TAX0
SKIKC
>P49238
ALLLL
>Q8NAP3
ENVVL
>Q9NUD7
EPLPF
>Q7Z4N2
TETEC
>Q8N103
KESYI
>P05166
ANIPL
>Q9H4B4
DRSPA
>Q86X24
PKEHI
>P24864
GPEMA
>Q6NXG1
EWVCI
>Q86X29
ESLVV
>P01730
TCSPI
>Q13416
EEEEA
>Q96QU1
QSTSL
>P01733
CASSL
>Q9UKC9
CCVIL
>O15294
VTESA
>O15296
NSVSI
>Q96QU8
GTVKL
>P98153
LNTVV
>Q96A54
DDTLL
>C9JDP6
RNLVI
>P98155
DDDLA
>Q96A59
EMFEF
>Q01668
CITTL

>Q9NR09
SDFQL
>Q17RC7
LITCV
>Q9BXI6
EDTYL
>Q9NPI7
SILGF
>Q9NPI8
GLSSV
>Q6BDI9
YISIL
>Q3SXP7
QSSSA
>P82970
PQSIV
>Q9UFH2
LLLQV
>Q96LZ3
LVLIV
>Q8TAY7
RESEV
>O60784
MLFAL
>Q8N118
KLSEC
>Q13421
ASTLA
>Q8NH42
KRLCI
>Q13424
LGLLA
>Q13425
MGLLV
>Q9BSN7
VESPC
>Q13426
LFDEI
>Q96QV1
TSYIV
>Q06889
VTTCA
>Q96A65
KITTV
>P98164
EDSEV
>Q9NR11
KPFEV
>Q9NR12
AFSHV
>Q6UVW9
PKYFL
>Q5SZD1
EEIKL

>P98168
SSFLV
>P98169
SSFLV
>Q0VG06
SLILL
>Q9BXJ0
SPVFA
>Q9BXJ2
EDDEL
>Q9BXJ3
ASELL
>Q9NPJ4
LKVQV
>O95670
YRISA
>Q8N4F4
KVTQF
>O95677
ELEYL
>P27448
NELKL
>P32745
RISYL
>A0A1B0GV22
RALGI
>Q9UNY4
VLFGI
>Q63ZE4
LKEKA
>Q8NC96
NWVQF
>P01298
SPLDL
>P36639
EVDTV
>A0A1B0GTK5
RALGI
>Q6PI25
TLVSF
>P05186
LSVLF
>Q8N126
KEYFI
>Q8N127
RIVNI
>Q8N129
DREDL
>Q6YHK3
MELWL
>Q8NH51
CNIFV
>Q13433
FRINF

>Q13434
MLLML
>Q13435
KEFKF
>P0C7M6
EILSI
>Q9HC73
SYVAL
>Q9Y3A3
GESEA
>Q13438
DEFDF
>Q96IG2
CCIIL
>Q9HC77
MDTEL
>Q96A73
TPTGF
>P32297
AREDA
>P98172
IYYKV
>O75112
HTINL
>Q9GZL7
SHVGA
>Q8NFN8
KPTLV
>P30101
AQEDL
>O00505
KEFNF
>Q5SR56
QCTEL
>Q9H7Y0
YNDKF
>O95684
LEDVA
>Q14761
HVTAL
>Q5T7R7
LNDTL
>A8MTJ6
EGSEV
>Q96DL1
LNYIC
>Q9BU76
RGSEA
>P31431
NEFYA
>Q8N135
IDLSA
>Q9H4E5
CCSII

>P27918
EEEEL
>A6NJ64
HCTQA
>Q13444
SSLYL
>Q9HC84
EATAV
>Q8NH69
NKILF
>Q13449
LLSKC
>O75121
YESHV
>Q9GZM8
LPLSV
>O94822
RETFF
>O94823
SSLTI
>Q6IN97
STDIF
>Q9NZC9
FTSPL
>O75128
VPLLV
>O94827
TASEV
>Q58HT5
KLFFL
>Q9BXL5
SYVLF
>Q9BXL7
DEDQL
>Q8N4H5
KLDSI
>Q5RHP9
NNVQV
>Q09470
LLTDV
>P84243
RGERA
>Q14773
MKSQA
>A6NMK7
PGSKI
>Q9Y6W6
VETVV
>Q9Y6W8
TDVTL
>Q96DM3
RPTTF
>Q5SW24
VMTMV

>Q4FZB7
LRLNA
>Q5VVB8
SKLLV
>Q8N141
HNVKI
>Q8N144
RDLAI
>Q8N145
VDLSA
>Q7Z4R8
LEILF
>Q86X67
SSLPA
>P17405
RPLFC
>Q9UKG1
RESEA
>Q9BSQ5
DQDSA
>Q9UKG4
ITDQA
>A6NJ78
AAIKL
>Q9Y3C4
TKDVL
>Q9UKG9
NSTHL
>Q9HC97
CVTLA
>Q9GZN1
EKFDI
>Q04721
MQVYA
>Q9NZD2
LNYKV
>Q6TFL3
TQIGL
>Q4KWH8
FLLRL
>P98194
SFLEV
>Q96II8
QLSAV
>Q6TFL4
KCFKL
>Q8IVV8
VYLPA
>Q9HAR2
LVTSL
>P98196
SSLSF
>P18283
LKVAI

>Q9H9B1
AADPL
>Q8WVB6
IRDLL
>P23582
SGLGC
>Q4KMQ1
PALYF
>P09543
SCTII
>O14924
HATFV
>Q5J8M3
GGLLL
>Q96NE9
PEFVV
>Q9Y6X3
LASLL
>Q9Y6X6
WDTTI
>Q96DN0
PKVEL
>P49286
QADAL
>Q92502
PETKL
>Q8NAU1
LRSKI
>Q96DN5
HLLAA
>Q6ZR37
ESSEI
>Q96K12
STLKV
>Q6UXB2
FALPL
>Q6UXB3
LSLRL
>P0C7P0
VGSPL
>Q9UKH3
VTVSV
>Q9BSR8
LYTGV
>Q13467
SLSHV
>A6NJ88
KIIVI
>Q9P2D3
KTSFF
>P35346
QTSKL
>Q96QZ7
TDLSI

>P35348
NGEEV
>P49748
NPLGF
>Q96IJ6
NQIIL
>Q9P2D8
QFYPL
>Q8IVW8
ASVKV
>O75145
RTYSC
>Q9NZE8
TNLKV
>O00533
FPLRA
>O00534
AIFAF
>P80370
GDEEI
>Q3T906
NRIRV
>Q9Y6Y0
KIFQF
>P04350
EEEVA
>O75603
NNEDF
>Q8TCE6
QMLKI
>Q9NUJ7
KLLWC
>Q8N9F7
HNFSA
>Q8NH92
KISSL
>Q86VQ1
NYVII
>P49753
IPSKV
>Q5R372
KFVYL
>P35354
RSTEL
>Q9P2E2
GSEPL
>P22736
DTLPF
>P49757
FEIEL
>Q3BBV2
YRSVF
>O75151
GKLLL

>Q9P2E8
RVTTV
>Q53R41
VKSCL
>Q8WVD3
FQVNI
>Q9P2E9
EGTSV
>Q9NR61
IATEV
>Q6ZW05
HVTTV
>Q8NFR7
TVFKI
>Q5GAN3
LYSGI
>O75157
NVSSA
>A6NHR9
TKTDV
>Q9BZ68
IEYCI
>Q7L1T6
HSFTA
>O94856
IYSLA
>A0A1W2PPE2
KRIMF
>Q6ZUI0
GSIGF
>P09565
SDLRA
>O00548
IATEV
>O14948
DGDEL
>Q13936
YVSSL
>A8MV57
PKVWA
>Q9Y6Z5
RRTRC
>P08243
SAVKA
>Q9NUK0
NQLKF
>Q9H607
GSVYF
>Q9H4I0
MFYNI
>Q8N9G6
GGSCL
>O14490
AQTRL

>Q96K31
IQILA
>O14498
FLTSF
>Q9UKJ0
PSSDF
>Q5T4B2
PRDEL
>Q6UXD5
YEVSI
>Q9UKJ1
SVLKA
>Q96K37
NRYDV
>Q9Y3F1
AKFLA
>A0A087WWA1
IFLIL
>Q9Y3F4
PDVKA
>Q9BST9
LQSPV
>Q86VR7
VQLSL
>Q8IVY1
DHFSL
>Q9P2F5
SVTSV
>P49768
HQFYI
>Q9NR71
EVVTI
>Q9GZQ8
MKLSV
>Q9BZ76
KKEEC
>P31930
FWLRF
>B3GLJ2
CNFKL
>Q9H9E3
KRLRL
>P30153
VLSLA
>Q9HAU8
VNVSA
>O00555
DDDWC
>O94868
EITLV
>P10997
NYLPL
>Q9H156
AISQL

>P31937
EEETF
>Q9BXP2
TCTDL
>Q8N4L2
EHSFA
>Q9NPP4
KLVTA
>Q9BXP5
DVDFF
>Q03426
ALDGL
>Q96F83
NSDVC
>Q8TCG1
VNLSI
>Q96F85
KESFL
>Q8TCG5
TSTDF
>A0A1B0GV85
KKTVL
>P08253
DWLGC
>P08254
SWLNC
>Q8NAX2
LQVYC
>Q6NZ63
NFLTL
>Q9NUL3
SNSAV
>Q6P0N0
NSDSA
>P45880
LELEA
>Q8N9H8
ASSPF
>P13569
QDTRL
>Q13491
CCTKF
>B8ZZ34
TEVTV
>Q13496
VQTHF
>P04839
NKENF
>Q9UKK9
PFLKF
>Q9GZR2
CSDDA
>Q8WVF1
MMDEL

>Q9NZH4
SDIDI
>Q9NR81
GESNV
>Q96IM9
SKSPF
>Q9NZH5
YDIDI
>P61619
GALLF
>Q7L1V2
LFTGL
>P26639
AEEEF
>O00566
HKLKL
>O14966
SWSCC
>Q5SZK8
DSSEV
>Q9H9F9
AGEQA
>P48444
KYEIL
>A8MV72
GGSCL
>P48448
SCTLL
>P00492
AKYKA
>Q8N4M7
YPELL
>O43309
KSVSV
>Q96NI6
RLELI
>P43234
SSIFV
>P30622
DDETF
>P08263
KIFRF
>Q96DR5
LQTLI
>A0A1B0GTR3
ETSVL
>P61165
VGIYV
>Q92546
STITI
>Q96DR7
LETNV
>Q8N9I5
ELVGL

>Q16513
IADWC
>Q16515
EEIAC
>Q8N196
EPLEL
>Q6NXP2
HFLGA
>P52434
KKLAF
>Q8IXB1
NKDEL
>Q6UXF1
QVSEI
>P0C7T2
KVTTF
>P04843
ILDAL
>P22760
LKENL
>P0C7T7
PSLPL
>Q00889
EILLL
>Q9GZS0
EEDLA
>Q9P2H5
HRLVF
>P49788
ELSNF
>Q04771
LKTDC
>Q6ZW31
INVCL
>Q9NZI5
TLTEI
>O75185
HPEDV
>O00570
PLTHI
>A8K7I4
QLSIA
>O00574
SMFQL
>Q9H9G7
TMYFA
>O14975
KTLKL
>B2RXF5
KSLLV
>Q9H175
LPLAV
>O94889
PLLTI

>A0A1W2PPH5
KRVMF
>Q9NPR2
RDSVV
>Q5TAP6
KEEKL
>O43310
QKLTA
>Q9BXR5
RTDCL
>Q8TE02
DDLDI
>O43313
TVSNF
>Q9NPR9
GRELL
>Q96NJ3
QRETL
>Q3SXY7
PEYYC
>P62955
STSPC
>Q9NUN5
SVYSA
>Q9H4L4
CKLTV
>Q9UM47
RQVLA
>P38405
QYELL
>Q6UXG3
KFVSA
>P0C7U1
EVVTI
>Q86VU5
LAFKI
>Q9UKM7
IWTPA
>A4D0Y5
AGLLL
>P49795
SSSEA
>P40692
VFERC
>Q9NZJ0
HSTEL
>Q9GZT4
QSVSV
>Q8NFV4
RGFLV
>Q9GZT9
GKDVF
>Q9H9H5
RILNV

>P55000
CNSEL
>O14986
LDVYL
>Q6BDS2
PFFEL
>Q9BXS4
AHSEI
>Q5W0A0
SKLWF
>Q9UPI3
SEDHL
>Q13976
WDIDF
>P21452
THVEI
>Q9BXS9
SVTRL
>A0A087WT03
CIVRV
>Q8TCJ0
DLFKF
>Q1EHB4
ETTHF
>P43250
LPTRL
>Q9NW61
SGLQA
>P53778
KETPL
>Q96DT5
LLLEA
>Q5SW96
DLFSF
>Q96DT6
EFVLL
>P13591
NESKA
>P00973
TCTIL
>Q16538
PQLTL
>Q9H4M7
LQSSF
>Q6UXH0
AALPA
>Q6UXH1
SREDL
>Q9Y519
SDDEF
>Q9UKN1
VASTV
>Q96K78
AKESI

```
>Q9UKN7
EITLL
>Q9P2J2
QATLL
>P61647
KCEVA
>O00592
EDTHL
>B2RXH2
TNLSL
>P17931
SYTMI
>Q68EM7
ESTAL
>Q9H195
TSSSV
>P55010
DIDAI
>Q8N4P3
RGLTI
>Q5W0B7
EMSPF
>Q96NL3
IHTRV
>Q03468
KPEYC
>P12724
LDTTI
>Q96NL6
NLENI
>Q6NSW5
QMLKI
>Q6NSW7
QPEDV
>Q2L4Q9
QPTSC
>Q92570
DTLPF
>A0A1B0GTU1
LPLEL
>P16619
LELSA
>Q6S9Z5
GALCL
>Q16540
SWFGL
>A0A0B4J1U6
CASSV
>Q9NUP7
PETTA
>Q16543
KDVSV
>Q03923
EKLQI
```

>Q3I5F7
KHSKI
>P21926
NREMV
>P0C7W0
AESLL
>Q86VW2
DEDEL
>Q902F8
VTVSV
>Q9BSY9
RHTKL
>Q902F9
VTVSV
>P12271
ENTAF
>Q6ZW61
GFLFL
>Q9P2K8
YRILF
>Q9GZV5
FLTWL
>Q9NZL6
SKITL
>Q9BXU1
ANFDC
>Q8N4Q0
VNSKL
>Q9NPU4
YAFTF
>Q96P47
SPSLL
>Q5GH72
YETTV
>Q5GH73
YESSL
>Q494W8
SKDFA
>O43345
TGEKL
>P60328
TPSCC
>Q5GH76
YETTL
>Q53HC9
YHILL
>P60329
TPTGC
>P53794
VYFSL
>P07437
AEEEA
>Q9NW81
SPVPA

>P57678
KMSSF
>Q5TH69
YDIIV
>P57679
RRSNL
>Q6MZM0
PTDAL
>P20142
FATAA
>Q9H668
YYTAF
>Q9NUQ2
VTIKA
>Q2Q1W2
RILVF
>Q9NUQ6
VTLVA
>Q9Y530
TVYTL
>P38432
STEPA
>Q6NXT2
RGERA
>Q50LG9
YEIHC
>P38435
VHSEF
>Q96SI9
HYSFF
>Q7L3B6
MMDTV
>G2XKQ0
GHSTV
>Q9GZW5
ETEDV
>Q9NZM3
QKTLL
>Q8WVK2
LDFIA
>Q9NZM5
REIQL
>Q9NZM6
KVSAL
>Q9GZW8
SRSWI
>Q8N695
NGTRL
>Q6IF36
PQSIA
>Q9H9K5
DTSLL
>A0A1W2PPL8
KKVMF

>Q8NCE0
DQDDL
>O75688
SGEKI
>Q9H672
KFDDI
>Q6MZN7
KGTLL
>P0CAP2
SSDEF
>Q92597
MEVSC
>P21941
ENTVV
>Q02161
LAVGF
>O43813
PAFEL
>Q1A5X7
IKDEI
>Q9BUB5
PPTAL
>P07902
TATIA
>Q16563
PPTGI
>O43815
AKVFV
>Q9H4P4
GVEEI
>Q8N9N8
EEEAA
>P15311
EFEAL
>Q07837
LYTSC
>P15313
PDTAL
>Q86VY9
SETRF
>Q8NFZ0
LFLVF
>Q9Y3M8
PETKI
>Q8NFZ3
STTRV
>Q9P2M7
QTSSC
>Q8NFZ4
STTRV
>P51151
SSSCC
>Q9GZX7
RTLGL

>Q6IF42
YGLCL
>Q8NFZ8
EEFFI
>P55040
DLSVL
>P55042
DLSVL
>Q8N4S0
EKFEL
>Q8TE54
DHSEV
>O43365
KLTHL
>Q8TE57
SKSNL
>A6NEF3
NITII
>Q15700
SKEKL
>P48960
SESGI
>Q99502
ELEYL
>Q96DX5
FLLHL
>Q5VVM6
QKSEL
>Q99504
ELDFL
>O43820
PKEAV
>Q86XA0
AKDSL
>O95302
KHDEL
>Q96SK3
QRIHI
>Q5J5C9
STSAV
>Q86XA9
KTSFL
>Q2KHT4
EEEQC
>Q5JTV8
RGICL
>Q86VZ1
QESVF
>Q9UKR8
LELLA
>Q9HCD6
VESNV
>Q8WVM0
ENYRL

>P51160
TCLML
>Q96AD5
GALGL
>Q9GZY6
AATEA
>O60422
TFSKA
>P51168
EGDAI
>Q96P70
QTIGI
>Q9BXX0
FLSHL
>P38919
VADLI
>Q6ZMB5
PSEDL
>Q8N4T8
LQLIL
>Q8TE68
EMEVI
>Q69384
VTVSV
>Q8NCG7
SVDVA
>A8MTW9
PGSTL
>P29275
LGVGL
>Q7Z6E9
KSVTV
>P29279
YGDMA
>Q16581
NSTTV
>Q9Y561
ALLLC
>Q8IZ08
GDTSL
>Q460N3
ITFTA
>Q9H4R4
VKVFV
>Q96SL1
VVVSV
>Q96SL4
KREDL
>O15403
QMEPV
>O15405
QVSIF
>Q8NHC7
YSENV

>Q9UKS6
ECVGA
>Q9HCE6
VPLML
>P0DP42
AETVI
>Q96IU2
TKVLL
>A0A1W2PR64
RRIMF
>P51170
MLDEL
>Q9BR01
FDFYL
>Q00G26
PELDF
>Q6IF63
RKSPL
>Q02643
LTSMC
>P17980
LQYYA
>P10124
EDFML
>P55060
SVTLL
>Q9NPY3
PGTDC
>P11908
SHVPL
>B7ZC32
MTLLL
>Q96P88
SITSI
>Q86SG2
PRTRC
>P60368
QKSSC
>Q5ZPR3
GQEIA
>P60369
QKSSC
>A6NMX2
NKFVV
>Q7L8A9
YQIRV
>Q96FA7
HRSLL
>Q99527
FSSAV
>Q8IZ16
AQSPL
>Q9BUE6
ESFNI

>Q16594
DYDNL
>Q6UXN2
KGLML
>O15417
VPVLC
>Q6UXN8
LRSSV
>Q96IV0
KFSDL
>P33260
CFIPV
>P33261
CFIPV
>Q9Y3P9
GKETC
>Q9BR10
PGSCL
>Q9BR11
EQVQL
>Q6JVE5
QASVC
>Q6JVE9
KEELI
>O60447
YSTTV
>Q6ZUT3
NYFLA
>Q6ZUT4
CSDNV
>Q8IU60
KILDL
>Q8NE00
SETKL
>P60370
SPLAC
>Q9UPP1
GKLLL
>P60371
QKSSC
>Q9UPP2
SRSLV
>P60372
CVSLL
>P56856
KHDYV
>Q9NPZ5
VKIEV
>Q6ZUT9
KGVDV
>Q86SH2
FKYII
>Q06547
NKEAV

>Q86SH4
KKIYC
>Q96NR3
QITTV
>Q8TCQ1
EVVSV
>Q96NR7
LSSFF
>O60902
AALGL
>Q5VVP1
LWEGI
>P23219
PSTEL
>O43852
RHDEF
>P07942
YSTCL
>Q9Y586
SLDKL
>P07947
PGENL
>O95336
KHSTL
>Q03989
LNTKL
>A8MYU2
YSEPL
>O15427
PETSV
>O15428
RREGL
>Q9Y3Q0
LKEVL
>O97980
LTYRL
>Q9Y3Q4
LPSNL
>Q14416
TTSSL
>Q9NZR2
RETVA
>Q96IW7
PDYDV
>Q9H9P8
QRFEL
>P11926
ASINV
>Q9UPQ0
QPTTL
>P10147
LELSA
>Q9UPQ3
VPTII

>A8MVA2
QHSCC
>P55087
VLSSV
>Q9UPQ7
SVTTV
>Q9UPQ8
ILLMA
>Q7L8C5
HQLHL
>Q9UPQ9
GSDSI
>Q8TE99
HREGF
>Q01344
EDSVF
>P60842
VADLI
>P07951
DITSL
>Q8N1C3
GYLYL
>Q8N9S9
KEEDL
>O43868
NTVCA
>O15432
LLSTA
>Q9Y597
QEYSL
>O95347
AHVEV
>Q6UXP7
IKVNL
>Q5JTZ9
ALSQL
>A3KN83
NLSNA
>O15439
FETAL
>Q9HCH3
LHTHI
>P0DP71
RALGI
>P02743
PLVWV
>Q9UKV8
TMYFA
>F2Z3F1
KEILL
>Q8WVQ1
GIEFI
>Q9P2R7
FQLPI

>Q15291
ISELL
>Q15293
NHDEL
>O60462
CCSEA
>Q9BZC7
TDTLC
>Q6IF99
TISLL
>P10155
TLDMI
>O60469
SYTLV
>Q9UHB4
TETWA
>Q8NCK7
LDTTC
>Q7Z6I5
IHTHL
>Q7Z6I6
KGEGL
>Q8N9T2
RKIVV
>Q9NUX5
AEDVI
>C9JH25
DTIDL
>O95352
DDETI
>P46821
CKIEL
>P28906
ADTEL
>P28907
CTSEI
>Q8IXM2
NFDQA
>O15444
ANSGL
>Q8WX93
ESEDL
>Q8WX94
CDFFC
>Q01814
LETSL
>Q9UKW6
QEDKL
>Q9P2S2
KEYYV
>P15374
ALSAA
>Q9HCI7
MRFDC

>Q7Z353
VSESL
>P20674
GLDKV
>Q9NRD0
GHVAA
>Q9NRD8
HYENF
>O95813
PGVSA
>P59901
PWEQI
>Q9NRD9
HYENF
>P56880
LKDYV
>Q86U28
FSIKL
>Q9UHC3
LVTQL
>Q96NU0
KKEEC
>Q9UPS8
KNYMI
>Q8IU99
YFSKV
>Q5D0E6
PLSHI
>Q9UHC7
YDLDL
>Q86SK9
GDSSA
>Q15760
PNTFV
>Q8NCL4
LWLFV
>Q01362
PPIDL
>Q8TCT6
RFLEV
>Q15762
PKTRV
>Q7Z6J0
FVENI
>P51677
LSIVF
>Q8NCL8
TSTIF
>Q7Z6J2
EESQL
>Q6TCH7
YVSHL
>P10620
SKLYL

>P51679
LHDAL
>Q5VVS0
RKETA
>Q15768
IYYKV
>O60938
QAVII
>Q99567
NHVNF
>Q99569
PDSWV
>Q96C12
ARVPA
>Q9HCJ0
SGESL
>P04000
SVSPA
>P04001
SVSPA
>Q6UXR8
SILIC
>Q9HCJ2
QETQI
>P18405
IPFLF
>Q9P2T1
FSEAC
>Q14449
ARIAL
>P02765
RHFKV
>P02768
AALGL
>Q9BZE2
IKSII
>Q9H9S0
QPEDV
>Q6DN12
KRSAL
>Q9H9S3
GALFF
>Q9H1C0
QDSAL
>O95822
KNSKL
>Q6P5W5
DDITF
>O95825
QVVQF
>Q5SRH9
SSVSL
>Q93034
FIYMA

>Q9UHD0
VMFSA
>Q9UPT5
FDTSA
>Q9UHD2
NVDCL
>A6NEM1
NITII
>P51681
ISVGL
>Q8NCM2
DEIHF
>Q5FWE3
DTIEL
>P51685
VDYIL
>Q63ZY6
QQFAF
>P51686
GALSL
>Q7Z6K1
EVTMI
>P09210
KIFRF
>Q15776
ESISV
>P10632
CFIPV
>Q9NUZ1
NQERC
>Q99574
DFEEL
>Q99575
IAIEV
>Q8N9V2
LNSHV
>Q99576
GGSAV
>Q6UXS0
DSILI
>O95377
KKTIL
>O95379
DEENI
>O15466
TCSCC
>Q96KB5
LETDV
>Q96C23
KFSVA
>Q01831
PFEQL
>Q96C24
QKLGL

>Q14451
TRVAL
>P02771
AALGV
>Q9HCK5
TMYFA
>Q00056
VPSSI
>Q4VX62
PASTC
>Q00059
GAEEC
>Q9NRF2
QYSFV
>Q9NZV7
ECDHC
>Q9NZV8
RVSAL
>Q9H1D0
WEYQI
>O95832
GKDYV
>Q9BZF9
QGLVC
>Q53RD9
SPYDF
>Q6P5X7
RFVKI
>P0C646
ISVCC
>O95835
DLVYV
>O95838
EESEI
>O60499
LLFSL
>Q9H1D9
EWLEF
>Q9UPU3
NCTSV
>Q14914
TIVKA
>Q14916
QHTRL
>Q13136
RTYSC
>Q9UPU9
RTSTI
>Q9UHE8
ICSQL
>Q86SM8
VDIAA
>Q96NW7
RELTV

>A0A0A0MT76
KVTVL
>Q7Z6L1
GPVCC
>P0DSO3
KQSQC
>Q86Z02
QYSYL
>O14602
DIDDI
>P10644
VSLSV
>P41181
RGTKA
>O00206
EATSI
>O14609
TDFSF
>Q8N9W4
LQSSL
>Q8N9W5
SKTGV
>Q09161
CALQA
>Q96C34
IKDAF
>Q96KC9
DEFMI
>Q9HCL2
SFVVL
>Q9P2V4
NEYFC
>Q9NRG1
EKYRV
>Q9NRG7
KEIVA
>Q8N6C5
LPVPI
>Q9NRG9
PHSHL
>Q6NUJ1
AGEHA
>O95847
GVSPF
>Q9UPV0
KVYRF
>Q93052
ASTDL
>Q6W5P4
KPEFI
>Q13145
KLEFV
>P15863
SELGF

>Q96FH0
PPSSA
>Q14929
PQEVF
>Q6XLA1
LPLPA
>Q96NX9
QLYSA
>A0A1B0GVH4
SSVGA
>Q7Z6M2
FSEDI
>Q7Z6M3
SELNF
>Q8IZ81
LTLKV
>Q5JVF3
LSTVC
>P09238
SWLHC
>P60891
SHVPL
>Q86XJ0
QHTDV
>Q9BUL8
FKTVA
>O15480
SSSHA
>Q8N1H7
QFTFF
>O95395
YGTEL
>Q10588
SRTQL
>O95397
ILDAL
>O95399
WKYCV
>P05814
NPISV
>Q9HCM4
LTTEL
>Q9P2W3
KCTIL
>Q9P2W7
PSVEI
>Q6ZWB6
QKYGL
>P06681
NFLPL
>Q9H1F0
CMSIL
>Q9BR84
LTECV

>Q9H9V4
LDELV
>Q8WVV9
TSSHL
>A0A1W2PPW3
KKIMF
>Q0Z7S8
IYEKV
>Q9UPW0
WDSIV
>O95859
EMEEL
>Q6NUK4
PQVYF
>P32927
PGEVC
>Q13151
GGSSF
>Q9UHG3
LKTEL
>Q00536
VDTEF
>B9EJG8
QTDQV
>Q86Z20
SSIIF
>Q5JVG2
QLSSI
>Q8N309
RMFAV
>Q8N1I0
KVSQL
>Q13613
VHTSV
>Q13614
VQTVV
>Q13616
YSYLA
>Q86XK3
EFIDV
>Q13617
YSYVA
>Q13618
YTYVA
>Q13619
YHYVA
>Q86XK7
GVVKA
>Q9Y3X0
PFESV
>P18440
RFFTI
>Q96C55
KGEPA

>Q96KE9
LIFYA
>Q9P2X0
RGLRF
>Q9HCN8
GHDEL
>Q9NZY2
QPLEL
>Q9NRI7
VALLL
>O95866
YAVVV
>Q5TZJ5
LWEGI
>Q93070
SKSRV
>Q9UPX0
ALSKL
>Q14940
RGSRL
>Q93075
RLYSL
>Q96H15
GLFTL
>Q7L8J4
RSVSL
>P15882
EDILF
>Q8NCQ2
QLLPL
>Q8NCQ3
CLYLA
>P0DKB5
PGSGL
>A2RTX5
AEEAF
>Q13620
YNYIA
>O00238
QDIKL
>O14638
FETTI
>Q9BUN5
EQSAA
>Q13625
QRSLA
>A6NL46
GGSRL
>Q6PIJ6
EDDYI
>P21108
SHVPL
>Q8IXS8
LISQV

>Q9Y3Y4
VGSDA
>F2Z3M2
VLFMF
>Q6NW40
LIVFL
>Q562E7
IRLLA
>Q5TCH4
DKDQL
>Q8N6F1
GPLGV
>Q9NRJ5
TYLIL
>Q9BZJ6
HRTVV
>O95872
MNLEF
>Q8N6F7
QFSHL
>Q4VNC1
NEEQL
>Q86U86
NLENV
>Q96PB8
ISTVV
>Q6NUM6
AISGC
>Q14952
EVSYA
>Q86SQ0
THFLL
>Q14953
EVSYA
>Q9UPY5
EEDKL
>Q14954
EVSYA
>Q14957
LESEV
>Q86SQ6
NETTV
>Q9UHI7
VCTKV
>Q86SQ7
PQSDC
>P54753
LPVQV
>A6NER3
KQSQC
>Q8NCR3
STYGL
>P54756
GMVPL

>Q66K64
TWIVL
>Q8NCR9
DGILF
>Q8N323
LNYIC
>Q66K66
PPVRV
>P85299
RQSVV
>Q8N326
SYFLV
>Q9H6D8
NTIDV
>Q6P2H8
NCVRC
>O14649
RRSSV
>Q03112
SISHV
>Q96SW2
VILCL
>P49914
DSSTA
>Q7RTR0
RGVLL
>P49917
NQYLI
>Q8IXT5
KLTLL
>Q6UXX5
LSYVL
>Q07002
RQSIF
>Q7RTR8
NALPL
>Q9P2Z0
KEETC
>P36382
DDLSV
>P36383
TSVWI
>Q5JS54
FPEKF
>Q96AP7
AGSLV
>Q9H9Y2
RKFHL
>Q6DN72
EEVLC
>Q9NRK6
SFISA
>Q86U90
HASYL

>O95881
LEDEL
>Q9BZK8
FVSHC
>Q9UJ14
GATIL
>Q9H9Y6
KLDVV
>O95886
AQTRL
>Q93091
LDSIL
>A0A1B0GX31
CASSL
>Q6ZMN7
SVTTV
>Q7L8L6
FTSAL
>A8MVJ9
DQFAA
>Q68D06
LYIFL
>P54764
RMVPV
>Q66K74
CKVEF
>Q9H6E4
SQSEL
>Q58EX2
FSSFV
>O00257
EYVTV
>Q13641
SNSDV
>P31639
WGFYA
>Q9UMF0
QLTSA
>P35523
DELIL
>Q8N1L9
AQVHF
>P21127
FSLKF
>Q8WXA2
CNEDL
>Q7RTS5
VYLGA
>Q9H310
ADTQA
>Q8WXA9
KTEAV
>Q96AQ1
HRSVL

>Q9H313
WQSSI
>Q96AQ2
IASLI
>Q9NRL2
KKSRI
>Q9NRL3
AKVFV
>Q96AQ8
YRLWI
>Q9BZL6
RISVL
>Q96PD2
FKEIL
>Q13190
VVFLA
>Q13191
PRLNL
>Q5T9S5
QGENV
>Q9BPU9
YGVEC
>Q52WX2
IEICV
>Q8NCT1
VSFIL
>P22459
VETDV
>Q8NCT3
LKDLI
>Q9NWH7
TQESF
>Q66K89
QTVIV
>O00267
KLLEA
>P48146
SILRC
>Q96SY0
STERI
>Q9BUQ8
ETIFA
>Q6UXZ0
SETAL
>P14598
LASAV
>Q8NHP1
PNYFI
>Q8WXB1
QKEDL
>P04080
ELTYF
>Q8IXV7
LRDGV

>O75334
RTYSC
>P0DPA2
NGLLV
>Q8N807
VKEEL
>Q9BZM1
EKTDL
>Q9NRM1
LLLQA
>Q9NRM2
NQLLL
>Q9BZM3
EISPL
>Q9H329
LMTEL
>Q0VGL1
EPIDV
>Q7LBE3
TLTAL
>P08F94
IQEQL
>Q9NRM7
QPVYV
>A8MX34
KATLC
>P59990
TPSCC
>P59991
ISSCC
>Q96PE1
SETTV
>P17152
ELYAV
>Q8TED1
KKEDL
>Q8IUB3
CMSIL
>A0A1B0GX51
CASSL
>Q96PE7
ELEQA
>P50895
FGDEC
>Q14982
FFIKF
>P22460
RETDL
>P35080
RDSGF
>Q9UHL4
PRLSL
>Q00587
DEVKV

>Q96FN4
NSEPA
>A0A1B0GVN3
SSSCV
>Q3B7I2
NHTEL
>Q9BW92
AEEIF
>Q96FN9
HLIEF
>Q8N350
DHSPA
>Q8NCU7
ALLLL
>Q8NCU8
KLDLA
>P17612
EFSEF
>P62191
EGLYL
>Q5SWL7
LHLCC
>Q9BUR4
VGELI
>Q86XP1
PQSEV
>Q8N1N2
PTDHL
>A6NL88
NEVTV
>Q86XP6
ADIHV
>O75340
VFSIV
>P25024
VSSNL
>P07108
KKYGI
>P25025
TSTTL
>O75342
NSISI
>Q8NHQ8
EGIYV
>Q8N813
RLFEC
>O75347
VKLEA
>Q8NHQ9
LEDDC
>P0DPB6
NESTF
>Q9NRN5
KEEEV

>Q9H1L0
GWSHC
>Q9NRN9
IRFSF
>Q86SU0
RSVVI
>Q9P107
AEDHL
>Q9BPW5
TVTSV
>Q00597
LRTQV
>Q14997
PCYYA
>H3BUK9
TKTSI
>P40394
TVLTF
>Q12802
EEIFC
>Q12805
GPFSF
>Q8N365
HLLNL
>Q68BL7
LHFVV
>Q96M20
RELLA
>P48165
DDLTV
>P48167
WSIYL
>Q9Y5E2
NNLGF
>A6NL99
ALEHF
>Q7RTV2
KIFRF
>Q8IXX5
FSLRA
>Q7RTV5
SVIHV
>Q8WXD2
IYSSL
>Q9H340
PHSRA
>O00743
TPYFL
>Q9H343
HKSQA
>Q9H344
HHIKI
>Q9H346
GKTSI

>O00748
RHTEL
>P27695
LYLAL
>Q9Y216
VFLTA
>Q6ZMR3
KELKL
>Q6NUR6
KTESL
>Q6ZMR5
SKTGI
>Q8IUD6
KQVKV
>Q68D42
HETIV
>Q9NY26
LFIQI
>A0A1B0GX78
CASGL
>Q6P444
LNSRI
>Q8NCW0
LISAL
>Q8N377
SLLGF
>Q08379
KITVI
>Q9Y5F1
NNLGF
>Q9Y5F2
FGFNF
>P49961
WKDMV
>P60014
QKSSC
>Q86XR7
RQFIA
>Q03167
SSSTA
>Q8NHS1
FPVCL
>Q8NHS2
IGIKL
>Q9HCU0
CRTSV
>Q3YEC7
DYEEL
>Q53T59
APSLF
>Q9HCU8
HLYPL
>P0DPD7
SAIQL

>Q6UW01
LIFPL
>P48637
NPYPV
>P17181
QQDFV
>P39900
SWFGC
>Q96PH1
FQENF
>Q9P121
LLLKF
>Q9Y226
SSTYF
>Q9Y228
DNLMI
>Q9NY37
RIEEC
>P36894
QDVKI
>Q9H814
DLDIF
>P36896
EDVKI
>Q05DH4
EDSCC
>Q92733
AKYGF
>P61353
QKLRF
>Q92738
ESVLL
>Q9NWL6
KETKL
>P17643
NQSVV
>Q8N387
LRTSV
>Q8IZA0
REEIL
>P35573
TLYDL
>Q86XS8
EVEWF
>P25054
LVTSV
>P78310
DGSIV
>O75373
TGEKL
>Q8WXF3
ISSLC
>P40879
VETKF

>P78317
HPIYI
>Q07065
IHEKV
>P0DPE3
ALDGL
>O00764
QATVL
>Q96R08
IGFIF
>Q9BZQ6
EKDEL
>O43502
PEEEL
>Q9Y232
KIDEF
>Q16254
PVLNL
>Q2WGJ8
SSLEV
>Q6ZMT1
ALTEI
>O75830
DLDSL
>Q6ZMT4
ARFFV
>Q9Y239
RIICF
>P11117
GEDHA
>A0A1B0GX95
CASSL
>O75838
FHIRI
>P29401
LITKA
>Q9UHP7
SDIHV
>Q5THK1
QSSGC
>Q9H825
RTLFI
>Q7Z6W1
FTSPI
>Q9NWM8
KHDEL
>P52630
MPSDF
>Q9BUV0
LWIPI
>P17655
CFSVL
>P17658
MLTEV

>Q8TB45
EELEC
>Q86XT4
QPTKL
>Q7RTY1
VASNV
>Q86XT9
SRTRL
>Q96KN8
KPITA
>Q96KN9
KSEWV
>Q5VU36
LWEGI
>Q7RTY8
VPSLL
>O75385
TGICA
>O75387
SEVTA
>Q8WXG9
ADTHL
>Q9NT99
QETQI
>A8MX80
GGSRL
>Q969G5
MESVA
>Q9BRB3
GEVAL
>Q9Y240
CEFPF
>Q8N6N7
EKYGI
>Q8NEA5
KMEDL
>Q9H832
GSLRV
>Q9NY59
GEEEA
>Q86SY8
CTVLF
>Q8WU39
TREEL
>P61371
SPIEA
>P0DUB6
AESKL
>Q92752
QSLQF
>Q96FS4
TADLA
>Q16720
VETSL

>P17661
QHEVL
>Q96M66
RSLAV
>P35590
TAEEA
>P42127
LSLNC
>Q9Y5I7
VDTRV
>A6NJV1
FEFRA
>P46013
DSEDI
>Q8WXH2
INFFI
>P78334
VCLNL
>O75398
EKVTV
>Q96R28
IMYIA
>Q96AX9
IQIFV
>Q3ZCM7
EEEVA
>Q9UJ90
GAERV
>Q9Y250
IATEI
>Q969H8
SRTEL
>P60508
QESPF
>Q86UA6
WAVIL
>Q9UJ98
DIEDF
>Q5VZ03
QNFSV
>Q8TEJ3
FVESF
>Q8NEB5
YCFDI
>Q9NY65
EGEEF
>A6NGB7
DSTLV
>A0A1B0GVT2
KSTMV
>Q96FT7
EDFAC
>Q16739
EILDV

>Q6P2Q9
EDLYA
>P17677
DQEHA
>Q6ZRR7
RNSPV
>Q9Y5J1
HYSDF
>Q8TB68
RTTAV
>P33402
RETSL
>Q8NHW3
ADFFL
>A6NJW4
LSTVV
>Q8WXI2
IETHV
>Q5VU57
PSTPF
>Q7Z3E1
NTVSI
>P78348
EDFTC
>Q6PP77
RQSVV
>Q8N878
QEFVV
>Q15418
PSTTL
>Q96AY3
VHEEL
>Q96AY4
PSSKC
>P0DPH9
ETSVL
>Q969I3
NLVPF
>P60510
ADYFL
>Q9H1R3
MALGV
>Q9Y264
RPLDI
>P53985
EESPV
>Q9Y267
SSDDV
>O75864
GQETL
>Q5H9K5
SILGF
>Q9NY74
PTSFL

>Q8IUI8
KVLVF
>Q8NEC7
VGVPI
>Q12860
VYLEF
>Q5DX21
AGSLV
>Q9H6N6
GITSV
>Q6ZRS4
ADVFL
>Q6PIU1
DDFWF
>Q71DI3
RGERA
>Q2T9J0
PRSKL
>P25092
ESTYF
>Q96KQ4
QRTLA
>P47813
DIDDI
>P78352
ARERL
>Q08881
AESGL
>Q5VU69
YVERF
>Q8N884
VFDEF
>Q8WXJ9
LNLEI
>Q969J3
RELRL
>P60520
NTFGF
>O43543
GVEFC
>O43548
VDFAL
>P12931
PGENL
>Q9Y277
FELEA
>Q9Y279
GKSVC
>A0JP26
TKTSI
>O15118
RLLNF
>O75879
KKLSL

>P0CE67
DASHL
>P34741
KEFYA
>Q9NWQ8
DITRL
>Q12879
IESDV
>Q2NKJ3
LASSC
>Q8IZF0
DLLDI
>Q6E213
ELEII
>Q96M95
QRVRI
>Q6ZRT6
RLFQA
>Q6PIV2
FLFDL
>Q96M96
KKSEC
>Q8IZF7
HEDVL
>Q9Y5L3
LPSTI
>Q86XX4
DGTEV
>P01100
TLLAL
>Q96KR1
DYDNF
>P01106
RNSCA
>Q8NHY2
VLELV
>Q8NHY3
EESWV
>P26885
RRTEL
>A6NJY4
EKLDL
>Q15431
EKLFV
>P20809
LKTRL
>Q15435
TFVRF
>Q8N895
VLLGI
>Q15436
VSSAA
>Q15437
VSSAC

>Q96R54
EAYFI
>Q9BZV2
MSTKL
>Q9BZV3
QVEEV
>Q969K4
IGLDC
>Q6UW63
TKDEL
>Q969K7
EPLTL
>Q86UD0
DSTFI
>Q14C87
LHENV
>Q86UD3
KETVV
>O43557
GAFMV
>Q86UD5
TSVQV
>O43559
TDLPL
>Q9Y285
TQEAA
>O75882
PGTCI
>Q9Y289
QETSL
>P11166
ADSQV
>A8MVU1
LASAV
>Q9NY91
HGYYA
>Q9UHU1
GTINL
>O75884
LKVPA
>Q8IUK5
EAEQC
>P11168
ATETV
>P11169
TTTNV
>Q8IUK8
LVFPL
>O75888
GFVKL
>Q8NEE8
YYEAV
>Q8WU76
PDLGF

>P30872
RITTL
>Q9H871
KQIFF
>P51808
IAIVL
>Q7L8W6
YIYNF
>Q9NY97
AHLKC
>P30874
LQTSI
>Q92791
EPELA
>Q92793
FVEGL
>Q92796
SPEKL
>Q16760
PAVEA
>Q16762
KSEKA
>Q6IC83
GLLRL
>Q12889
VDEEA
>Q16769
EYLHL
>Q8N1W1
NIVYL
>Q8TB92
ASFNA
>C0HM01
ELSPC
>Q8NB14
GRLVF
>Q96KS9
RFSLC
>Q8NHZ8
GSLEF
>O60613
KLERI
>P10301
PCVLL
>P0DPK5
GNLAL
>Q6AWC2
PADDV
>A0A075B6H9
WGTGI
>Q06710
AFDHL
>P16389
MLTDV

>Q96R69
SSVKF
>Q8IW35
VGVTV
>O43566
TDSAL
>Q9H1U4
HNYKA
>O15131
DGFQL
>P11172
SRLGV
>P51812
TSTAL
>P11177
KTLNI
>O95049
PATDL
>O75896
ILYEV
>O75897
FHFQF
>O75899
MVSGL
>Q96FX2
ELVKC
>Q9NWS1
QFFRL
>Q12894
RADIL
>Q96FX8
FYTSA
>Q16772
KIFRF
>Q16773
WKVEL
>Q9NWS8
GRVFF
>O95500
LNDYV
>Q16774
QRTGA
>Q9H6Q3
SLDDA
>Q6ZRV3
GALGF
>Q8NB25
RYFTF
>P63119
IGYPF
>Q9UMR7
MKIHL
>P46060
TLYKV

>Q96CD0
PTLVA
>P01127
ETLGA
>P46063
KIDDA
>P15529
KFTSL
>P78382
RVIGV
>A0A075B6I0
MGSGI
>A0A075B6I4
SSLSA
>A0A075B6I6
NSLNA
>A0A075B6I7
FISGL
>Q9NRX1
SADRF
>Q969M3
LISVF
>O43572
KSTKL
>Q86UF4
KFLPL
>Q685J3
MTTSF
>A0A087WV53
RVDVA
>Q9H1V8
PESEL
>Q8NEG0
YTIEI
>O15146
GTVSV
>Q96PP8
PCVLL
>Q96PP9
LGSRI
>Q15911
DTFRL
>Q8NEG5
EGVQL
>Q15916
RQSLL
>Q99712
QQSNV
>A0A1B0GVY4
EAVEL
>Q9BWD3
EDEDF
>P63120
IGYPF

>P63121
IGYPF
>P63122
IGYPF
>P63123
TGYPF
>P63124
IGYPF
>A6NLC5
IETTV
>P63125
IGYPF
>Q9UMS4
KFYSL
>P63127
IGYPF
>P01130
EDDVA
>Q8IZI9
GDLCV
>A6NLC8
KKIMF
>P63129
IGYPF
>Q06278
WNVPI
>P01135
SETVV
>A0A075B6J2
SSYTF
>O60636
SRDVI
>O60637
GGTYA
>Q9BT09
PPDEL
>A0A075B6J9
SSSTF
>A4D126
QLLIA
>O43581
HQLKA
>Q70Z53
QDLFL
>Q7L211
NVTII
>O43586
YLEKL
>Q86UG4
EETDL
>O15151
KVFIA
>Q6GMV2
EMTDV

>Q96HA4
RISNV
>Q96PQ7
VTVKL
>Q9HB19
RTSDV
>Q9UHX1
SDLSA
>A6NGH7
TGDFF
>Q14146
KRYTA
>P36021
PEEPI
>Q16790
AETGA
>O95521
LHLCC
>Q16795
KTVNI
>Q8IZJ0
GDLCV
>P63131
IGYPF
>Q8WZ60
GAVSV
>Q9Y5P0
GQSRA
>Q14602
LMSLL
>Q8NB42
NRETL
>P63135
VTVSV
>Q9Y5P4
KPILF
>Q5T6L9
SKVLL
>P0CJ69
LSSVF
>Q53EL9
GDERI
>Q9Y5P8
DLEPL
>P46089
SPSDV
>O60641
IKDFL
>Q5VST6
ELVNL
>Q7LBR1
LRDQV
>Q9H1X1
LPFML

>Q9HB21
PVSDV
>Q8NG06
RDDHL
>P0DW11
KRIMF
>Q96PR1
IPSIL
>P0DW12
KRIMF
>P0DW13
KRIMF
>P0DW14
KRIMF
>P51843
LCTKI
>P16870
ETLNF
>P15090
VYERA
>Q01538
RGIQV
>H7C350
GFLPF
>Q9NWV4
QFVKC
>Q9BWF3
RYSAF
>Q9H6T0
EWVCL
>Q5T848
DSFKV
>Q6NZY7
DVIGL
>Q9Y5Q3
PEFFL
>P47871
AESPF
>Q9Y5Q8
ILDYV
>P47872
RTSII
>P46095
SPSEV
>A0A075B6L6
RASSL
>P58304
LEDMA
>Q8IW70
VETSL
>Q9BRK0
GGDSA
>Q53RY4
QRSWV

>Q969P5
NLFKF
>Q969P6
EDFEF
>A8TX70
EATDI
>Q9BRK4
TATEI
>Q9BRK5
VHEEF
>Q9UJA9
PLLQA
>Q8NEJ0
LMIPL
>Q96HC4
HSVNF
>P0DW28
VTSSC
>Q5H9R4
LISKL
>Q8TER5
PTTPL
>P51858
DHESL
>Q9NWW6
LQVTA
>Q8N3C7
VDENC
>Q9Y5R2
VQEWV
>P0CJ85
LLEEL
>Q14624
WSVEL
>Q09328
CKDCL
>P0CJ86
LLEEL
>P0CJ87
LLEEL
>Q8NB66
TEESA
>P0CJ88
LLEEL
>P0CJ89
LLEEL
>P47881
RRSLA
>P29965
GLLKL
>Q96CH1
QQVLA
>P36507
TRTAV

>P36508
SESGC
>Q8WXQ3
YSEYV
>Q52LC2
SKIYV
>P47887
NPFLL
>Q01094
TPLDF
>O60662
KLSKL
>Q86W11
THTRV
>Q02878
HKLVF
>Q969Q0
QVIQF
>O60667
INVPA
>O60669
RETNI
>Q13304
AKSEL
>A6NI28
YVVFL
>Q13309
KPSCL
>Q8IUQ4
TISMC
>A6NGK3
KQSQC
>P33947
LSLPA
>Q5SYB0
ASTAL
>P23435
LVFPL
>P68371
EEEVA
>Q9NWX5
VEDDI
>O95551
LDIIL
>P11686
PLYYI
>P59646
SATTC
>P0CJ90
LLEEL
>P63165
GHSTV
>Q8IZM8
DDESA

>Q9UEG4
PGVSL
>Q9Y5S8
NKENF
>Q6PXP3
KETSF
>A0A075B6N1
CASSI
>O60673
LLDQF
>A0A075B6N3
ATSDL
>O60676
KCEDA
>Q9BT49
SSSMA
>Q86W26
KNTYI
>Q8N6Y0
GDTFL
>Q8N6Y1
DISNI
>Q8NEL0
EVTRL
>O15197
GSVEV
>Q14181
QVVRI
>Q8IUR6
PTSKV
>Q4AE62
PREDL
>Q8IUR7
QQYLA
>Q6UUV7
RADRL
>Q14183
ALSSA
>Q6UUV9
RMDRL
>Q8NEL9
NLDPI
>Q9BY07
WSYSL
>P18146
TIEIC
>Q9NWY4
DQLAA
>Q99767
TPLYI
>P62308
ALERV
>P55769
ERLLV

>P06396
AELAA
>P27338
LLVRV
>Q8IZN3
KLSSV
>P63172
FGLSI
>P04201
VETVV
>Q9UMX5
IKDEF
>P49146
EATNV
>Q5TEC6
RGERA
>Q8WXS5
KTTPV
>O60682
CGTTA
>O60684
EGFQL
>Q86W33
KAINA
>P58335
KELTA
>Q969S3
VQVRF
>Q8NG41
DCLAA
>Q13322
IRVAL
>Q969S8
PHLVA
>Q13325
LRLSI
>Q13326
NHICL
>Q9BRN9
GSLYI
>Q96HF1
RKLQC
>Q96PV6
QLSAF
>Q8NEM2
GTFLF
>A0A1B0GXF2
CASSL
>Q9NYA1
PEEPL
>Q9BY11
YVEAI
>Q8TEU7
QVSAV

>P98066
RFSHL
>P51888
QSVVI
>P55771
TASAL
>P55774
LKLNA
>Q9BWJ2
EPTSF
>Q6NT46
KQSQC
>Q9NWZ8
IPLKF
>Q0VDI3
VRIDV
>P00325
TVLTF
>P00326
TVLTF
>Q5XX13
QNVFI
>P59665
WAFCC
>P59666
WAFCC
>Q8NB91
KLSNL
>Q53EQ6
PYDGV
>P18615
LVDGF
>Q9Y5U5
GDLWV
>Q2V2M9
SELQL
>P19484
EGDVL
>Q86W42
FSLSF
>P31321
ISLTV
>Q9HB71
GDTEF
>Q13336
VESPL
>Q96HG1
LQVYI
>Q9Y2A7
VTSSA
>Q9P1A6
AQTRL
>A6NGN4
DQYCC

>Q9BY21
DYTDV
>Q9NYB5
KETQL
>P05543
NPTEA
>O75015
VKTNI
>Q8TEV9
FLYKI
>Q04609
LSEVA
>Q8NEN9
PSESV
>O14807
QCVIL
>Q99784
RSDEL
>Q9H009
MELTV
>P0DKX4
ETEVV
>Q9H6Y7
PVILV
>Q8N3G9
KTYTV
>P00338
KELQF
>A6NLJ0
ALLLL
>Q96MC6
QDTNV
>Q8TBB1
PGTFL
>Q9Y5V0
ADVQA
>Q9UMZ2
LPDLL
>Q00266
RKLVF
>P18627
EPEQL
>A0A0B4J238
SYLCA
>P36544
SKDFA
>Q6IPM2
KNFPV
>Q9NTG1
VYLVV
>Q52LG2
CRSTC
>Q70SY1
VNTTF

>Q8N8C0
LLFLI
>P61006
RCVLL
>A0A1W2PRV1
KRIMF
>P27816
QETSI
>Q5JSJ4
SRSSC
>Q00722
QESRL
>Q9BRP4
QLSDL
>Q9Y2B0
SHDEL
>Q9UJF2
KNSSC
>P35221
AMDSI
>Q9Y2B2
SLSFL
>P35222
FDTDL
>Q9BRP8
LELGL
>Q13347
FEFEA
>Q96PX1
GPLAV
>Q8NG68
AFIKL
>Q6QNK2
DLSAV
>P54922
TVISL
>Q96HH4
ETSTV
>P35228
EMSAL
>Q9NQ31
LVFPV
>Q9BY32
GSLAA
>Q68DE3
CSSAV
>P23470
MESLV
>Q6P4E1
FNDVL
>Q9NQ35
NTSEA
>P23471
LESLV

>P09430
YRSHL
>Q9NYC9
LLLQI
>Q8WUA8
GPTIL
>Q9H013
FLVPA
>Q9H015
LITAF
>O14817
DTYCA
>Q9BWL3
LESTL
>P50583
CSIEA
>Q8TBC3
NETSF
>Q96CM8
RHLNL
>P06881
RDLQA
>Q08043
GESDL
>P17302
DDLEI
>Q969V1
LKSHF
>A6NI72
LASAV
>Q969V6
WDSCL
>P22612
EFSEF
>Q8NG78
KRTFL
>Q9Y2C5
TFTHL
>P33993
RITFV
>P19961
AESKL
>Q9BY42
TSYCF
>P98095
TTFAL
>P0DUQ1
DQYCC
>Q8NEP7
ILDFI
>Q9BY44
LELGI
>P0DUQ2
DQYCC

>Q8NEP9
MVLPL
>O75038
VLLRL
>O00421
HSTEV
>Q6P4F7
KPVDL
>O14827
PRLPA
>Q96ME1
YCFVI
>B7Z1M9
THLSL
>A1L0T0
GSIAV
>Q2M385
GQSPA
>P53618
KKTSI
>Q7L5N1
RGLFF
>Q14684
AMDFF
>Q9Y5X5
NSSEI
>A2A3L6
ICTIV
>P49189
VESAF
>Q96CN7
LLSKV
>Q9NTI7
TAVWV
>P09917
NSVAI
>P0C874
LNLSI
>P0C875
LKLEL
>Q96RA2
AASCL
>Q6UWB1
PQVLA
>Q5BKX6
TESVV
>Q9BRR3
HPSLL
>Q13363
ASDQL
>P35240
FFEEL
>A6NI86
NITII

>Q86UP3
DMFSV
>Q9Y2D2
NPTKA
>Q96PZ2
SDEDL
>Q96PZ7
VCTVV
>Q09MP3
LKENF
>Q149M9
VCLIV
>O14832
ERTNL
>Q13822
YESEI
>P30039
GTLTA
>Q8N3J6
KEYFI
>Q8N3J9
KISVI
>Q96MF2
FLEEI
>Q9Y5Y0
SESAI
>O75503
TLSGL
>Q8IZS7
VRFNI
>Q9NV12
KLESC
>Q9Y5Y3
NQSAV
>Q14694
RVDLL
>P49190
TEDVL
>Q14696
KREDL
>Q9Y5Y6
ENTGV
>O75509
LPDLL
>P08138
ATSPV
>Q9UL01
SQSQC
>Q8N8F6
LLTIF
>Q9H3H5
LFYDV
>Q86UQ4
HHLPI

>Q9Y2E8
EQELL
>Q9NYF0
LMTTV
>O75052
DEIAV
>O75056
EEFYA
>P30040
EKEEL
>O14841
AQEAV
>Q9H040
SEESL
>P57058
VKTQC
>Q9NQ66
FDTPL
>Q9BY67
KEYFI
>O00443
AATYL
>P30044
IISQL
>B7Z368
ELELF
>Q6P4H8
LPIQA
>P30046
VMTFL
>Q13835
FTSRF
>Q53EV4
EQSLI
>Q8TBF5
GHFSL
>Q9NV23
SISNF
>Q9Y5Z6
KHLRC
>P43119
ACSLC
>Q9NV29
RSLFA
>Q5VUE5
SYFYV
>C9J6K1
PNEEA
>P62826
EDDDL
>P62829
AGSIA
>Q86W92
EDSNV

>Q9UL18
TMYFA
>Q502W6
VPETL
>Q9BRT3
PCVIL
>Q96RC9
KRVLF
>A0A0C4DH59
CASSL
>Q6UWD8
TSVGF
>Q9Y2F9
LIFYA
>Q9NYG2
YQYVV
>O75063
PLSHL
>Q04656
DDTAL
>Q9NYG8
KGVPV
>Q9BY77
FKIKL
>O00451
LKLAL
>Q9BY78
LNVYL
>O94766
PAIEV
>O94768
SDLLC
>O00458
VGEFF
>P00387
RCFVF
>P25205
IIFLI
>O75521
RKSKL
>Q9NV31
FDLEA
>Q8IZU9
MQTHV
>Q8WZA6
SKSVA
>P08151
LNSSA
>Q9H511
VPVSI
>Q9NV39
LELQV
>P23975
HWLAI

>P62834
SCLLL
>Q6PH81
QRLIL
>Q5JU69
IAFFL
>A8MZ25
RSLHL
>Q6PH85
KRSLF
>P17342
HFSVA
>Q9UL26
KRSCC
>Q96RD6
STVEF
>Q969Z4
SNLVI
>Q9BRU2
RPYPI
>Q96RD7
LDSSC
>A8MXK1
EDVEC
>Q13394
SLEKL
>Q9Y2G3
DSSTC
>O96001
DKIAI
>Q9P1G2
CPSQA
>O96006
DSSFL
>O75071
VALGI
>B5MCN3
KMEKF
>Q8NET5
VYENL
>O75074
ALLVC
>Q8WUF5
QRSKV
>Q8NET8
PETSV
>Q8WUF8
KHEEL
>O00463
DLEDL
>P62380
RKEIL
>Q9H063
PVICI

>P09486
KDLVI
>Q8WZB0
DVSSL
>Q6B9Z1
RSDLI
>O75533
LDYIL
>Q5T6X5
RMSSI
>D6REC4
RLEAA
>Q6P9F7
CLDKC
>Q8N8I0
NSIQI
>P13473
GYEQF
>P09958
DQSAL
>P31391
RTTTF
>Q9NTM9
KNILV
>Q8IWB4
LWEGI
>Q9Y2H0
AQTRL
>Q6UWF7
LNYIC
>Q9Y2H2
RIIQI
>P04746
AESKL
>P49682
SYSGL
>Q86UT5
ASDLL
>P49683
VSVVI
>Q11203
LSSGI
>P49685
RSVSL
>Q11206
NLTSF
>Q92903
PTLKV
>P57081
ATLSC
>O96018
QPVYL
>Q9NQ90
QHTNV

>O75084
GETAV
>Q8NEU8
AESEA
>P57087
KSFII
>P09493
DMTSI
>O14874
ESFRI
>P17813
TSSMA
>O00476
KLTRL
>P48357
CDLTV
>Q8TD06
IQSEL
>P07307
TGEVA
>Q6P9G0
DLTEL
>E5RQL4
FFFFF
>P43146
TGSAF
>P30530
QEDGA
>Q8NBA8
NSVKI
>Q9NV58
IQTEI
>P62851
AGEDA
>Q6IPT4
SYFLF
>Q5JU85
ISTVV
>A2AJT9
VATGI
>Q9NTN9
DESSV
>Q96RF0
TVLSF
>Q96J65
AEVRL
>Q86UU0
ANLPF
>Q9Y2I2
SPLVF
>Q9Y2I6
AALSV
>Q9UJM8
AVSKI

>Q9Y2I7
LGLNC
>Q92911
QETNL
>Q8WUH1
MTLLF
>O75093
CALCA
>Q92918
GITGL
>Q7L0X2
INEEI
>Q9H8H0
VLELF
>A0A1W2PQ09
KKIMF
>O75096
SESQV
>Q8NEV8
KESEL
>Q8N567
KVVNF
>P52803
MLLTL
>Q0VF96
VASQI
>Q8TD10
MRTVI
>P60201
RGTKF
>Q5T8A7
SVVKV
>Q9BWS9
FYDLL
>C9J1S8
CCVHL
>Q96MK2
EITIF
>Q9NV66
AISGC
>P0CB47
IEVDV
>P0CB48
MEVDV
>Q6Y288
FREEL
>P20020
LETSL
>Q9UL51
LSSNL
>Q9UL52
SKTGI
>Q7LDG7
FDIHL

>Q5STR5
EELEL
>Q7Z3Y7
QRVPF
>P13498
TDEVV
>A8MZ59
LQSSV
>Q9Y2J4
VEILI
>Q9NYK1
FKETV
>P08648
ATSDA
>Q96HP8
EVSGI
>P16050
NSVAI
>Q13884
LGLVA
>Q13885
GEDEA
>Q9BWT7
SSSEA
>Q13888
APSGV
>Q13889
LKVSA
>Q8IZY2
AETVL
>P07327
TILMF
>Q8IZY5
SAEAL
>O75563
EMYDI
>P78504
MEYIV
>P43166
ASFRA
>P78508
RISNV
>Q6EIG7
NKIYL
>Q9H553
TKLLV
>P08195
FPYAA
>Q5VUJ5
PDECV
>P30559
QPSTA
>Q5VUJ6
QLSVA

>Q8N8L2
HTINI
>Q9UL62
VTTRL
>Q9UL63
DLITL
>Q7Z3Z3
RLFYL
>Q7Z3Z4
HLFYL
>A4D1B5
MLLGL
>Q9UL68
RGIQV
>P25705
AGFEA
>Q96J84
MQTHV
>Q6UWI4
PAVTV
>Q86UW1
LNLKA
>Q92930
RCSLL
>P22695
FVDEL
>Q86UW8
TPLHV
>Q5VZF2
PQTAC
>Q9HBA9
LSEVA
>Q9Y2K9
KWYQF
>Q8WUJ1
CSFPL
>Q6ZV65
KRTQA
>Q8NEX9
PADSV
>Q02505
TSSSV
>Q9H8J5
IYVDI
>Q15126
IRSRL
>Q3SYC2
HLEFC
>Q8N587
MSVTI
>P52823
SHESA
>P48380
TYTAV

>Q8TD33
SQDGA
>Q5T036
FLSPI
>Q5VW22
PDECV
>B6SEH8
WPLSL
>B6SEH9
EEFSL
>O75575
EEDPA
>Q9H560
LKSHL
>P03901
NLLQC
>P30566
AELCL
>Q92482
HKEQI
>Q6IPW1
ARISV
>Q9BTA0
RFTLC
>Q6UY11
KTTAL
>Q92485
CTLVL
>Q6UY13
IPSTL
>O43704
FRTEI
>O43707
GESDL
>O43708
TELRA
>Q9BTA9
NSFMV
>Q6UWJ1
RRSSL
>Q8IWF2
NKEEL
>P43631
EVSYA
>Q96J94
RLYYL
>P43632
EVSYA
>Q96RI8
FSEHI
>P0CG21
LKELA
>P0CG22
TPSRL

>Q6UWJ8
YQTLI
>Q8IWF9
MKSFL
>Q9Y2L5
IISNV
>Q9Y2L6
PGTLV
>Q86UX7
GHEAF
>Q569H4
TTTTV
>Q9Y2L8
EGSLL
>Q9HBB8
DDSYI
>P33121
STIKV
>Q6ZV70
FSVFV
>Q9Y2L9
NALSA
>Q5CZA5
QVSSL
>P61565
VTVSV
>P61566
VTVSV
>Q6ZV73
EGTIL
>P61567
VTVSV
>P16070
MKIGV
>Q15139
RVSIL
>Q9BWV1
PPLTI
>Q9BWV2
MNEQI
>P56715
EQEDL
>O43252
SLEKA
>Q6PCB5
DDILC
>Q6PCB7
GAFAL
>Q96MN2
TRVEI
>O75582
SDSVA
>Q9NV92
YFFLL

>Q2M3A8
QGEEF
>Q9UEU5
KQSQC
>Q9NV96
ADITI
>Q8NBE8
CVYNV
>A6NDE8
KQSQC
>Q96CW1
YETRC
>P62891
TKLGL
>A0A804HLA8
VCSFL
>Q96T17
LNTFC
>O43711
VTSLV
>Q6QAJ8
CKTVI
>Q6UY27
LPEGV
>Q8N8N7
EEISL
>O43719
DDDDI
>Q14CM0
KETTV
>P06028
RLIKA
>Q8NGA0
RLFPF
>Q9Y2M2
GCTIV
>A8MXQ7
PGTGL
>Q8NGA8
SSVKF
>P34910
PAELL
>Q8NEZ3
TEEEL
>P26599
SKSTI
>Q15147
PATVV
>O60318
DMVDI
>Q6ZTQ3
TETTV
>P35790
RKLGV

>Q9BWW7
SPVQA
>Q06418
PHSSC
>Q03395
DLSEA
>Q5JNZ3
ILTSA
>P51511
LQEWV
>P51512
MQEWV
>Q8NBF2
LRYVF
>P78536
KETEC
>Q9H583
LQSYF
>Q5VUM1
RCIDF
>Q96T21
MNLNL
>Q96T23
NSEQL
>P41002
GLVRL
>Q9BTC0
TASQA
>Q9H3Q1
DEIRV
>Q9H3Q3
PFLGA
>Q96RK1
GSVSC
>O95206
ANENV
>Q14CN2
LSTTI
>Q86WA8
LNSKL
>Q6UWL2
SFSAV
>Q86WA9
ALLKA
>O95208
NPFLL
>A4D1E9
KMDII
>Q6UWL6
LQTHV
>P43657
NESAA
>P61580
HFVMC

>Q9UBB5
SGDEA
>P08684
TVSGA
>P61581
HFVMC
>P61582
HFVMC
>P61586
GCLVL
>Q68DQ2
DIEIL
>O60320
VLDQV
>P56730
SVTKL
>P42331
PKTEA
>Q5JPB2
LVIEI
>Q59GN2
TKLGL
>Q8N3T6
LKDKA
>P78540
ARVRI
>A0A0A0MS05
CASSL
>Q15617
RRTLL
>Q96T37
ALTLL
>O43734
QVVPL
>Q8N8P6
CSVFL
>P0C024
ATSRL
>Q9Y466
KSSDI
>A4D1F6
RAIKF
>Q8NGC4
TPSEV
>Q8NGC5
VKLQC
>Q9UBC0
TCTKA
>Q8WW59
VPEGL
>Q92973
AFYGV
>Q02543
PNTFF

>Q9BQ04
RYSAF
>P0DMM9
FRSEL
>Q8N3U4
GVSMF
>A2RU67
APYEI
>P78559
CKIEF
>Q5QFB9
FFFFF
>Q9BTE0
SAEPC
>Q99424
WRSKL
>Q9BTE1
PLTQV
>Q99426
GLDEI
>Q9NTU7
LVFPL
>Q9H3S1
GTEVA
>Q8IY17
SATDA
>O95221
TSSFL
>Q9H3S3
QDSLL
>Q8N8Q9
NLTAF
>O43749
VVFSV
>O15315
TQLIF
>Q6UWN5
VGLSA
>Q9Y2P4
GEIKL
>Q9HBF5
TYFYF
>Q3YBM2
TAIVL
>Q9NYQ3
QFSRL
>Q15170
RPYPI
>Q5VZK9
EFIFV
>P0DMN0
FRSEL
>Q9NQA5
EVYHF

>Q9NYQ8
EEVMF
>Q9UQ74
LAIGI
>Q02556
QQITV
>Q9BQ16
HDVYI
>O60346
YDTPL
>P56750
STSYV
>O43291
NTYVL
>P42357
ESEDL
>Q8TD84
SYTLV
>O43296
QVSSL
>Q96EB1
KHLDF
>A6NDI0
CCIHF
>Q8NBI3
SFINV
>Q6J9G0
NYSML
>P03952
MQSPA
>Q9UEY8
EKVEA
>Q9BTF0
HSSGL
>O43752
LFLVL
>Q8N8R3
QPSSL
>Q8N8R5
VHEVF
>Q9H3T2
GRFNF
>O95231
TGDAF
>Q96RN1
SNEDV
>Q9Y484
DDDDF
>Q9Y485
VKFML
>P32302
SLTTF
>Q9Y487
DDSVA

>Q96RN5
CLSAA
>Q8NGE1
KIVKL
>Q8NGE2
KLLNL
>Q8IWK6
HETTV
>Q8NGE5
KLDVF
>Q9UJU3
DSVLF
>P50225
FRSEL
>A6NIE6
DTSSF
>P50226
FRSEL
>Q8NGE7
KNIIL
>Q09019
SGTVV
>A0A0U1RRN3
HVEGI
>A0A0K0K1C0
CASSL
>A0A0K0K1C4
CASSL
>Q9H8P0
LPFLF
>Q495D7
TCDRC
>Q3KQU3
VTEVL
>Q9UQ88
FSLKF
>O60359
RTTPV
>Q53GI3
GGSRL
>Q8TD94
FTTCL
>Q6NTE8
FDDDV
>P28331
EPSIC
>Q5T8I9
EQFEF
>P28335
RISSV
>P51553
RAVEA
>O60810
DQYCC

>Q8NBJ5
ARDEL
>O60811
LHLCC
>O60813
DQYCC
>P16581
PSYIL
>Q8NBJ9
KIYVF
>P83881
QVIQF
>O43763
LASVV
>P07858
YWEKI
>O95248
CLSDA
>Q5JSZ5
EESKA
>Q8IWL3
KKIPL
>P43694
DIITA
>Q9HBH0
LCLLL
>A8MXV4
RKSHL
>Q8NGF7
LDSVF
>Q9HBH9
VGDHA
>Q9UBF9
ESEEL
>Q6IMI6
FRTEI
>Q5TBA9
SGTSL
>P0DMP2
ECYGF
>Q02575
HVLDV
>Q02577
HVLDV
>Q68DU8
RKYHL
>Q86T13
GSSDA
>P58012
SRLDL
>O95707
GTIDL
>Q9H0A6
KILEC

>Q13002
KETMA
>Q96MT0
RLEEL
>Q96MT1
PVESA
>Q13007
KFYKL
>Q13009
LNTEI
>Q96MT4
SNEAF
>A0A578
CASSL
>Q15654
VTTDC
>P03973
SPVKA
>Q99453
KSSMF
>Q9NTX7
TVTEV
>Q99459
LKSKF
>P07864
KDLIF
>P46721
LKTKL
>O95258
KRLQI
>Q96RP8
LVTEV
>P11387
EDYEF
>Q96B02
HDDTC
>P11388
EDDLF
>Q14330
NSEML
>Q8NGG5
FKIPI
>Q9UJW3
LTSSL
>Q14332
GETTV
>P34972
DLSDC
>P02656
SAVAA
>Q9BYD1
EDYRL
>Q96HY7
AKTFA

>Q9BYD3
TPYHC
>P55916
RESPF
>A0A0K0K1E9
CASSL
>Q9BYD6
EKEDA
>O76015
TGSRF
>A1L429
KQSQC
>P24462
TVSGA
>Q86T26
SKTGL
>A6NNC1
ASFSF
>Q13011
TFSKL
>Q3LI58
CYSSC
>Q13015
ELDLL
>P51570
KVLCL
>A0A0B4J2E0
CASSL
>Q8NBL1
LKTEL
>P51571
SHIQA
>Q8NBL3
RDSTV
>A0A584
CASSL
>P23141
EHIEL
>P23142
SEYWF
>Q96T83
LEDNA
>P09104
NPSVL
>Q8N8U2
KIYEV
>Q1AE95
FSLRA
>P60763
KCTVF
>Q99469
VLENI
>Q71SY5
LMDLI

>Q86WG5
CISDA
>P14416
KILHC
>P27037
KESSL
>O15354
VGTHC
>Q96JA3
SDEVV
>Q8IWN6
DRVSF
>Q8IWN7
DDLDF
>Q6UWR7
LFLLA
>Q9Y2T3
FSSSV
>Q96B18
VMTTV
>Q9HBJ8
RLTPL
>Q9NYU1
THDEL
>Q9NYU2
KREEL
>P0DO97
SDENL
>Q9BQ50
PSLEA
>Q9BQ51
VNSAI
>Q99928
GYLYL
>O76024
FLSAA
>Q8WUS8
VILSL
>P0DMR2
SQDGA
>Q8N5A5
KMTEF
>O60384
NLYEC
>P0DMR3
RKLKC
>Q9BYE9
DTTDL
>Q3LI60
GGVWI
>O95727
PESIV
>Q5JXX7
ILIFF

>Q6DKJ4
KPEPI
>P40200
EMETL
>P51582
RADRL
>Q9NVA1
NDEGL
>A0A597
CASSL
>O60841
VFEII
>A0A599
CASSL
>Q8NBM8
VKTEL
>Q7Z5K2
YLEHC
>O60844
SCSRC
>Q8IY63
MEVLI
>Q8N8V4
VDTSL
>P42858
KVTTC
>Q5KU26
LSSAL
>O95273
SELEL
>O95274
AGVLL
>Q96JB5
MGTSL
>Q96B23
RKYLL
>Q9UBI1
RATQL
>Q9UBI6
TCIIL
>Q99932
RMTPF
>P34995
GLSHF
>P02679
PEDDL
>Q9BYF1
VQTSF
>Q538Z0
HSSPF
>Q6ZN03
GPTQI
>P34998
QSTAV

>Q9NYV6
QPSPL
>Q68DX3
STDIF
>Q71H61
MSLVV
>Q9BQ66
ASSCC
>Q9BQ67
RTISV
>O76039
KETAL
>Q5JPI3
TGDQV
>P59827
LVLSA
>Q3LI77
YQFTC
>Q6DKK2
NSVKL
>Q13033
AKVFV
>Q06495
NATRL
>Q5JPI9
PSLAF
>A6NNE9
RVTSV
>Q7Z5L2
DQSTV
>P24941
PHLRL
>Q7Z5L4
SSSDC
>Q86Y07
ALFFL
>O14508
YKFQV
>Q9H3Y0
KFTWF
>Q99487
HLSSL
>Q10472
LPEIF
>Q99489
PKSNL
>Q9HD33
KSSLV
>P28838
SQDNA
>Q96B33
CDSDL
>Q01740
FLIFL

>Q8NGJ4
IHTRF
>A8MXZ3
QPSCC
>Q9Y2V0
CHSIA
>Q5KSL6
SRSQL
>O15379
SDVEI
>Q9HBL6
DSSPA
>Q99942
WLLSI
>P02689
IYEKV
>A1L453
GWSVL
>Q9BYG4
PAVTL
>Q9BYG5
TIITL
>Q9NYW8
DVENL
>Q6ZN16
TKDKA
>P10092
RDLQA
>Q9H0E3
RKEKV
>Q6ZTZ1
TKSSV
>P0CL80
KQSQC
>P0CL81
KQSQC
>P0CL82
KQSQC
>P49321
ESTAC
>Q9UGF5
AMFKL
>P22303
RCSDL
>Q7Z745
TSIPL
>Q96EH5
TKLGL
>Q9NVC3
VDLLA
>Q7Z5M5
ICSDV
>Q6UB98
ELTPI

>Q6UB99
VLLPA
>O14514
LQTEV
>Q5TEZ4
PLEAA
>O00115
RAYKI
>Q8IY82
QKIFA
>Q8IY84
FCSIL
>Q8IY85
PNSKF
>Q9ULB1
KEYYV
>Q9BTL4
AVVAF
>O95292
GKIAL
>Q9HD43
HKLEV
>Q13508
LFVAL
>Q0VAK6
PKELA
>O15382
WMFPV
>P50281
LLDKV
>Q6UWU4
AFSEL
>Q9Y2W3
LLLNV
>Q14376
FGTQA
>Q9Y2W6
DDYLL
>P23634
LETSV
>Q99952
EWTRV
>Q9BYH1
YEVSI
>P0DMU2
HQTAC
>P0DMU3
GGTVA
>Q6ZN28
TSEEV
>O95754
DETSI
>Q9BQ87
CVLDL

>O95758
SKSTI
>O95759
KLSNL
>Q96G04
LNLTL
>Q14831
NNLVI
>Q14832
TTSSL
>P01374
GAFAL
>P01375
GIIAL
>Q587I9
KVLPV
>Q96MY7
LVSLA
>Q8TBX8
TNIFA
>O14520
ALEHF
>Q7Z5N4
FSSFV
>Q9BV47
QGLEA
>O14522
YLSSF
>Q86Y25
KAYEV
>Q86Y27
LCFIF
>Q86Y28
DGTAL
>O14526
YLVSC
>Q05193
PPFDL
>Q05195
ACLGL
>Q86WK6
TPIVV
>Q9ULC8
YEISV
>O15394
DDSKA
>Q96RU8
SSFFC
>P46776
CVLVA
>Q14CX7
KKLKI
>P46778
YEFMA

>Q6UWV6
LSEVA
>Q9UBL0
WQVKF
>O15399
LESEV
>A1A5B4
RSTDV
>P09603
VELPV
>A5YM69
LLSVL
>Q6ZVC0
WDTAI
>Q99966
FPSSC
>Q99967
SRVSC
>Q9H8W5
RTVAL
>C9J3I9
NSFSI
>P45452
SILWC
>P17017
MGEKV
>Q00444
SKEAL
>Q13064
FNLIL
>Q13066
KQSQC
>Q14847
YVEAI
>Q13069
KQSQC
>Q587J8
PVTRL
>Q8TBY0
QASFF
>Q2M3M2
WGYFA
>O60882
SWIGC
>Q8TBY8
GSSYC
>O60883
LGTPC
>Q504Q3
SVLAL
>Q13520
EMESV
>Q13523
IQEKI

>Q9ULD4
TSSYL
>Q9ULD6
FGLTL
>Q9ULD8
EGTGV
>Q8NGM1
ENIKL
>Q14392
QQYKA
>A6NIM6
WETAL
>Q6UWW8
AQEDL
>A8K010
TTTYC
>Q9Y2Y6
EQSAI
>Q9NYZ3
PLLKF
>Q9BYJ1
NSVSI
>O76071
RPEGL
>P09619
EDSFL
>P14920
PPSHL
>Q6ZVD7
PVINV
>Q6ZVD8
FDTAL
>Q9H0H0
SVSGI
>O76076
QNSAF
>O95772
PLLEL
>P0DMW3
LQVHI
>Q9H0H3
KHLPA
>Q13070
KQSQC
>Q13072
LCFIF
>P17028
NVVKV
>Q4LDE5
RRTGF
>P17029
LKSCV
>Q9UGI6
SSSSC

>A8MUI8
GSSHL
>Q9UGI9
DALGA
>Q8NBR6
SCVIL
>Q8TBZ9
APSRF
>O00141
TDSFL
>O60896
TDTLL
>O00144
NPTHL
>Q6V1P9
DEVQI
>P48023
GLYKL
>Q9ULE0
PADDV
>P49810
HQLYI
>Q9ULE4
KYFSF
>Q13536
GLDSL
>Q9ULE6
PEDLL
>P49815
FTEFV
>Q8IWT0
VIIDI
>Q8IWT1
PPSKV
>Q8IWT6
DKEQA
>Q14CZ8
VEISA
>O94910
LVTSL
>Q9UBN1
RTTPV
>Q9Y2Z4
KWLQL
>O94913
TVESV
>Q9UBN4
VTTRL
>O94915
VSTGF
>P09622
KSINF
>Q9UBN6
ATSCL

>Q9NS28
VAIWL
>P30203
DISAA
>Q9H205
AHSTL
>Q9P1Z9
QGLYV
>O76081
KSIEA
>P41587
ETSVI
>O76082
KSTAF
>Q6ZN54
PDVEA
>Q6ZVE7
TSSMV
>O76083
EGDCA
>Q86T90
DIFFI
>P00533
EFIGA
>O76087
KQSQC
>Q495M9
EDTEL
>Q9UI14
QMEPV
>Q86T96
FFFPF
>A0A1B0GW35
CPLPL
>Q14865
PSTKL
>Q13087
SKEEL
>Q8TDB8
TTTNV
>P18827
EEFYA
>Q5T0B9
RISLI
>Q96EL1
DVSYL
>Q6ZS17
ASTAF
>Q8NBS9
AKDEL
>Q7Z5Q5
PSFCL
>P62070
HCVIF

>P10599
INELV
>O14559
TRSYC
>Q9ULF5
FDIQF
>P48039
KVDSV
>Q6UWY2
PGEAA
>Q96B86
LPVFC
>Q8IWU9
QYLGI
>Q9H211
AEEGL
>Q9NS39
FLLTL
>P41594
SSSSL
>Q05682
SPTKV
>P41595
QVSYV
>Q9NQL2
PRVLL
>Q495N2
TGVHA
>P41597
DKEGA
>Q8N5H3
FHISL
>Q9UQB3
PDSWV
>O76095
QIESI
>Q6ZN66
HKLPF
>Q8N5H7
RSSEL
>P17041
QRLTL
>Q8TDC0
ESEEL
>Q96G42
LQTSL
>Q9C009
ETLLA
>P04435
CASSL
>P08311
METPL
>P26232
AMDSF

>P08319
TILIF
>Q9NVH2
AYTRF
>A2RUB6
FGSSF
>Q9NVH6
LGLQA
>O00160
YVEKI
>Q6P1J6
RTVAL
>P62081
PEFQL
>Q5JUK3
DETQL
>O00167
ELEYL
>Q504T8
EFVVA
>Q6P1J9
SHLRF
>D3W0D1
GTSGV
>Q9Y4C0
REYYV
>P48047
MREIV
>B4DS77
TEVTV
>Q9ULG6
HYSSL
>Q8IWV2
ARSSL
>Q9Y4C8
QTLQL
>Q9UBP0
GDTTV
>Q8NGP8
HKIAV
>Q9UBP4
GGEEI
>O94933
TTYRF
>Q7L2R6
EKLHV
>Q9UBP9
LDSRC
>O00628
LTIPA
>O00629
EGFQF
>Q8N5I4
LDVTL

>Q9UI32
LESMV
>P00558
ALSNI
>Q8TDD2
NLLEI
>Q9NX02
HDFMI
>Q5T0D9
GKIGF
>Q9NX09
LIEEC
>Q9UGL9
CCSGC
>A0A5A2
CASSL
>P61225
ACVIL
>A0A5A6
CASSL
>P13631
LKSPA
>Q92608
LSTDL
>Q92609
SPLDI
>Q6P1K1
ILSDF
>Q96TA1
VQTEF
>Q9BV99
SSFVF
>Q08257
MILLL
>P13639
FLDKL
>P48050
RESAI
>O00175
NQTTC
>Q7Z5S9
AFSKI
>O14576
VELSA
>Q86Y78
LGLML
>Q6P1K8
APSGV
>P48051
NESKV
>O14578
DQSSV
>O00178
PASGC

>Q9ULH0
RESIL
>Q13563
SNVHV
>Q13564
ATFQL
>Q86WP2
DDDDV
>Q9Y4D1
TKLNF
>Q9ULH4
MESTV
>Q0VAQ4
EEYFI
>Q6UQ28
STLLF
>A6NK89
CESLV
>Q13569
EESHA
>Q9Y4D7
CYSEA
>P01889
VSLTA
>Q8NGQ3
KISSL
>P05771
FVINV
>O94941
LRVHF
>Q9UBQ7
SELKL
>O00634
RCSAA
>Q9UQD0
RESKC
>Q6ZN84
SRLLV
>Q8N5J2
DCILL
>Q9UI40
CPVSI
>P35900
VEENI
>Q9C026
RASIA
>A0A1B0GW64
EVTCL
>Q92614
TETNA
>A0A5B6
CASSL
>Q6ZS46
AVSLF

>Q92619
QPEFV
>O00187
IISDF
>Q13571
PYSEV
>A6ZKI3
EDEDF
>Q9ULI0
FETFL
>O00189
YVIRI
>P48065
KETHL
>Q86WQ0
LGEKV
>P48066
KETHF
>Q13574
QETAV
>Q9ULI3
RRDYF
>P48067
QDSRI
>Q9ULI4
QEVDV
>P01893
VSLTA
>A6NK97
LKETI
>Q8NI99
RPLKL
>Q9Y4E6
HRFMV
>P67775
PDYFL
>A6NIR3
PDECV
>Q9NS62
EKLVI
>Q5VT06
RMLLV
>Q8NGR9
KTFFL
>Q9UBR5
KKEVL
>Q9NS67
KGIGL
>O94956
EDSRV
>Q6IEE7
IKEIA
>O75712
NLTPI

>Q96G75
KRIIF
>A8MUN3
APFCL
>Q8TDF5
NTTRV
>A6NNN8
VWEMF
>Q9NX24
LPLPL
>O75716
HTTQI
>B7ZBB8
PADAL
>Q92620
ARFGL
>P61247
VQESV
>Q92628
SRTHL
>Q96EP9
AQTSL
>Q92629
TSVCL
>Q8N271
TSLKL
>Q6P1M0
GEEKL
>B7Z8K6
KLFFL
